# Exponential-family embedding with application to cell developmental trajectories for single-cell RNA-seq data

**DOI:** 10.1101/2020.09.25.313882

**Authors:** Kevin Z. Lin, Jing Lei, Kathryn Roeder

## Abstract

Scientists often embed cells into a lower-dimensional space when studying single-cell RNA-seq data for improved downstream analyses such as developmental trajectory analyses, but the statistical properties of such non-linear embedding methods are often not well understood. In this article, we develop the *eSVD* (exponential-family SVD), a non-linear embedding method for both cells and genes jointly with respect to a random dot product model using exponential-family distributions. Our estimator uses alternating minimization, which enables us to have a computationally-efficient method, prove the identifiability conditions and consistency of our method, and provide statistically-principled procedures to tune our method. All these qualities help advance the single-cell embedding literature, and we provide extensive simulations to demonstrate that the eSVD is competitive compared to other embedding methods.

We apply the eSVD via Gaussian distributions where the standard deviations are proportional to the means to analyze a single-cell dataset of oligodendrocytes in mouse brains (Marques et al., 2016). Using the eSVD estimated embedding, we then investigate the cell developmental trajectories of the oligodendrocytes. While previous results are not able to distinguish the trajectories among the mature oligodendrocyte cell types, our diagnostics and results demonstrate there are two major developmental trajectories that diverge at mature oligodendrocytes.

## 1 Introduction

Single-cell RNA-sequencing data give scientists an unprecedented opportunity to analyze the dynamics among individual cells based on their gene expressions, but many analyses require first embedding each cell into a lower-dimensional space as an important preprocessing step in order to make downstream methods more statistically or computationally tractable. For example, these low-dimensional embeddings can be used to visualize high-dimensional data, to control for batch effects, to cluster cells into cell types, to denoise or impute single-cell data, or to estimate trajectories to understand how cells develop over time; see Sun et al. (2019) for a comprehensive overview. Typically, these embeddings are computed from an *n* × *p* gene expression matrix, where each of the *n* rows and *p* columns represent a different cell and a different gene respectively. Two of the most common methods to compute these embeddings, uniform manifold approximation and projection (UMAP, McInnes et al. (2018)) and the singular value decomposition (SVD), have different weaknesses that we strive to remedy in this work. On one hand, UMAP produces flexible, non-linear embeddings that have seen widespread usage for visualization purposes (Becht et al., 2019). However, since UMAP does not yet have proven statistical properties such as consistency, there is a lack of consensus on how to tune this method, and methods that build on these UMAP embeddings inherit this statistical ambiguity; see Cao et al. (2019) and Bergen et al. (2020) for example. On the other hand, the SVD has been extensively studied in the statistical literature, but is often restrictive in practice since it yields only linear embeddings. In this article, we advance the literature by developing the eSVD (exponential-family SVD), a non-linear embedding method that retains desirable statistical properties. As the name suggests, the eSVD is a generalization of the SVD, and embeds each cell in a non-linear fashion into a lower-dimensional space with respect to any one-parameter exponential-family distribution, allowing the researcher to have much broader modeling flexibility. Methodologically, we design the eSVD such that it can be appropriately tuned using matrix-completion ideas. Theoretically, unlike similar work that also bridges this gap between the SVD and UMAP for single-cell applications such as Durif et al. (2017) and Risso et al. (2018), we leverage recent theoretical developments in the nonconvex optimization literature that formalize the statistical properties of the eSVD. With these insights, we use the eSVD to analyze single-cell data^1^ in order to demonstrate better downstream analysis results.

To illustrate the importance of embeddings, we focus on analyzing oligodendrocytes – cells that enable rapid transmission of signals by producing myelin and providing metabolic support to neurons in the central nervous system. These cells are intriguing to study due to their constant development throughout a subject’s lifetime, unlike many other cell types that mature at adulthood (Menn et al., 2006). As mentioned in Marques et al. (2016) and Cai and Xiao (2016), understanding how oligodendrocytes develop can lead to new insights into the cause of myelin disorders such as multiple sclerosis and Alzheimer’s disease. We discuss the oligodendrocyte dataset and present a preliminary analysis in Section 2, where we provide various diagnostics demonstrating the shortcomings of the SVD. To better understand this phenomenon, we review the hierarchical model that the SVD implicitly assumes in Section 3. Specifically, suppose a hierarchical model where each cell and each gene has its own low-dimensional latent random vector. In the language of exponential-family distributions, this model assumes that the cell’s expression of a particular gene is a one-parameter exponential-family random variable whose natural parameter is the inner product of the two corresponding latent vectors. By formulating this hierarchical model, we see that the SVD implicitly assumes a Gaussian distribution with constant variance. However, this assumption is often violated since the variance of each cell’s gene expressions is observed to vary dramatically with their mean expression level (Love et al., 2014; Hicks et al., 2017). Hence, as we will review later, there is a rich line of work that extends hierarchical models of this type to analyze single-cell data by replacing the Gaussian distribution with more appropriate exponential-family distributions (Pierson and Yau, 2015; Townes et al., 2017; Durif et al., 2017; Risso et al., 2018), of which this article continues.

The aforementioned work often add additional nuances on top of the hierarchical model in order to model single-cell data better, but this often results in complicated estimators that become too intractable to statistically analyze. Hence, we design the eSVD in such a way that the posited statistical model retains the most important aspects common to the aforementioned work, while we can leverage recent theoretical developments to analyze the estimator’s statistical properties. Specifically, the eSVD uses alternating minimization, a popular and computationally efficient approach used in the matrix factorization literature to solve the nonconvex optimization problem at hand, described in Section 4. We present the eSVD’s statistical theory in Section 5, which builds upon the theoretical analyses of Zhao et al. (2015) and Lei (2018). These statistical properties include identifiability conditions and consistency, which ensure that researchers well-understand the estimated embedding and have a solid statistical foundation to build downstream analyses on top of. However, to ensure that the eSVD does not sacrifice too much modeling flexibility for theoretical tractability, we compare the eSVD to competing methods used to analyze single-cell data in Section 6 using synthetic data.

Finally, we return to our preliminary analysis of oligodendrocytes in Section 7, where we show that the eSVD embedding improves our analysis of cell developmental trajectories to match the latest scientific findings. These trajectories explain the heterogeneity among the oligodendrocytes by describing the smooth transition of gene expression among individual cells along a continuum, reflecting the cells’ gradual transcriptional changes during development (Trapnell et al., 2014). Although early research suggest oligodendrocytes develop along a single trajectory (Kessaris et al., 2006), recent work suggest that oligodendrocytes could potentially branch out into various mature types (Marques et al., 2016; van Bruggen et al., 2017; Marques et al., 2018). Our improved analysis match these findings – we show the eSVD embedding estimates two distinct trajectories. We develop visualization tools to show our developmental trajectory findings, and conclude in Section 8 with practical extensions and theoretical questions left open for future work. While we focus on using the eSVD embedding to estimate cell developmental trajectories in this article, we emphasize that this embedding can be used for other applications highlighted earlier in this section, and provide additional analyses on another single-cell dataset in the appendix.

## 2 Preliminary analysis

We analyze a dataset of oligodendrocytes from mice brains collected by Marques et al. (2016) as a prototypical example to demonstrate shortcomings of the SVD embedding when applied to single-cell data. This dataset, henceforth called the Marques dataset, contains the gene expression of 5,069 oligodendrocytes that are clustered into thirteen cell sub-types using a biclustering algorithm (Zeisel et al., 2015) in Marques et al. (2016). These thirteen cell sub-types were later grouped into six major cell types and manually labeled based on cell-type specific marker genes (Zhang et al., 2014), shown in Figure 1.. We preprocess the data by selecting 983 highly informative genes, normalizing each cell by its library size (i.e., total counts across all genes), and log_2_-transforming each entry. These details are described in Appendix C. As suggested in the literature, it is common to log_2_-transform the gene expression matrix prior to using the SVD, since the log_2_-transformation can ideally stabilize the variance (Townes et al., 2017; Butler et al., 2018). However, as we see in the preliminary analysis in this section, this analysis strategy will result in modeling concerns that we wish to remedy in the rest of this article.

**Figure 1:**
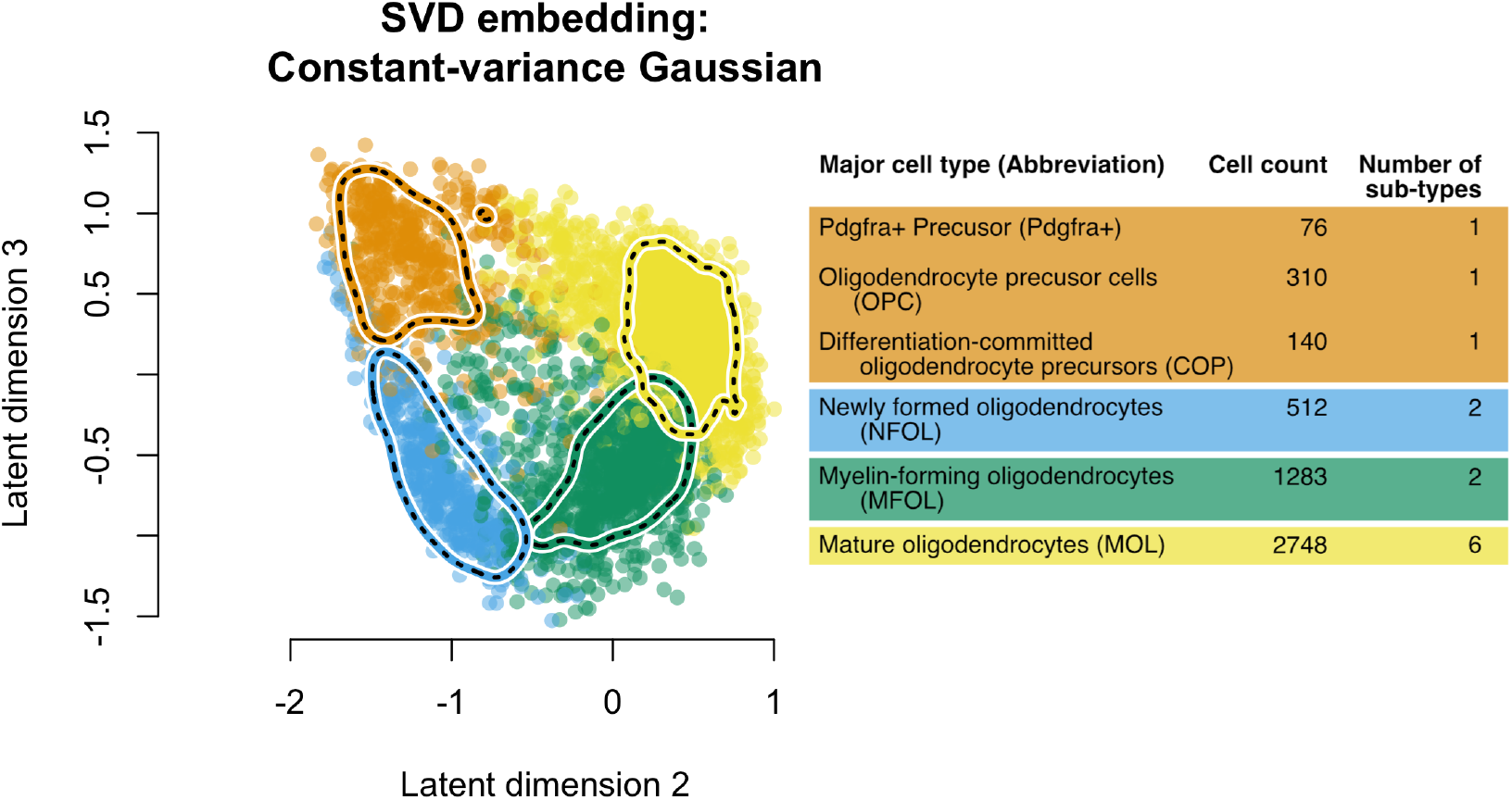
The SVD embedding of the oligodendrocytes from Marques et al. (2016) after preprocessing including a log_2_-transformation, shown alongside a table summarizing the cell types. The six major cell types are listed in the table with the number of cells in each type, along with how they are differentiated into the thirteen different cell sub-types. The rows are organized from the “youngest” cell types to “most mature” cell types from top to bottom. The youngest three major cell types are colored orange. while the oldest three are colored blue, green and yellow respectively. The second and third latent dimensions are shown on the left, along with contours of the estimated densities to visualize high-density regions (one for each color of cells).

We review the SVD embedding, as it provides motivation for the eSVD in the next section. Let *A* ∈ ℝ^*n*×*p*^ represent the observed single-cell RNA-sequencing data matrix with rank *m*, where *n* is the number of cells and *p* is the number of genes. Here, loosely speaking, each entry *A*_*ij*_ measures how many instances of genetic material for gene *j* is observed for cell *i* after pre-processing. Let the SVD of *A* be denoted as 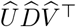 where 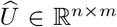 and 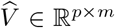 are both orthonormal matrices and 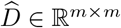 is a diagonal matrix. For a given latent dimensionality *k* ≤ *m*, the SVD embedding for each cell *i* ∈ {1,…, *n*} (denoted as 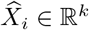) and each gene *j* ∈ {1,…, *p*} (denoted as 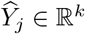) is defined as

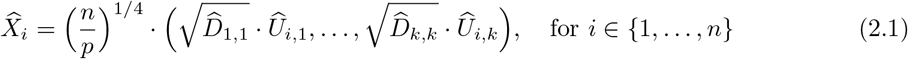

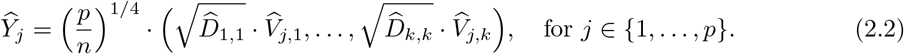

We can see that the SVD embedding is a linear embedding since 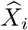 is the first *k* elements of the ith row in 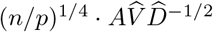. A scatterplot of the second and third latent dimensions of such an embedding is shown in Figure 1. Later in this article, we will show that this embedding implicitly assumes a constant-variance Gaussian distribution in Section 3, and show that this particular formulation handles identifiability concerns discussed in Section 4.

Now, we show that the SVD embedding (and its equivalent reparameterizations) does not model the data well, which could produce misleading results in downstream analyses. First, we visualize the quality of fit of the SVD embedding by purposefully omitting a small subset of randomly-selected entries in *A* and estimating the embedding as a matrix-completion problem. We can then assess the quality of fit of the embedding by comparing the values of these omitted entries in *A* to their predicted values. Figure 2 demonstrates this diagnostic, where the left plot shows the observed values in *A* that are not omitted (i.e., the “training set”) verses their respective predicted values, while the right plot shows the observed values that are omitted (i.e., the “testing set”) verses their respective predicted values. This missing-value diagnostic is commonly used both for assessing the quality of fit as well as for model selection (Li et al., 2020), and we will return to it in detail in Section 4. We see that while the embedding’s performance on the testing set is more-or-less equivalent to that on the training set, the variability decreases as the gene expression increases. This is in opposition with the working model in the literature that suggests that larger gene expressions should be more variable than smaller ones (Witten, 2011; Risso et al., 2018). In fact, prior to taking the log_2_-transformation, Figure 3 demonstrates that the variance in gene expression increases with its mean, reinforcing this model. Combined, all the diagnostics demonstrate that applying a log_2_-transform and using the SVD in conjunction seem to distort the properties of the data. This inspires us to develop a more appropriate embedding method that still retains desirable statistical properties. In the next section, we review the optimization problem that the SVD solves and see how it can be extended to one-parameter exponential families more generally, which will motivate our method, the eSVD.

**Figure 2:**
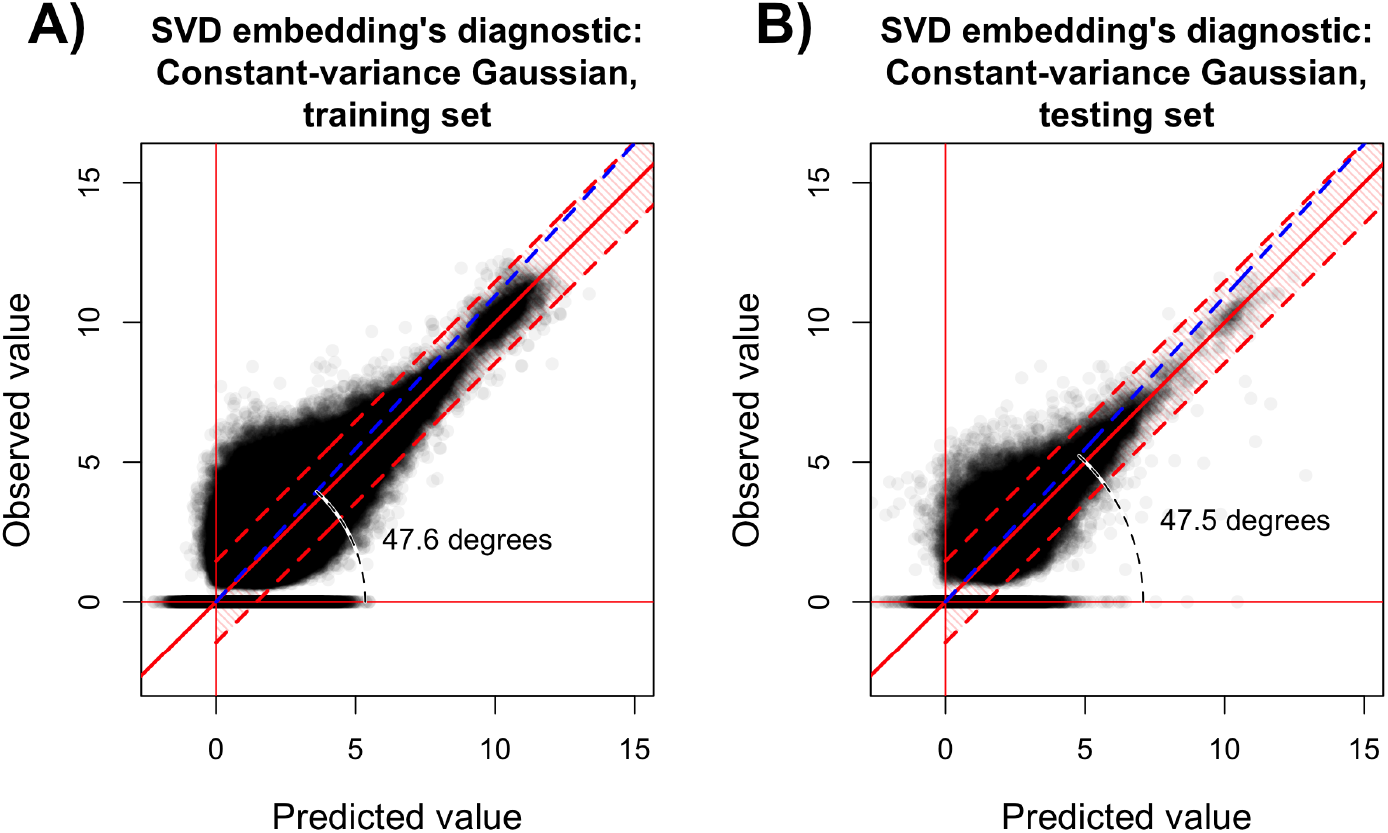
Diagnostic based on matrix completion to assess the fit using the SVD embedding for either the observed values that are not omitted (i.e., the “training set”) (A) or the observed values that are purposefully omitted (i.e., the “testing set”) (B), both verses their respective predicted values. This embedding is estimated using softImpute (Mazumder et al., 2010). The shaded red region is centered around the identity function (the ideal mean function) and marks the 10th to 90th quantiles of the constant-variance Gaussian model (based on the empirical variance) for different values of the predicted mean. The blue dotted line represents the principal angle between the observed values and their predicted value counterparts, where we mark its divergence from the identity function’s 45°. More details of this diagnostic is discussed in Section 4, while details of the fitting process using softImpute can be found in Appendix C.

**Figure 3:**
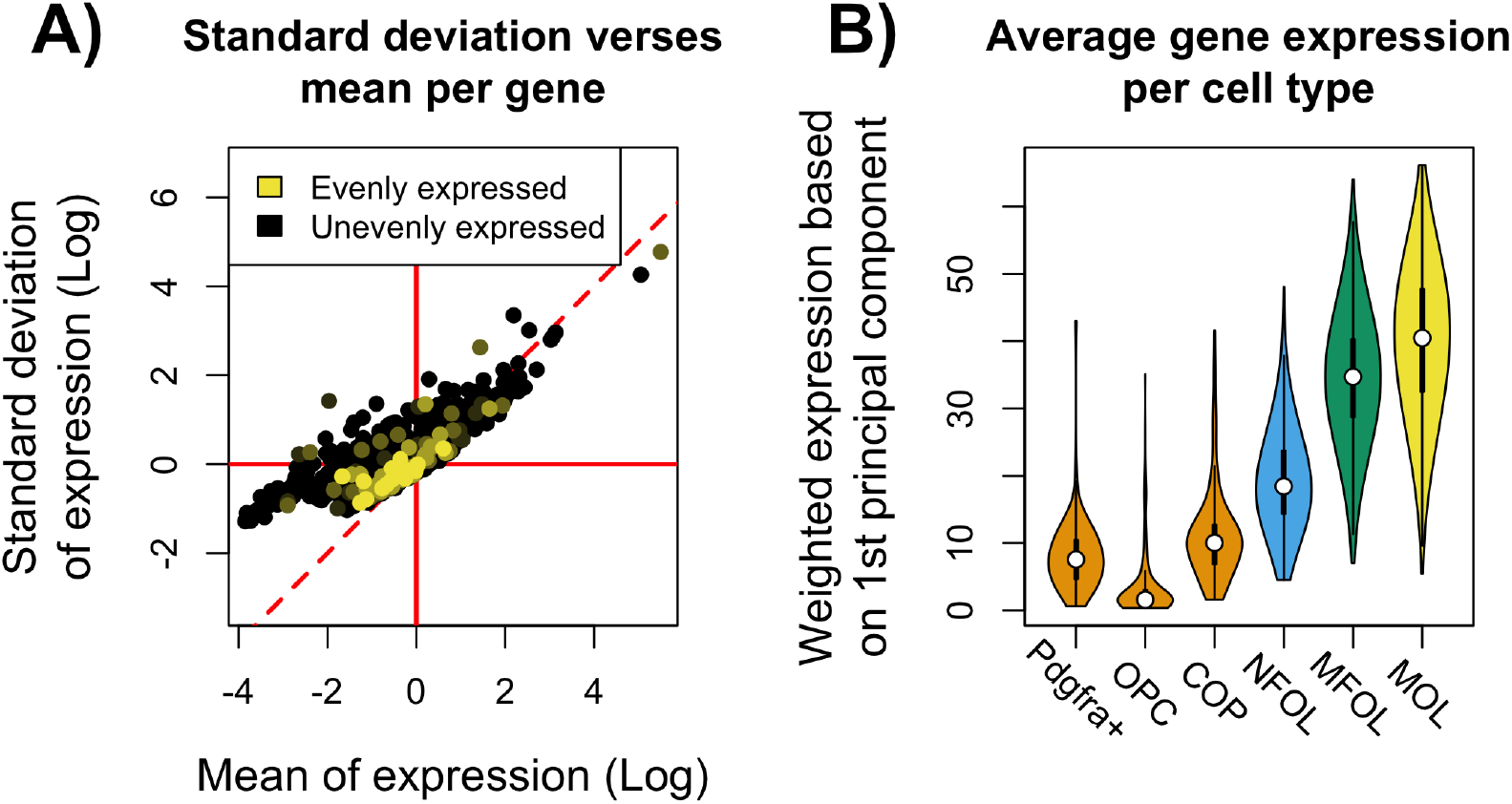
(A) The standard deviation of the expression verses the mean expression across all the cells, where each point represents one of the 983 genes in the preprocessed single-cell dataset. The color of each point depends on how evenly the gene is expressed among each of the six oligodendrocyte cell types show in Figure 1. The solid red horizontal and vertical lines and the dashed red line denoting the line *y* = *x* are for visual reference. (B) Violin plot of the average expression of the genes reweighted according to the first principal component among the six oligodendrocyte cell types, using the color scheme in Figure 1. The statistics in both plots are computed prior to taking the log_2_-transformation, and is shown on the logarithm scale in Plot A purely for visualization purposes. More details about these plots are in Appendix C.

## 3 Statistical model and background

In this section, we explain the random dot product model that we investigate in this article, and its relation to other work.

### 3.1 Statistical model and estimation strategy

We model the entries of the single-cell dataset *A* ∈ ℝ^*n*×*p*^ as conditionally independent random variables drawn from a random dot product model – a latent hierarchical model commonly used in other work (Pierson and Yau, 2015; Townes et al., 2017; Durif et al., 2017; Risso et al., 2018). Specifically, for an appropriate one-parameter exponential-family distribution *F* parameterized by its natural parameter *θ*_*ij*_, we impose model,

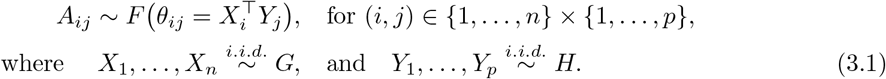

where *G* and *H* represent two latent *k*-dimensional distributions, where *k* is much smaller than *n* or *p*. We assume all the latent random vectors *X*_*i*_’s and *Y*_*j*_’s are jointly independent, and the observed *A*_*ij*_’s are independent conditioned on the *X*_*i*_’s and *Y*_*j*_’s. Let the density of the exponential-family distribution *F* be denoted as

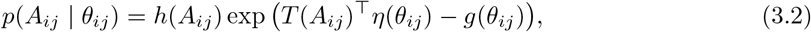

where *g*(·) is a known log-partition function for *F* with a domain 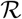, *η*(·) is a known natural parameter function, and *T*(·) is a known sufficient statistic function. For notational convenience, we denote *X* ∈ ℝ^*n*×*k*^ and *Y* ∈ ℝ^*p*×*k*^ as the matrices that collect all the latent vectors *X*_1_,…, *X*_*n*_ and *Y*_1_,…, *Y*_*p*_ row-wise, and denote 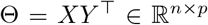 as the rank-*k* natural parameter matrix that collects all elements *θ*_*ij*_. Given the exponential-family form for *F* shown in (3.2), we need to impose the following assumption to ensure the inner products 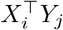 for all (*i*, *j*) ∈ {1,…,*n*} × {1,…,*p*} yield valid natural parameters,

#### Assumption 3.1

(Bounded inner product). *Let* 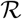 *denote the domain of the natural parameters for the distribution *F*. Assume that for any X*_*i*_ ∼ *G and Y*_*j*_ ∼ *H*,

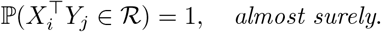

Given the above model, our goal is to estimate the latent random vectors *X*_1_,…, *X*_*n*_ since these latent vectors represent the low-dimensional embedding of all *n* cells. For a given exponential-family distribution *F*, an intuitive strategy for estimating our desired embedding is to minimize the negative log-likelihood based on the observed data *A* over all possible vectors *X*_1_,…, *X*_*n*_ and *Y*_1_,…, *Y*_*p*_. Specifically, plugging into the exponential-family form (3.2) into the model (3.1), we derive the loss function

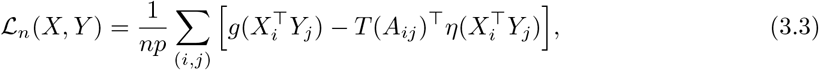

with the constraints 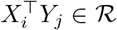 for all pairs (*i*, *j*). The above loss function is nonconvex, but if *F* is the the constant-variance Gaussian distribution, this loss function is proportional to

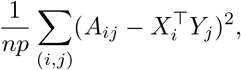

which is specifically what the SVD minimizes (Maezika, 2016). This particular model is convenient to use since the SVD provides a closed-form solution to the corresponding nonconvex optimization problem, and leads to the SVD embedding shown in (2.1).

### 3.2 Relation to other work modeling single-cell data

As we have discussed previously in Section 2, this constant-variance assumption is too restrictive to properly model single-cell data. Hence, many articles cited above replace the constant-variance Gaussian distribution with other exponential-family distributions for *F* that allow the variance to increase with the mean. For example, Witten (2011) and Risso et al. (2018) consider the Poisson and negative binomial distribution specifically, after a suitable transformation of the natural parameters. Additionally, these models often add other random effects on top of the existing random dot product model (3.1) that influence the entries in *A*. For example, many methods like pCMF (Durif et al., 2017) allow researchers to incorporate *dropout* into the model – a characteristic of single-cell data where a substantial fraction of the gene expression for a cell is recorded as exactly 0 due to low amounts of RNA in the cell (Kharchenko et al., 2014). Other methods such as ZINB-WaVE (Risso et al., 2018) go further and allow covariate information such as gene length and cell size. Most recently, Lopez et al. (2018) use deep autoencoders to estimate the embedding. However, there is often a lack of theoretical analyses for the aforementioned estimators. This is because replacing the exponential-family distribution *F* with any distribution aside from the constant-variance Gaussian distribution leads to non-trivial nonconvex estimators that minimize the loss function shown in (3.3), which make traditional statistical techniques for analyzing these estimators unsuitable. Therefore, concerns such as identifiability are typically not addressed theoretically, leading to ambiguity on performance of the estimators of such models.

To resolve this theoretical ambiguity, we design the eSVD to estimate the embedding based on the random dot product model (3.1) for any exponential-family distribution *F* with no other random effects. In this way, we can tractably analyze the statistical properties of eSVD while retaining abundant flexibility to effectively model single-cell data.

### 3.3 Matrix factorization

To the best of our knowledge, the first statistical results for estimators that extended the SVD to generic exponential-family distributions by minimizing a loss function similar to (3.3) come from Gunasekar et al. (2014) and Lafond (2015). There, the authors minimize the loss function over the natural parameter matrix Θ and add a trace penalization term to encourage the estimate to be low-rank. While this formulation yields a convex optimization problem, it requires solving a semidefinite program which can be computationally prohibitive for large datasets. This consideration has motivated researchers to investigate the statistical properties of estimators that minimize the loss function (3.3) directly as a nonconvex optimization problem. Specifically, *alternating minimization* is a suitable candidate for this task, where each iteration alternates between optimizing either one of two low-rank matrices *X* and *Y* while treating the other fixed. This algorithmic strategy pre-dates the convex relaxation approach; see Collins et al. (2002), Jain et al. (2013), Udell et al. (2016), Landgraf and Lee (2019) and the references within for discussions and additional variants. From an algorithmic standpoint, our method is a direct continuation of such work. However, the statistical properties of such estimators have only recently been characterized rigorously. For example, to accommodate the constraints in Assumption 3.1, Wang et al. (2016), Yu et al. (2020) and Chi et al. (2019) adapt the theoretical framework to study slightly different estimators based on alternating projected gradient descent. In contrast, our work will build on techniques used in Zhao et al. (2015) and Balakrishnan et al. (2017) to retain our focus on alternating minimization.

However, all the aforementioned theoretical results do not directly apply the random dot product model (3.1), which contains an additional source of randomness induced by the hierarchical structure. In contrast, our theory is able to account for this additional source of randomness by drawing upon connections to the network literature. Specifically, the random dot product model (3.1) is similar to those used in latent position random graphs studied in the network literature (Hoff et al. (2002) and Athreya et al. (2017)). Hence, we draw inspiration from Lei (2018) on how to address these identifiability concerns and to develop proof techniques in this article.

There are many other embedding methods in the literature more broadly, such as non-negative matrix factorization, kernel PCA, and manifold-based embedding methods such as UMAP and Isomap. We defer a thorough discussion contrasting eSVD with such estimators to Appendix B.

## 4 Method: eSVD (Exponential-family SVD)

We describe the eSVD in this section, which is designed to be a general framework to minimize the loss function (3.3) for any choice of a one-parameter exponential-family distribution *F*. To keep the presentation clear, we describe some of the more nuanced implementation details in Appendix B. We also describe an important diagnostic to assess the quality of fit, as demonstrated in Section 2. This diagnostic can be also be used as a tuning procedure to select the most appropriate choice for *F* or nuisance parameters.

### 4.1 eSVD and its application to the curved Gaussian distribution

Similar to other nonconvex matrix factorization methods (Wang et al., 2016; Yu et al., 2020), our method requires an initial estimate of the rank-*k* matrix of natural parameters, 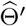, where *k* is pre-determined. To achieve this, we use an initialization method based on Wang et al. (2016). To simplify the presentation here, we provide details in Appendix B.1. This initialization scheme performs a rank-*k* SVD based on transforming each entry of *A* via the inverse of the log-partition function *g*(·). Given this initial estimate, consider its 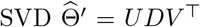. To start the alternating minimization stage of our method, we set 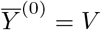.

After initialization, the eSVD then refines the estimate by performing alternating minimizations. Denoting a generic matrix and its SVD by 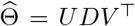, let 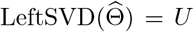, the function that maps a matrix to its left singular vectors. Then, for iterations *t* ∈ {0,…, *T* − 1},

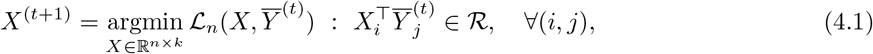

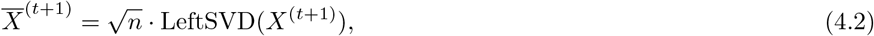

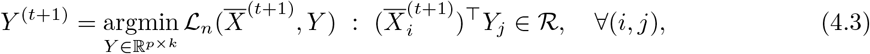

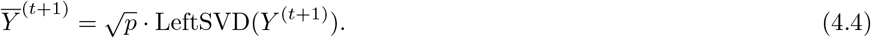

After all *T* iterations, the eSVD outputs the final estimate after a reparameterization. That is, Letting 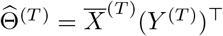 have a rank-*k* SVD of 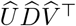, the final estimates are

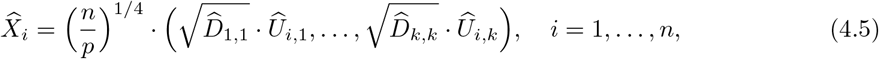

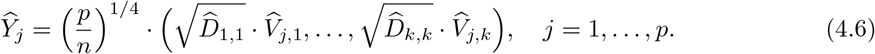

This is the same reparameterization used in (2.1) and (2.2).

#### Remarks about algorithmic design

We make a few remarks about the design of our algorithm. Optimizing over *X* and *Y* directly raises identifiability issues, since for any orthogonal matrix *Q*, 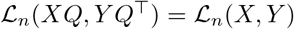. To address this, Ge et al. (2017) append a penalty term

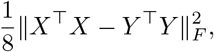

while Zhao et al. (2015) use the QR-decomposition between iterations, and we use the LeftSVD(·) operator. In practice, we found all three choices behave similarly. The factors 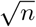 and 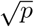 in (4.2) and (4.4) are included for theoretical reasons to ensure the spectrum of the Hessian is well-controlled and to ensure the values do not underflow if *n* or *p* are too large empirically. Also, the final reparameterizations in (4.5) and (4.6) are designed such that the sample second-moment matrices of 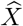 and 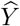 are both equal and diagonal, i.e.,

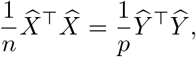

which is important for our statistical analysis later on.

Lastly, to perform the constrained optimization (4.1) and (4.3), we use a first-order method called Frank-Wolfe (Jaggi, 2013), which we found more stable compared to using projected gradient descent. While there are theoretical guidelines for choosing step-sizes related to the convexity and smoothness for this method, we found these choices often led to poor empirical performance.

#### Example with the curved Gaussian distribution

To make the eSVD’s workings more concrete, we demonstrate what the minimization in (4.1) would entail when we set *F* to be the *curved Gaussian* distribution. This will be useful later in this article when we use this distribution to analyze the oligodendrocytes. Specifically, we say the random variable *A*_*ij*_ follows a curved Gaussian distribution with a known parameter τ > 0 if 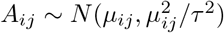, for an unknown mean parameter *μ*_*ij*_ > 0 (Efron et al., 1978; Liu and Martin, 2020)^2^. This sets the standard deviation to be linearly proportional to the mean. Writing this distribution in exponential-family form (3.2) yields,

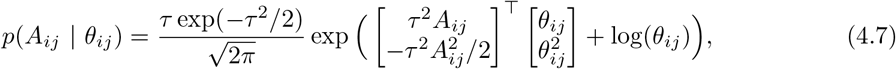

where the relation between the natural parameter *θ*_*ij*_ and the mean parameter *μ*_*ij*_ can be derived to be *μ*_*ij*_ = 1/*θ*_*ij*_. Here, the domain of the natural parameters would be 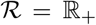, the positive half-line. After simple calculations, one can derive the negative log-likelihood and conclude that the minimization in (4.1) becomes

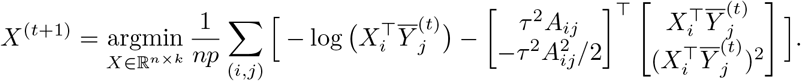

Analogous calculations for other common distributions are shown in Appendix B.

The curved Gaussian distribution is relevant in practice since if τ ≥ 2, this distribution reflects the phenomenon that genes with larger expression also exhibit larger variance, while most of the distribution’s mass is still positive. Additionally, in many instances, this distribution can capture more variability than the Poisson and negative binomial distributions, which can be beneficial when the single-cell data is intrinsically noisier. In general however, if the researcher wants to use the eSVD for an arbitrary one-parameter exponential-family distribution *F*, all she needs to pass into our implementation is the computation of the loss function (3.3), its gradients, and information about the domain 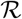.

### 4.2 Matrix-completion diagnostic and tuning procedure

We provide the following diagnostic to assess the embedding’s quality of fit or to determine which choice of *F* is most appropriate for our data, which was used in Figure 2. Inspired by network cross-validation work such as Li et al. (2020), we use matrix completion to determine the quality of our model fit. As alluded to in Section 2, to do this, we omit a small percentage of the entries of *A* when estimating the embedding and compare these values to their predicted expected value counterparts. To compute this expected value, recall that for exponential-family distributions (3.2), *g*′(·) (the derivative of the log-partition function *g*(·)) maps the natural parameter to the expected value. We note that we are able to adopt this matrix completion strategy to tune our method since our alternating minimization procedure can be adapted to handle missing values. In contrast, embedding methods like ZINB-WaVE (Risso et al., 2018) and PCMF (Durif et al., 2017) do not offer an analogous tuning procedure. This procedure is formalized below.

1. For bootstrap trials *b* ∈ {1,…, *B*}:

a. Randomly sample *m* of the entries of *A*, denoted as 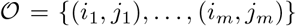, which will be omitted in the following estimation step. Here, *m* can be any small number, such as [0.01 · (*np*)].
b. Estimate the latent vectors by 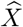 and 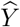 according to Subsection 4.1 where the loss function (3.3) omits the entries in 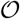 and is parameterized based on the desired distribution of *F*.
c. Compute *v*_1_, defined as the leading eigenvector of the matrix formed by the omitted observed values 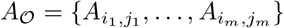 and their predicted expected value counterparts 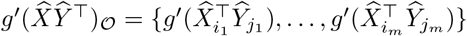.
d. Compute model fit quality, *q*^(*b*)^ defined as the angle between *v*_1_ and the vector (1, 1), representing the identity function.
2. Average the model fit qualities across all trials, *q*^(1)^,…, *q*^(*B*)^.

Observe that we define the quality of fit *q*^(*b*)^ above by how much the leading eigenvector *v*_1_ deviates from 45°. This angle of *v*_1_ is what we called the *principal angle* in Figure 2. Having an eigenvector’s angle close to 45° means that on average, the predicted values correspond closely with the observed value. The advantage of this quality-of-fit’s definition is that it is easily comparable even across different distributions *F*, unlike the negative log-likelihood or MSE.

While we advocate constructing plots such as Figure 2 to obtain a more holistic sense of how well the embedding fits the data in general, we can also use the above procedure to obtain an automated model selection method in the following way – if we try the above diagnostic for multiple distributions for *F*, the distribution that yields the smallest average of *q*^(1)^,…, *q*^(*B*)^ is deemed the most appropriate model for *A*. In this way, we can also use this diagnostic as a tuning procedure to select the dimensionality of the latent space *k* or nuisance parameters for exponential-family distributions such as *τ* in the curved Gaussian distribution (4.7) in a grid-search fashion. An additional variant of this tuning procedure is detailed in Appendix B.

## 5 Statistical theory

In this section, we prove the consistency for the eSVD when applied to the random dot product model (3.1), which is important to ensure that the eSVD is estimating a well-defined quantity. This result provides the needed statistical foundation for downstream tasks such as clustering and RNA velocity, as mentioned in Section 1. Additionally, our theory gives us better insights into the eSVD since our analysis also reveals the identifiability conditions that formalize to what degree the embedding can be estimated. While our theorems currently assume the correctly-specified setting (i.e., the eSVD’s choice of *F* matches the true generating distribution and *k* is correctly-specified), we hope these theorems provide a roadmap for future work to prove analogous statements for broader settings or for more complex methods like ZINB-WaVE which currently do not exist.

We discuss the additional notation here. For a generic matrix *A*, let ||*A*||_*F*_ denote its Frobenius norm. For two sequences *a*_*n*_ and *b*_*n*_ and two random sequences *A*_*n*_ and *B*_*n*_, let *a*_*n*_ = *O*(*b*_*n*_) and *A*_*n*_ = *O*_*P*_(*B*_*n*_) denote that *a*_*n*_/*b*_*n*_ or *A*_*n*_/*B*_*n*_ is bounded for large enough *n* deterministically or in probability respectively. For all *i* ∈ {1,…, *n*} and *j* ∈ {1,…, *p*}, let the the population second moment matrices of *X*_*i*_ and *Y*_*j*_ and the corresponding eigen-decompositions be defined respectively as

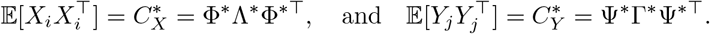

Our proposition below requires the following assumptions.

### Assumption 5.1

(Sub-Gaussian distribution of latent vectors). *Assume that X*_*i*_ *for all i* ∈ {1,…, *n*} *are i.i.d. sub-Gaussian random vectors. That is, there exists a fixed constant D such that for any vector v* ∈ ℝ^*k*^ *where* ||*v*||_2_ = 1 *and any integer c* ≥ 1, 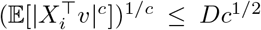, *and a similar assumption holds for Y*_*j*_ *for all j* ∈ {1,…, *p*} *with also the same constant D*.

### Assumption 5.2

(Second moment properties). *First, assume the population second moment matrices* 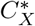 *and* 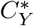 *are equal and are both diagonal matrices, where* 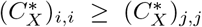 *for any* 1 ≤ *i* < *j* ≤ *k. Second, assume there exists positive numbers c*_1_ ≤ *c*_2_ *and* 1 < *α* ≤ *β such that for all* ℓ ∈ {1,…, *k*}, *the eigenvalues satisfy*

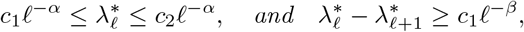

*with the convention that* 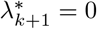.

Both assumptions are common in work that study the spectrum associated with random dot product models (Lei, 2018). Assumption 5.1 assumes *G* and *H* are sub-Gaussian distributions, which enables sharp rates for estimating their second-moment matrices 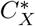 and 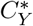 respectively (Vershynin, 2012). On the other hand, the second part of Assumption 5.2 enables our estimator to accurately estimate its eigenvalues and eigenvectors. Importantly however, the first part of Assumption 5.2 can be interpreted instead as an identifiability condition, which we formalize below.

### Proposition 5.1.

*Given two k-dimensional distributions G and H, each with at least two moments where the population second moment matrices are full rank, consider two independent random variables X′ ~ G and Y′ ~ H. Then there exists a linear and invertible transformation R such that the population second moment matrices of X* = *RX′ and* 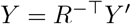 *are the same, i.e.*,

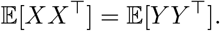

*Furthermore, both population second moment matrices of X and Y are diagonal matrices*.

The proof is in Appendix I, which provides an explicit construction of the matrix *R*. Note that since *R* is invertible, we guarantee that the distribution of the inner product is preserved, i.e.,

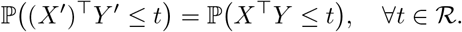

Hence, the first part of Assumption 5.2 can be interpreted as an identifiability condition, since we can only estimate *G* and *H* only up to a linear transformation.

### Proposition 5.2.

*Assume the model in* (3.1) *where Assumptions 5.1 and 5.2 hold. If the estimator* 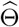 *satisfies* 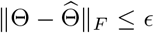 *conditioned on X and Y, and k* = *o*(min {*n, p*}), *then up to sign*,^3^ *eSVD achieves the rate after reparameterizations* (4.5) *and* (4.6),

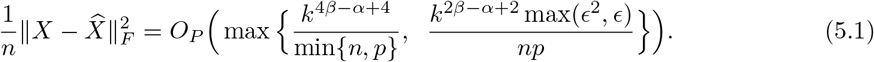

### Discussion of consistency

Assuming *k* is fixed, the above proposition states that the eSVD embedding is consistent as long as ɛ(the rate of convergence for the matrix of natural parameters, 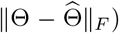 is faster than 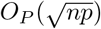. We formalize this statement in Appendix D, where we add assumptions common to the literature such as strong convexity and smoothness of the negative log-likelihood function associated with *F*. The details are deferred because these assumptions are technical to describe and detract from the main text. More generally, however, the above proposition addresses the additional source of randomness induced by the random dot product model mentioned in Section 3 that other theoretical investigations typically do not address. Therefore, *any* such estimator equipped with a rate for 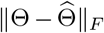 can be plugged into Proposition 5.2 directly. Additionally, note that a similar rate (5.1) holds for estimating *Y*, the embedding of the genes.

### Application to curved Gaussian model

While Proposition 5.2 and the results in Appendix D apply for any generic one-parameter exponential-family distribution *F* satisfying certain conditions, we specifically apply these results to the curved Gaussian distribution (4.7) in Appendix E to demonstrate what the rates are for a particular exponential-family distribution. At a high level, we show that when *n* and *p* grow asymptotically at the same rate and *k* is fixed,

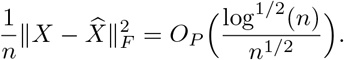

## 6 Numerical study

In this section, we study the performance of the eSVD and other competitive methods based on synthetic data. Our setup for all the simulations in this section are as follows: based on model (3.1), we set the dimensionality to *k* = 2 and sample *X*_1_,…, *X*_*n*_ i.i.d. uniformly from four connected linear segments (which we call the “trajectories”) with additive Gaussian noise, as illustrated in Figure 4. These four segments loosely represent four cell types. We also sample *Y*_1_,…, *Y*_*p*_ i.i.d. from a mixture of two Gaussians. These sampling procedures represent *G* and *H* respectively, up to identifiability conditions. We enforce 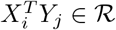 for all pairs (*i*, *j*). The distribution family *F* varies among different simulations. We do not use `R` packages such as Splatter (Zappia et al., 2017) to generate our synthetic data because we want to have precise control over the true embedding. The full details of the simulation setups and usage of various estimators in this section are in Appendix F.

**Figure 4:**
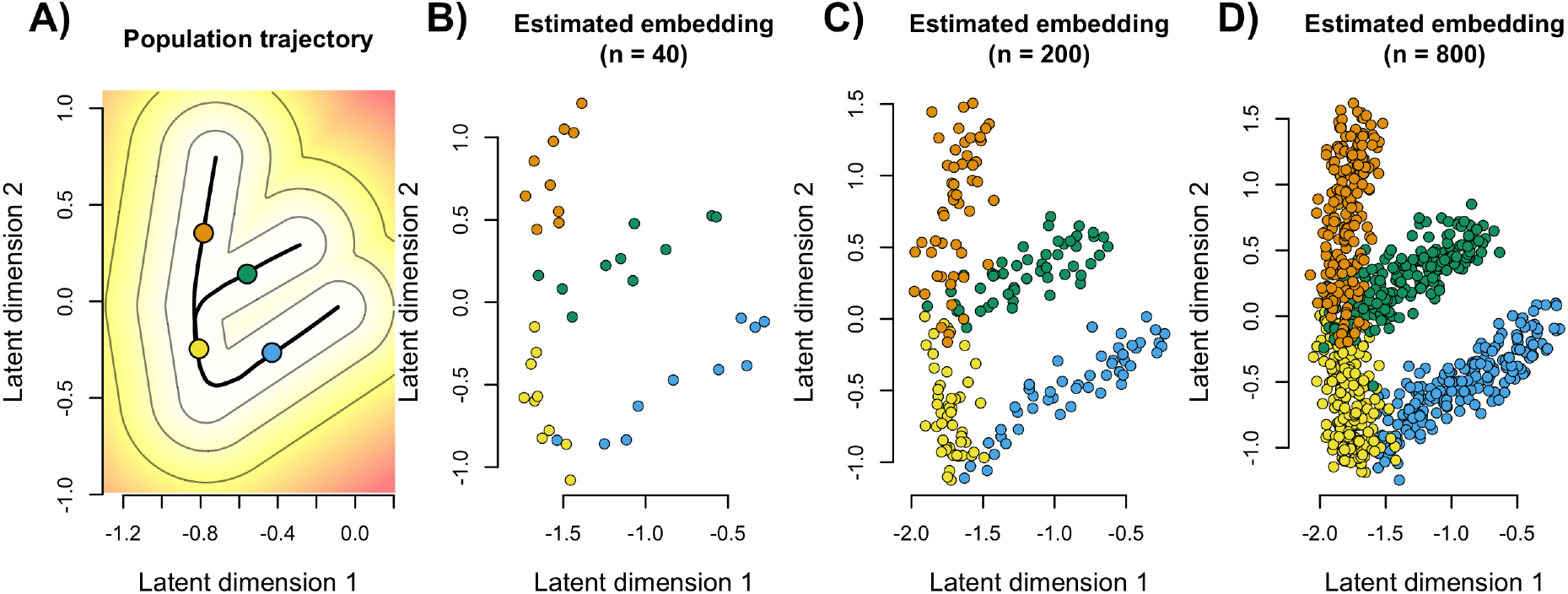
(A) The two-dimensional population density of *G*, visualized as a heat map with contour lines of the density along with the true “trajectories” (black lines). The mean vector for each of four cell types are labeled in a different color (blue, yellow, green, orange). (B to D) The estimated embedding 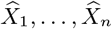 of the synthetically-generated *A* for varying levels of *n*, (i.e., number of cells or number of rows), colored by the true cell type, which are labels used only for visual reference and not used during estimation.

### Consistency of the estimated embedding

In this first simulation suite, we demonstrate that the estimated embedding converges towards the true embedding. Specifically, we generate *A* ∈ ℝ^*n*×*p*^ where each entry *A*_*ij*_ is sampled independently from the negative binomial distribution with a natural parameter 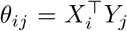 and *r* = 50, and fit the eSVD using the correctly specified model. Figure 4 is an illustration that demonstrates the asymptotic properties of the eSVD. Specifically, we see that the distribution of the embedding 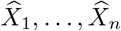 approximates *G* as *n* increases. We provide details and additional results that verify the consistency of the eSVD’s embedding in Appendix F.

### Comparison of different embedding methods

In our second simulation suite, we demonstrate that the cells’ latent positions estimated by the eSVD are more accurate in relation to one another than those estimated by other methods. Here, we fix *n* = 200 and *p* = 400. We compare the eSVD via the negative binomial distribution to nine other methods commonly used to embed single-cell data: ZINB-WaVE (Risso et al., 2018), pCMF (Durif et al., 2017), SVD, non-negative matrix factorization (NMF), independent component analysis (ICA), UMAP (Becht et al., 2019), t-SNE (Maaten and Hinton, 2008), Isomap (Tenenbaum et al., 2000) and diffusion map (Haghverdi et al., 2015). The first two methods and our tuning procedures are explained in Appendix F. Importantly, SVD implicitly assumes *F* is a constant-variance Gaussian distribution as mentioned in Section 3, while ZINB-WaVE and pCMF assume *F* is a negative binomial and Poisson distribution respectively.

We simulate data from a negative binomial model in this simulation, which is the distribution that is most commonly used to model sequencing data (Love et al., 2014). Specifically, we sample the observed count matrix *A* conditionally independent on *X*_1_,…, *X*_*n*_ and *Y*_1_,…, *Y*_*p*_ where *A*_*ij*_’s are sampled from a negative binomial distribution with natural parameter 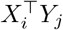 and dispersion parameter *r* = 50. Then, when we estimate the embedding using eSVD, we use the tuning procedure mentioned in Section 4 to select the most appropriate value of the dispersion parameter *r* from the set {5, 50, 100}.

We find that on average across 100 trials, our method estimates the relative latent positions of each cell to be more accurate than other methods (Figure 5A). To define our notion of accuracy, consider each cell *i* and its Euclidean distance to all other *n* − 1 cells in the latent space in both the true and estimated embedding. We then compute the Kendall’s tau correlation between these two vectors, which only relies on the ranks of the distances, and then average this value over all *n* cells. Hence, a high averaged Kendall’s tau value suggests the latent positions of the *n* cells are well-estimated with respect to one another. We call this notion of accuracy the *relative embedding correlation*. We define our notion of accuracy in this way to ensure it is insensitive to arbitrary rotations or constant rescalings of the embedding. Figure 5B compares the different estimated embeddings to the true embedding as an illustration. Both the eSVD and ZINB-WaVE estimate embeddings where the four cell types are relatively in the correct configuration, and their accuracy are quite high. For the other methods, the high variability due to the overdispersion of the negative binomial seems to dramatically skew the embedding for certain cells. We defer additional simulations using other distributions *F* to Appendix F.

**Figure 5:**
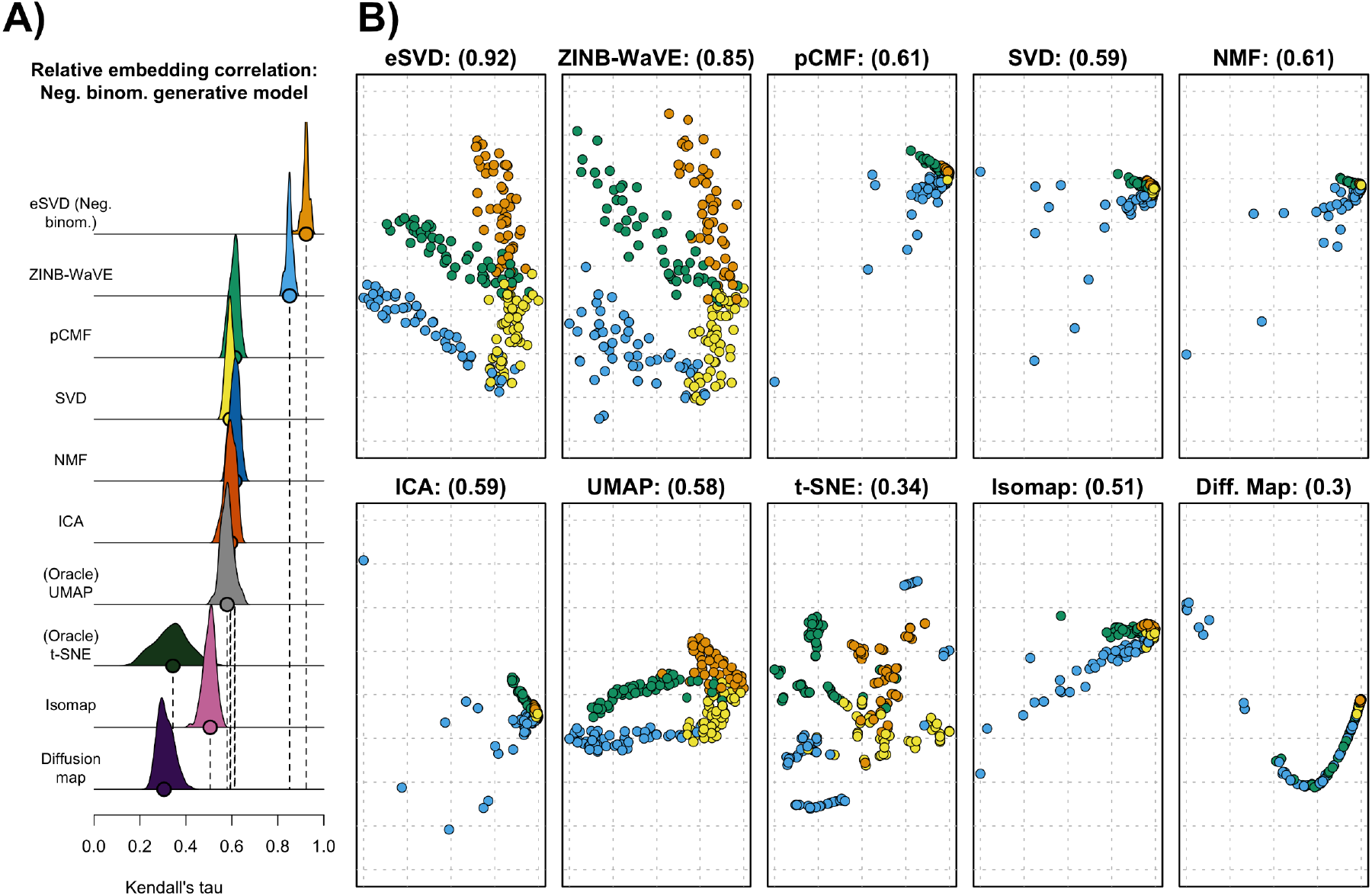
(A) The density plot of each embedding methods’ accuracy (eSVD, ZINB-WaVE, pCMF, SVD, NMF, ICA, UMAP, t-SNE, Isomap and diffusion map), based on the relative embedding correlation. The circles along each method’s x-axis denotes the median accuracy across the 100 trials. Here, both the data-generating distribution *F* and the eSVD use the negative binomial distribution. See Appendix F to see how we tuned the methods. (B) The ten estimated embedding, chosen among the trial with the median accuracy, noted in each plot’s title in parenthesis. The coloring of the samples persists from Figure 4. The x- and y-axes represent the coordinate system estimated by the respective embedding methods, reflected in the dashed grids.

### Investigation using misspecified models and the effect of *k*

In our third simulation suite, we empirically compare the eSVD to other methods when *F* is *misspecified*. We also demonstrate how different values of *k* can affect the quality of the embedding. While the theorems we developed in Section 5 do not currently handle such settings, these results help us understand how the eSVD performs in more realistic and more challenging scenarios. Due to space constraints, we defer these results to Appendix F, which show that the eSVD can roughly estimate the relative positions of the cells’ embedding well compared to other methods despite the model misspecification. All-in-all, our takeaway message is that the eSVD’s flexibility in choosing which exponential-family distribution *F* to use and the diagnostics provided in Section 4 allow our method to remain competitive among the ten methods.

## 7 Single-cell analysis

We return to modeling the Marques dataset (Marques et al., 2016), as described in Section 2, to determine if the embedding based on the curved Gaussian distribution (4.7) is more appropriate than that based on the constant-variance Gaussian distribution, and if so, investigate how the embedding affects the downstream trajectory analysis. As alluded to in Section 2, the six major cell types in Figure 1 have a determined ordering, starting from Pdgfra+ precursors and ending with the mature oligodendrocytes. Our goal in this analysis is to estimate the trajectories among the cell sub-types constrained to this ordering. For example, in Marques et al. (2016), after embedding the cells into a latent space, the authors estimate one developmental trajectory connecting the first five major cell types starting from the Pdgfra+ precursors, but do not definitively conclude how the six mature oligodendrocyte cell sub-types differentiate. Instead, they relied on analyzing the percentage of different cell types across different brain regions to hypothesize that these six sub-types differentiate into multiple different trajectories.

### 7.1 Details of estimating cell developmental trajectories

We provide more details on how we estimate the developmental trajectories based on the low-dimensional embedding 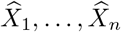. As alluded in Section 1, these trajectories show how these different cell sub-types develop from one to another, assuming the latent vectors 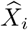 gradually change along the trajectories. Trajectory analyses are an important step in studying the cellular dynamics from single-cell data, as most single-cell technologies provide only a snapshot of all the cells. This is because most technologies destroy the cells during the sequencing step, which prevent longitudinal studies. In this article, we use Slingshot (Street et al., 2018) (with minor modifications) to estimate these cell developmental trajectories. Roughly speaking, Slingshot is a two-step algorithm that requires the latent vectors to already be clustered, where we treat each cell sub-type as a cluster. In the first stage, Slingshot estimates the number of trajectories and ordering of the cell sub-types based on minimizing the distances between cell sub-type centers via a minimum spanning tree. In the second stage, Slingshot fits variants of principal curves (Hastie and Stuetzle, 1989) that pass through the cell sub-type centers in the estimated ordering. These principal curves can be thought of as smooth curves that pass through high-density regions in the latent space. Throughout our analysis in this section, we apply Slingshot to the embedding using all latent dimensions, but only visualize the estimated trajectories with respect to the first three latent dimensions. More details about Slingshot and our modifications of it are given in Appendix G.

We briefly mention that the original study (Marques et al., 2016) uses the Monocle algorithm (Trapnell et al., 2014), to estimate the cell developmental trajectories. We use Slingshot instead as it is the current state-of-the-art method based on extensive benchmarking comparisons in Saelens et al. (2019).

### 7.2 Analysis using the constant-variance Gaussian distribution

Building on the analysis in Section 2, we perform a trajectory analysis using the SVD embedding shown in Figure 1 on the log_2_-transformed data, which assumes the constant-variance Gaussian model. Applying Slingshot directly to this embedding results in two trajectories, both heavily overlapping one another when visualized (Figure 6A). These results are similar to Marques et al. (2016) in two ways. First, the authors show that all cells develop from Pdgfra+ precursors to myelin-forming oligodendrocytes in the same way, which we estimate as well. Second, the authors do not definitively conclude if the mature oligodendrocytes diverge in their development. Our trajectories themselves also leave this ambiguity unresolved due to the heavy overlap between the two trajectories. However, we perform the following additional visual diagnostic to quantify if these two trajectories are well approximated by a single trajectory.

**Figure 6:**
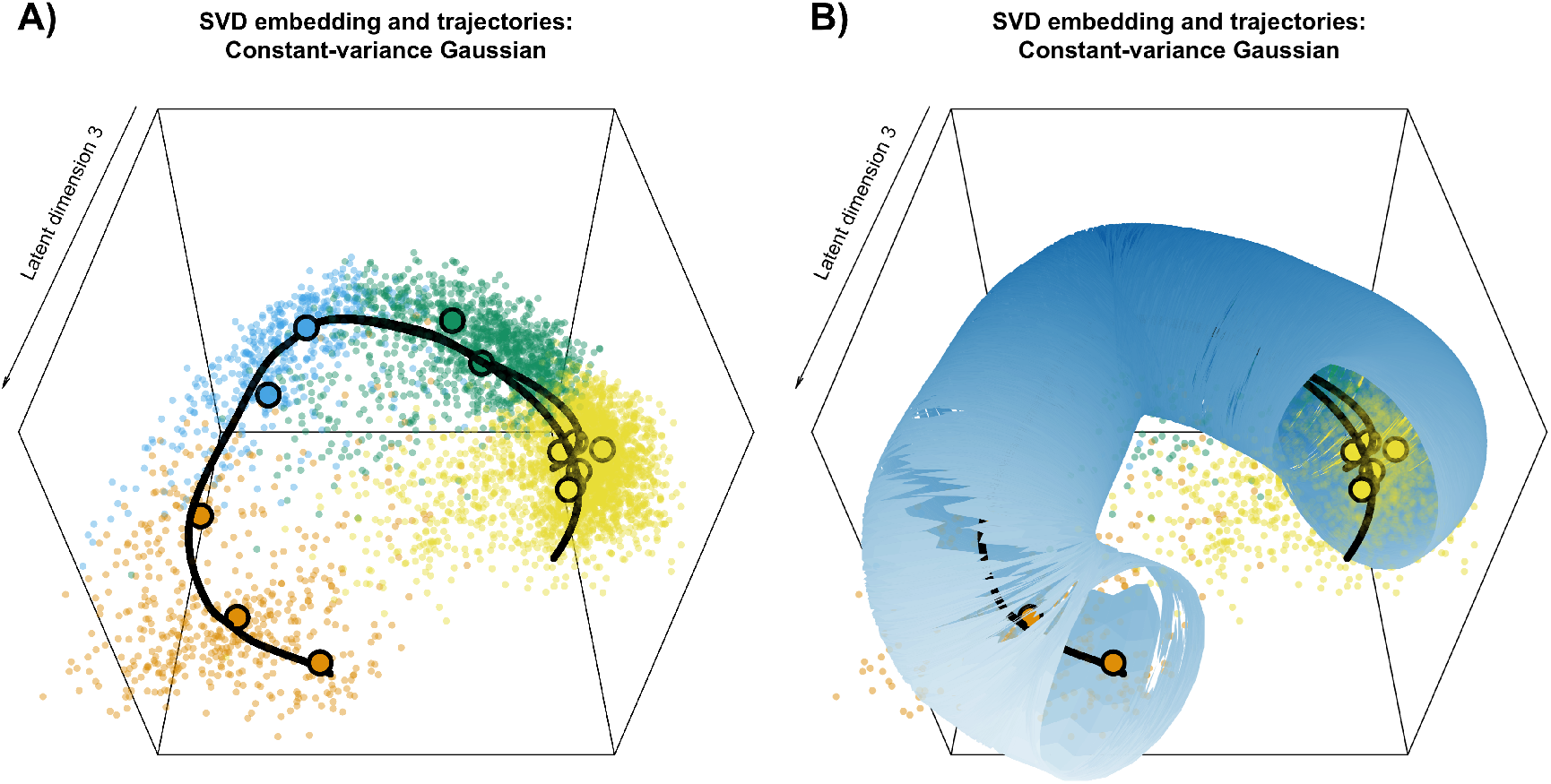
(A) Three-dimensional plot of the estimated latent positions via the SVD embedding with the two estimated cell developmental trajectories laid on top, corresponding to the data shown in the Figure 1. The thirteen bolded points correspond to the cluster centers of the thirteen cell sub-types, where the color scheme persists from Figure 1. (B) The uncertainty tube overlaid on top of Figure A.

To formalize to what degree the different trajectories are the same, we use a bootstrap resampling procedure to construct a uniform uncertainty tube around each trajectory. These tubes capture the variance of each estimated trajectory, and plotting these tubes is a useful descriptive tool. This is an important tool for our analysis because Slingshot is sensitive to small perturbations in the data due to its graph-based strategy to estimate the ordering of the cell sub-types. Specifically, small variations can dramatically change the number of estimated trajectories or ordering of cell sub-types within those trajectories. Hence, our procedure to construct these uniform uncertainty tubes first samples with replacement from all embedded cells within each of the thirteen cell sub-types. For each bootstrap sample, we apply Slingshot to estimate a new set of trajectories. We then compute the ℓ_2_ distance between the new trajectories and the original trajectories. After applying this procedure multiple times, the 95% quantile of the ℓ_2_ distances determines the uniform radius of the uncertainty tube, centered around the original trajectory. More details of this procedure are in Appendix G. Based on this construction, both trajectories lie in a single uncertainty tube (Figure 6B); hence, we conclude there is effectively one trajectory that connects all thirteen cell sub-types. This result explains why previous work such as Marques et al. (2016) had difficulty explaining how mature oligodendrocytes differentiate in their trajectory analysis.

### 7.3 Analysis using the curved Gaussian model

The above conclusions, however, rest on the questionable constant-variance Gaussian distributional assumption (see Figure 3). As we have seen in Figure 2, our matrix-completion diagnostic suggests that this assumption is not suitable for modeling the oligodendrocyte dataset at hand.

This finding motivates us to analyze the data using the eSVD to embed each cell with respect to the curved Gaussian distribution (4.7), and to re-examine the resulting diagnostics. Following suggestions from articles like Risso et al. (2018) and Durif et al. (2017), we no longer log_2_-transform the entries of *A* for our eSVD analysis, but rather model the counts in *A* directly after accounting for the library size. Based on our tuning procedure, the curved Gaussian distribution with *k* = 5 and *τ* = 2 best fits the data, determined among a grid of candidate values. When we plot the resulting diagnostic for the eSVD in Figure 7, we obtain results that suggest a much better fit compared to that of the SVD. Specifically, the variance is appropriately increasing with the mean, unlike the trend shown in Figure 2. We conclude that the curved Gaussian model (without a log_2_-transformation) is more appropriate than the constant-variance Gaussian model (with a log_2_-transformation) for modeling our oligodendrocyte dataset.

**Figure 7:**
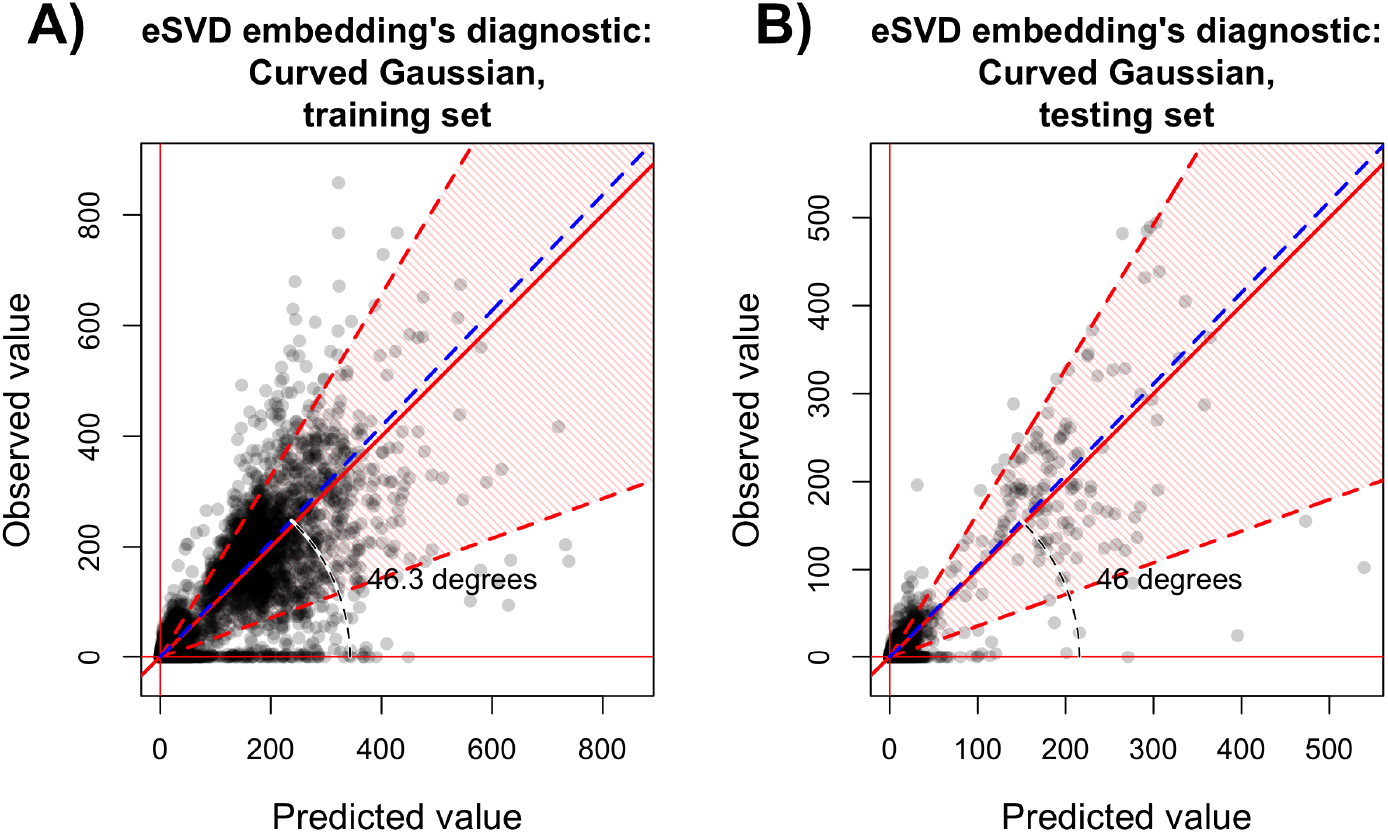
Diagnostic based on matrix completion to assess the fit using the eSVD embedding via the curved Gaussian model with *k* = 5 and *τ* = 2 for either the observed values that are not omitted (i.e., the “training set”) (A) or the observed values that are purposefully omitted (i.e., the “testing set”) (B), both verses their respective predicted values. Both plots are comparable to those in Figure 2. Specifically, the 10th to 90th quantiles of the curved Gaussian model is marked by the shaded red region.

We visualize the eSVD embedding in Figure 8 alongside its estimated trajectories and uncertainty tubes, and we see two distinct trajectories that differentiate among the mature oligodendrocytes. Specifically, when we apply Slingshot to the eSVD embedding, we find that we still retain the conclusion that all cells from Pdgfra+ precursors to myelin-forming oligodendrocytes develop in the same way, similar to Marques et al. (2016). However, in contrast to that work, we are now able to observe substantial differentiation among the mature oligodendrocytes, with two distinct trajectories supported by the uncertainty tubes. Specifically, within this major cell type, only one of the six mature oligodendrocytes sub-types is shared between the two trajectories. Among the five remaining mature oligodendrocytes sub-types, three sub-types branch off in one trajectories while two sub-types branch into the other trajectory. This is in contrast with the analysis using the SVD embedding where all the estimated trajectories lay within one uncertainty tube (Figure 6B). We show additional plots corresponding to these results, as well as follow-up analyses and diagnostics of the oligodendrocytes using UMAP or ZINB-WaVE as well as plots based on the highly informative genes in Appendix H.

**Figure 8:**
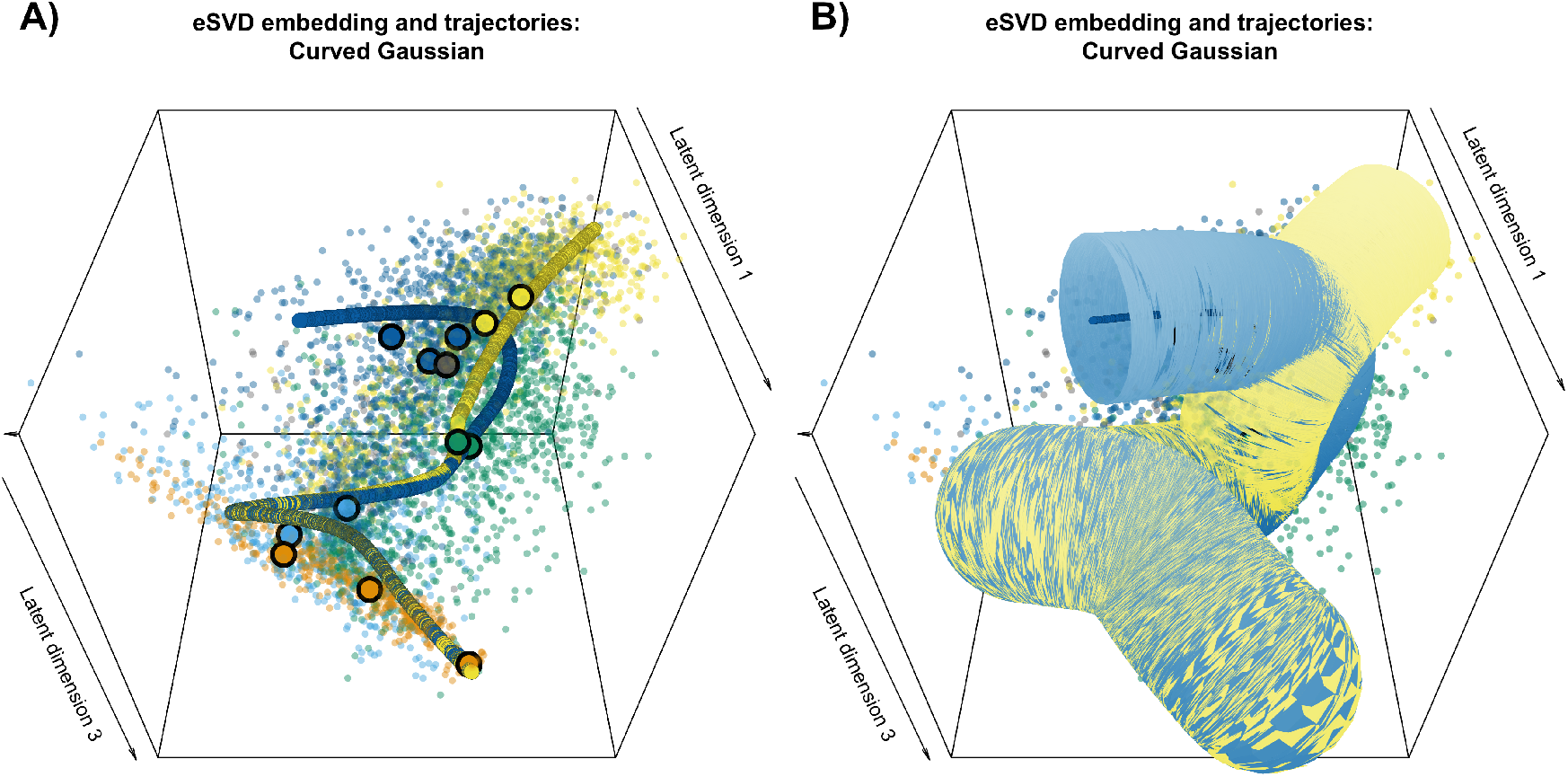
(A) Three-dimensional plot of the estimated latent positions via the eSVD embedding with the curved Gaussian distribution for *k* = 5 and *τ* = 2 with the estimated cell developmental trajectory laid on top. The thirteen bolded points correspond to the cluster centers of the thirteen cell sub-types. The two estimated cell developmental trajectories are colored in yellow and blue. These correspond with the two of six mature oligodendrocytes cell sub-types unique to one trajectory (colored in yellow) and three mature oligodendrocytes cell sub-types unique to the other trajectory (colored in blue). The remaining mature oligodendrocyte cell sub-type is common to both trajectories, prior to the branching (colored in gray). The coloring of cells of other cell types persists from Figure 1. (B) The uncertainty tubes overlaid on top of Figure A. Both plots are comparable to Figure 6.

In summary, from the diagnostic (Figure 7), we conclude that the curved Gaussian distribution is more appropriate for the Marques data, and using this model we identify two distinct developmental trajectories (Figure 8). This is an improvement from the analysis in Marques et al. (2016) which suggested multiple trajectories, but was not able to directly verify this conjecture. Our comparison of the results obtained using the SVD versus the eSVD embeddings can help explain why previous scientific findings suggest that oligodendrocytes effectively follow a single developmental trajectory, while newer analyses based on more flexible statistical models (van Bruggen et al., 2017; Marques et al., 2018) suggest multiple trajectories.

We include an analysis of the single-cell dataset released in Zeisel et al. (2015) in Appendix H to demonstrate eSVD’s performance in a different setting. There, the downstream task is to cluster cells rather than to infer developmental trajectories.

## 8 Discussion

In this article, we develop an estimator to non-linearly embed the cells in a single-cell RNA-sequencing dataset into a lower dimensional space with respect to a random dot product model where the inner product of two latent vectors is the natural parameter of a one-parameter exponential-family distribution F. This embedding method can greatly improve the estimation of cell developmental trajectories overall because it can handle distributions beyond the constant-variance Gaussian distribution, both in theory and practice. While the spirit of such embedding is not new, our contribution is two-fold. First, we develop the eSVD, an alternating minimizing estimator which is computationally efficient and also enables both a tuning procedure based on matrix completion and a theoretical investigation of its statistical properties such as identifiability and consistency. Second, we apply our estimator to analyze the oligodendrocytes in mouse brains, and our results coincide with recent scientific hypotheses (van Bruggen et al., 2017; Marques et al., 2018).

For future work, we plan to further the eSVD both in its modeling flexibility in practice as well as its theoretical properties, as we believe embeddings based on the random dot product model are appealing for single-cell analyses. Specifically, we plan to extend the eSVD to model the dropout effect, allow different nuisance parameters for each gene, or incorporate the library size directly into the statistical model directly. These trends occur in work such as Witten (2011), Pierson and Yau (2015), Townes et al. (2017) and Risso et al. (2018). While the models within these investigations are more flexible, we reiterate that their corresponding estimators were previously often believed to be too complicated to analyze from a theoretic perspective. Therefore, we are interested in studying how flexible our methods can be while still retaining tractability for theoretical analyses or how they perform in misspecified settings. Additionally, our current theory also does not address the trajectory estimation itself, which we think is another promising direction for theoretical investigation. On the methodological side, we plan to provide tuning procedures that are less computationally demanding compared to our current grid-search approach and to work on investigating other downstream applications of the eSVD embedding, such as cell clustering, batch correction, imputation, and RNA velocity (La Manno et al., 2018). Also, while this article focuses trajectory analysis based solely on the cells’ latent vectors, other downstream applications such as gene clustering and finding marker genes could be developed based on the genes’ latent vectors.

## A Code and reproducibility

The code for the method, simulation, and data analysis, as well as the original data used, can be found at https://github.com/linnykos/esvd, in the `eSVD, simulation, main` and `data` folders respectively. The dataset we analyzed was originally collected by Marques et al. (2016), found at the Gene Expression Omnibus with accession number GSE75330 (https://www.ncbi.nlm.nih.gov/geo/query/acc.cgi?acc=GSE75330).

## B Discussion on estimator

## B.1 Initialization method

We first define notation needed to describe the initialization method, inspired by Wang et al. (2016). For a given one-parameter exponential-family distribution, let 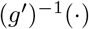 be the inverse function of the *g*′(·), the derivative of the log-partition function for *F*, which is guaranteed to exist by the convexity of *g*(·). Furthermore, for a generic matrix Θ ∈ ℝ^*n*×*p*^ that is rank *k*, let 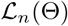 be equivalent to the loss function 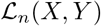 defined in (3.3) for any *X* ∈ ℝ^*n*×*k*^ and *Y* ∈ ℝ^*p*×*k*^ such that 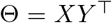. Lastly, define Π_*k*_(·) to be the projection operator (based on alternating between truncating the singular values and thresholding) to project a given matrix onto the set

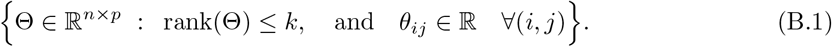

We initialize our estimate, 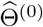, to be 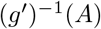, where the function 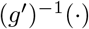 is applied entrywise. Afterwards, we perform projected gradient descent. That is, for *t* ∈ {0,…, *T*′ − 1} iterations, for a stepsize γ > 0, we iterate,

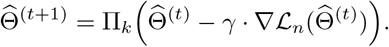

Let 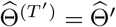 defined in Subsection 4.1 be the initial estimate of Θ.

## Determining γ

In our implementation, we use project gradient steps where within each iteration, γ is selected within each iteration via binary search to be the largest value such that the objective function 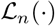 decreases (akin to backtracking line search)

## Lack of convergence

Unfortunately, in our experiments, we have found that there are a handful of instances where the initial projection onto the set (B.1) does not converge empirically. While there is a rich body of literature studying the projection onto the intersection of many convex sets (Kundu et al. (2017) and Tibshirani (2017)), the set of all rank-*k* matrices is nonconvex. In instances of lack of convergence, we terminate the above alternating projection and instead fit a *k*-block model to *g*^−1^(Θ) based applying *k*-means to both the first *k* left and right singular vectors separately. This procedure is reminiscent of those used to fit stochastic block models described in various papers such as Li et al. (2020). We hope to further investigate this projection issue in future work.

We do not use the non-negative matrix factorization (NMF) of *g*^−1^(Θ) since the computational complexity of most non-negative matrix factorization methods is of a similar order to the entire eSVD method. See Subsection B.4 for additional discussion of how NMF compares to the eSVD.

## B.2 Additional variant of tuning procedure

The metric described in Subsection 4.2 uses the angle of the leading eigenvector to assess the quality of the model fit. While this is a useful metric when comparing among different exponential-family distributions *F* (and generally applicable across a broad range of tuning procedures), when the specific distribution *F* is already decided, using the negative log-likelihood can sometimes be another useful metric to use. Specifically, one can compute the negative log-likelihood on all the entries 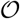 that were omitted during estimation (i.e., the “testing set”), and then choose the model that yields the smallest negative log-likelihood on average across the *B* trials. The advantage of this negative log-likelihood metric over the angle metric described in Subsection 4.2 is that it accounts for the predicted variance among each entry in 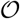. However, as mentioned in Subsection 4.2, the disadvantage is that it is hard to compare the negative log-likelihood across different exponential-family distributions *F*.

## B.3 Usage for common one-parameter exponential-family distributions

The eSVD can be applied to any one-parameter exponential-family distribution, so here, we derive all the necessary ingredients (calculation of the objective function and gradient with respect to *X* and *Y*) for common one-parameter exponential-family distributions. We explain the pipeline to derive how to fit a given one-parameter exponential-family distribution *F* into the eSVD framework.

1. **Writing distribution in exponential-family form:** For a given one-parameter exponential distribution *F*, write the probability density function (or probability mass function) in the form described in (3.2). That is, determine the functions *g*(·) (the log-partition function for *F*), *η*(·) (the natural parameter function) and *T* (·) (the sufficient statistic function) such that

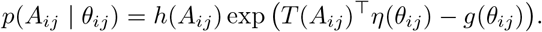
2. **Determine the domain:** Next, based on the log-partition function *g*(·), determine its domain
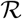.
3. **Determining the objective function:** Next, with the functions *g*(·), *η*(·) and *T* (·) explicitly derived, plug them into the objective function,

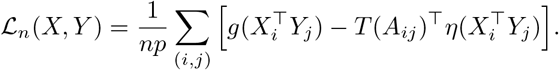
4. **Calculating the gradients:** Lastly, derive the gradient of the above objective function with respect to *X* and *Y*.

Using the above pipeline, we provide the calculations for the following five distributions as an example for readers. We only write down the gradients with respect to *X*, as the gradients with respect to *Y* are similar.

## • Gaussian

For a Gaussian with known variance σ^2^, the density is

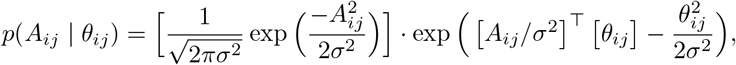

where *θ*_*ij*_ = *μ*_*ij*_, and the domain of *θ*_*ij*_ is 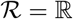. Hence, we can complete the square to derive the objective function, yielding

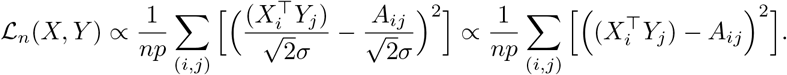

Then the gradient with respect to *X* is a *n* × *k* matrix where the *i*th row is

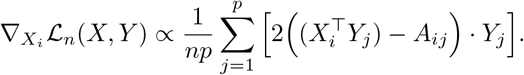

## • Curved Gaussian

For a curved Gaussian with parameter *τ* (i.e., 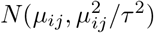), we derive after some algebraic manipulations that the density is,

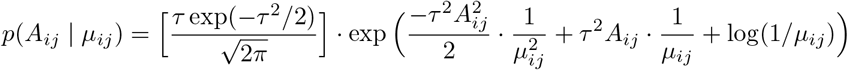

Hence, replacing *θ*_*ij*_ = 1/*μ*_*ij*_ (meaning 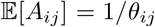, and the domain of *θ*_*ij*_ is 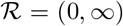. Hence, the objective function becomes

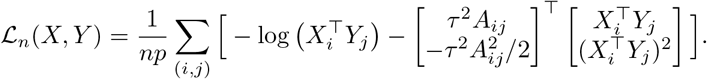

Then the gradient with respect to *X* is a *n* × *k* matrix where the *i*th row is

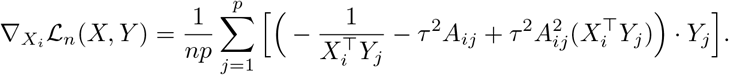

## • Exponential

The density is

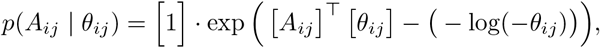

where *θ*_*ij*_ = −λ_*ij*_ (meaning 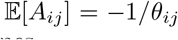, and the domain of *θ*_*ij*_ is 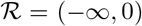. Hence, the objective function becomes

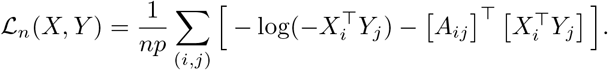

Then the gradient with respect to *X* is a *n* × *k* matrix where the *i*th row is

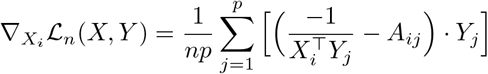

## • Poisson

The density is

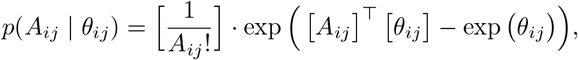

where *θ*_*ij*_ = log(λ_*ij*_) (meaning 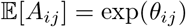, and the domain of *θ*_*ij*_ is 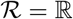. Hence, the objective function becomes

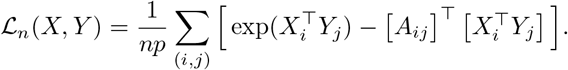

Then the gradient with respect to *X* is a *n* × *k* matrix where the *i*th row is

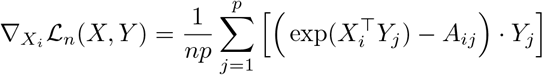

## • Negative Binomial

The negative binomial represents the number of successes before a specified number of failures *r* occurs. (This is different from the binomial distribution which represents the number of success among a fixed number of trials.) This parameter *r* is often also called the dispersion parameter. For a fixed number of failures *r*, the density of Negative Binomial(*r*, *p*_*ij*_) is

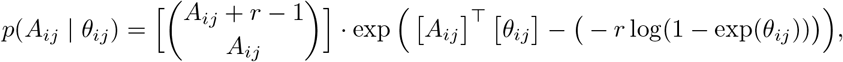

where *θ*_*ij*_ = log(*p*_*ij*_) (meaning 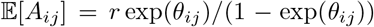, and the domain of *θ*_*ij*_ is 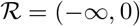. Hence, the objective function becomes

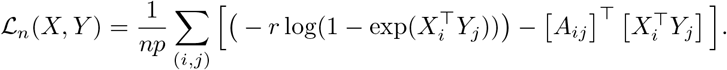

Then the gradient with respect to *X* is a *n* × *k* matrix where the *i*th row is

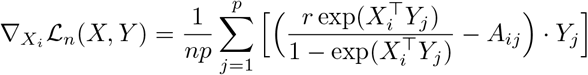

## B.4 Additional comparison of eSVD to estimators in the literature

Continuing Section 3, we discuss additional nuances for the eSVD that makes it different from other methods.

## Comparison to alternating gradient descent

The majority of theoretical work that investigate nonconvex estimators to perform matrix factorization use alternating projected gradient descent to refine the initial estimate where each iteration updates the current estimates with a gradient step (Wang et al. (2016), Yu et al. (2020), and Chi et al. (2019)). This is in contrast with our choice of using alternating constrained minimization in the eSVD. While we have found alternating projected gradient descent is more amendable for theoretical analysis, in practice, we have found it more numerically unstable due to its sensitivity to the chosen step-sizes.

## Comparison to non-negative matrix factorization

There is an extremely rich literature on non-negative matrix factorization (NMF, Donoho and Stodden (2004), Arora et al. (2016), Gillis (2017)), and upon first glance, the eSVD seems similar to NMF. However, there are two importance distinctions. First, the key difference between the eSVD and NMF can be seen in Assumption 3.1. Specifically, the eSVD only assumes that the inner products between *X* and *Y* lie in 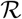, such as the positive half-line. This is in contrast to NMF, where either each entry in *X* or *Y* (or both) lies on the positive half-line. Intuitively, this means the eSVD’s task is easier than NMF’s task (loosely speaking), since ensuring that each entry in *X* and *Y* lying on the positive half-line implies that the inner product between each pair of rows in *X* and *Y* lies on the positive half-line. Second, the eSVD’s identifiability assumptions are more tractable than NMF’s identifiability assumptions. The eSVD’s identifiability assumptions are outlined in Proposition 5.1, which shows that there exists a linear transformation such that the population second moment matrix for *X* and *Y* are equal. NMF’s identifiability conditions are much more nuanced, and leads to concepts such as simplicial cones, separability and anchor words as discussed in Donoho and Stodden (2004) and Arora et al. (2012).

## Comparison to embeddings based on symmetric matrices

There are many embeddings in the literature that are based on symmetric matrices such as covariance matrices, distance matrices or adjacency matrices of neighborhood graphs. These are in contrast with the eSVD as well as other embeddings based on the random dot product model such as ZINB-WaVE (Risso et al., 2018), pCMF (Durif et al., 2017) and NMF. We discuss the broad merits of these two categories of embeddings, and contrast the specific embedding methods with eSVD below. Embeddings based on symmetric matrices can typically be mathematically analyzed in more detail when compared to those for embeddings based on the random dot product models since the former methods typically have more tractable identifiability conditions and often rely simply on the eigen-decompositions, which are mathematically well understood. However, we feel embeddings based on the random dot product models are generally more appealing for single-cell applications since these methods estimate latent vectors for each cell and each gene jointly and can often incorporate more modeling aspects that reflect our understanding of single-cell data. For example, these latter methods can inherently accommodate downstream tasks like bi-clustering and marker gene detection.

## • Comparison to low-rank covariance matrix estimation for exponential families

Liu et al. (2018) and Zhang et al. (2018) estimate a low-rank covariance matrix where each entry in the observed matrix *A* is drawn from an exponential-family distribution. While this task is non-trivial due to the possible dependency between the mean and covariance (which prevents naively centering the variables around 0), this is a different task than our goals posited in this paper since eSVD estimates a low-rank matrix of natural parameters.

## • Comparison to other embedding methods

Methods such as kernel PCA, Isomap (Tenenbaum et al., 2000), t-SNE (Maaten and Hinton, 2008), diffusion maps (Haghverdi et al., 2015), and UMAP (McInnes et al., 2018) also perform non-linear embeddings. We lump these methods together as they generally involve constructing an *n* × *n* symmetric matrix that captures which cells is “neighbors” to which cell or how similar a cell is to another cell given some appropriate notion of similarity. See papers like Lee and Wasserman (2010), Kraemer et al. (2018) and Sun et al. (2019) and the references within for more details. While some of these methods also have rigorous statistical theory (Zwald and Blanchard (2006) and Blanchard et al. (2007)), we have found that in practice, it is hard to assess the quality of fit for these embeddings, and hence also hard to tune the possible parameters of these embeddings effectively. Nonetheless, since these embeddings are often quite successful in practice for complex data, we are interested finding ways to incorporating ideas of the eSVD to these method for future work.

## C Formal description of analysis pipeline

## C.1 Main analysis pipeline

The following procedure describes how we analyze the data to obtain the eSVD-related results in Section 7.

1. **Screening genes:** The dataset starts with 23,556 genes. However, since not all the genes are informative for our analysis, we use the following two methods to select genes, based loosely on Zhu et al. (2019).

- Sparse principal component analysis (Witten et al., 2009): This method uses a tuning parameter that controls the ℓ_1_-norm of the eigenvectors, of which we try 10 values spaced exponentially between 0 and log(23, 556). We choose the model such that the first *K* = 5 sparse eigenvectors involve more than 500 genes and would capture more than 90% of the variance.
- **DESCEND** (Wang et al., 2018): Based on the statistical model developed in Wang et al. (2018) that fits a cubic spline regressing the Gini index onto the log of the estimated mean expression of each gene, we find highly variable genes that deviate from the mean by 50 times the normalized difference. Together, this results in 983 unique genes being selected. In the following parts of the analysis, we will denote the preprocessed data as 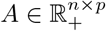 where there are *n* = 5069 cells and *p* = 983 genes.
2. **Rescaling:** We normalize each cell by its library size (i.e., divide each row by its sum) and then multiply the entire matrix by a scalar such that the maximum value is 1000. This is to prevent small numbers from underflowing our method later on. Note that similar to works like Durif et al. (2017) and Risso et al. (2018), we choose to not log_2_-transform our library-normalized matrix *A*.
3. **Tuning the dimensionality of the embedding *k* and the nuisance parameter of the curved Gaussian distribution *τ*** : Omitting four entries per row and per column of *A* for each trial of *b* over a total of *B* = 3 trials, we use the matrix-completion procedure outlined in Subsection 4.2 to select the number of latent dimensions *k* ∈ {3, 5, 10, 20} and nuisance parameter controlling the relation between the mean and variance *τ* ∈ {0.5, 1, 2, 4} (for a total of 16 different parameter settings). We end up selecting *k* = 5 and *τ* = 2. Note that it is important to refit across the different values of *k* since non-linear embeddings are not typically nested, as discussed in Durif et al. (2017).
4. **Embedding via eSVD:** We apply the eSVD where *F* is the the curved Gaussian distribution with *k* = 5 and *τ* = 2 to minimize the loss function (3.3), as prescribed in Subsection 4.1. The initialization procedure is described in Appendix B.1.
5. **Estimating the cell developmental trajectories and uncertainty tube:** Using the thirteen cell sub-types from Marques et al. (2016) as the cluster labels, we apply our modified Slingshot and construct the uncertainty tubes, as alluded to in Section 7 and detailed in Appendix G.

## Screening before rescaling

In contrast to our order of Steps 1 and 2 above, most single-cell preprocessing pipelines, such as ones recommended by Seurat (Butler et al., 2018), often first rescale the counts for each cell and then screen the genes. However, we found preprocessing the data in this fashion hindered the effectiveness of the downstream trajectory analysis based on the eSVD embedding, although the results were qualitatively the same. This is due to the nature of our dataset, where the library size for older cells are substantially higher than younger cells, as seen in Figure 9. By screening genes before rescaling the counts, we select more genes that more informative in distinguishing among the six mature oligodendrocyte cell sub-types. This is critical for our application, as our goal is to distinguish the lineages among the different mature oligodendrocytes.

**Figure 9:**
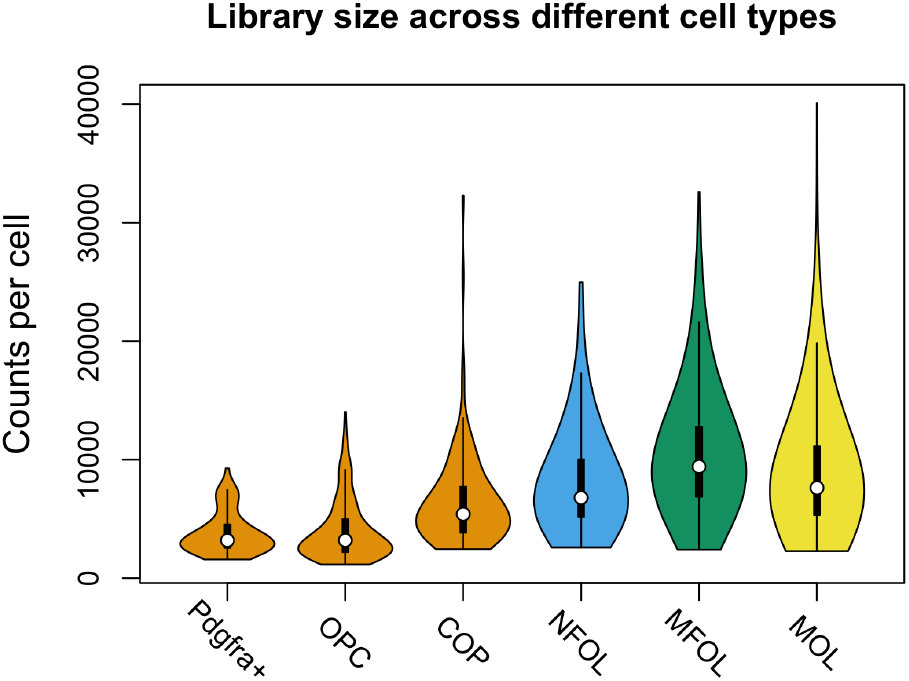
Library size of the raw data across all 5069 cells, split across the six major cell types displayed in Figure 1.

## C.2 Additional details of analysis in Section 2 and SVD-related results in Section 7

We describe additional details of the analysis used to produce the various figures in Section 2, as well as the SVD-related results in Subsection 7.2. The pipeline changes slightly from Appendix C.1 when performing the SVD analysis in the following way:

1. **Different transformation:** As opposed to Step 2’s rescaling, we instead normalize each cell by its library size, and then transform each entry according to *x* → log_2_(10, 000 · *x* + 1), as suggested by work such as Seurat (Butler et al., 2018). We call this the log_2_-transformation in the main text and elsewhere in this supplement.
2. **Different tuning selection:** As opposed to Step 3’s tuning via the eSVD, we instead tune via SoftImpute, a common matrix completion algorithm designed for the constant-variance Gaussian family. This is detailed below under “Details for Figure 2.”
3. **Different embedding:** As opposed to Step 4’s embedding via the eSVD, we use the embedding described in the main text at (2.1) and (2.2), which is the embedding the SVD.

We note the results in Section 2 and Subsection 7.2 are qualitatively the same if we did not use the log_2_-transformation and only use the SVD on *A* directly.

## Details for Figure 1

The scatter plot is produced by the SVD embedding described in (2.1) and (2.2), where the coloring of the points is based on the cell-type information provided in Marques et al. (2016). The contour of the densities estimated based on the `MASS::kde2d` function (using the default bandwidth), where the level of the contour is chosen to be the 92.5% quantile of the estimated density across a grid of points in this two-dimensional space. This quantile level is chosen solely based on the suitably of the figure, and provides the reader a sense of the density of the points that is otherwise hard to gauge based on only the scatter plot.

## Details for Figure 2

SoftImpute (Mazumder et al., 2010) requires the dimensionality of the latent space *k* and a tuning parameter λ to determine the amount of the spectral regularization. To choose this, we try 50 different value of λ from 1 to the value given in `softImpute::lambda0` (a function that computes the smallest value of λ that yields the all-0 estimated matrix), as well as *k* ∈ {5, 10, 20, 50} for a total of 200 different parameter settings. We then choose λ and *k* based on the matrix-completion procedure outlined in Subsection 4.2, where *F* is set to be the Gaussian distribution with constant variance. Importantly, this results in choosing *k* = 50, which is used when fitting the SVD embedding used for downstream analysis in Subsection 7.2. (Note that this is different from *k* = 5 used for the eSVD embedding shown in Subsection 7.3.)

## Details for Figure 3

For this plot specifically, let *A* denote the preprocessed single-cell dataset *prior* to the log_2_-transformation. Figure 3A is created by computing the logarithm of the column-wise (i.e., gene-wise) mean and standard deviation of *A*, the preprocessed single-cell dataset. The color of the point is based on the ANOVA *p*-value that tests if values in a column of *A* (i.e., the expression of each gene) is equal in mean across all six major cell types shown in Figure 1, with blacker points denoting a *p*-value closer to 0 and more yellow points denoting *p*-values closer to 1.

Figure 3B is created by first computing the first principal component of *A* (i.e., the leading eigenvector of the empirical covariance matrix of *A*), setting all the negative entries to 0, and then renormalizing all the entries of the resulting vector to sum to 1. We set all the negative values to 0 so the resulting vector can meaningfully represent a weighted average. We then compute the inner product between the resulting vector and each row of *A*, and then create a violin plot based on the grouping the resulting inner product by the six major cell types shown in in Figure 1.

## D Statistical theory for estimation of matrix of natural parameters

As mentioned in Section 5, an important aspect of interpreting the convergence rate showed in Proposition 5.2 is knowing under what assumptions can we control the estimation error 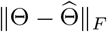. This is the goal of this section, where we operate under the correctly-specified model setting, meaning the eSVD uses the log-likelihood of *F*, the true data-generating distribution. Throughout this section, conditioned on *X* and *Y* (the matrix of random latent vectors for each cell and gene that we are trying to estimate), let the SVD of 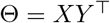 be denoted as

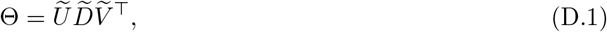

where we denote the singular values as 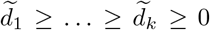. Also, whenever we put a bar over a matrix, as in 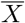 and 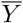, we are explicitly requiring such matrices to have spectral norm 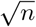 and 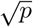 respectively.

The following assumptions ensure that the exponential-family distribution *F* is well-behaved. We define notation to distinguish between sample and population optimizers, borrowing terminology from Balakrishnan et al. (2017). Let 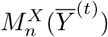 and 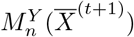 denote *X*^(t+1)^ and *Y*^(t+1)^ respectively in (4.1) and (4.3). In the nonconvex literature, these denote the *sample minimization operators*. Likewise, we define the population loss function,

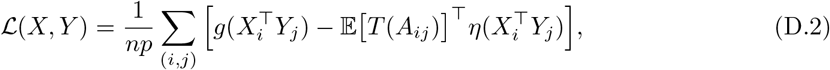

and the corresponding *population minimization operators*,

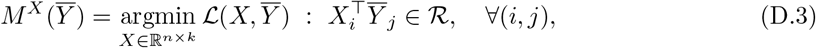

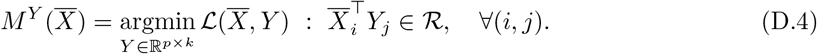

To handle identifiability issues, let us define the set of matrix pairs,

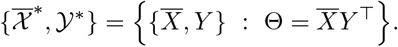

By the SVD of Θ in (D.1), we can see that the set 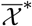 represents all matrices that are equal to 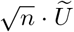 up to rotation. Similarly, we can define the pair of spaces 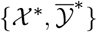. Observe that by the above definitions,

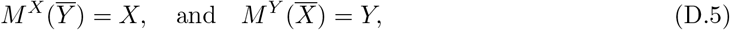

for pairs of matrices 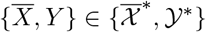 and 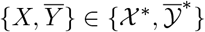.

We now lay down the key assumptions needed to prove the estimation of Θ. While we call these “assumptions,” Assumptions D.1–D.4 should be thought of more as general conditions restricting what kind of random dot product model is allowable. The technical requirements relating all the four assumptions below are stated in Proposition D.1 to come.

### Assumption D.1

(Strong convexity and gradient Lipschitz). *Assume that the population loss function* 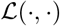 *is μ-strongly convex and its gradient is L-Lipschitz for L* ≥ *μ* > 0 *after fixing one of the input matrices. Specifically, for any matrices X, X′* ∈ ℝ^*n*×*k*^ *and Y* ∈ ℝ^*p*×*k*^ *satisfying Assumption 3.1*,

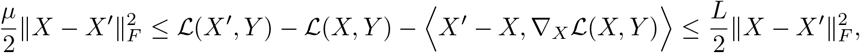

**and a similar assumption holds for any matrices Y, Y*′* ∈ ℝ^*p*×*k*^ *and X* ∈ ℝ^*n*×*k*^.

For the below assumptions, let

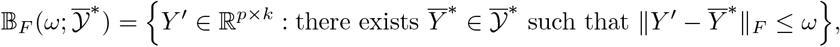

in other words, the union of Frobenius balls of matrices around any matrix in 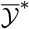 with radius some determined radius *ω*. We invoke a similar definition for 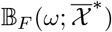.

### Assumption D.2

(Gradient Lipschitz with respect to alternating variable). *Assume that there exists a S > 0 such that for any pairs* 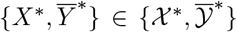 *and* 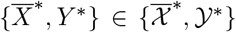 *and any rescaled orthonormal matrices* 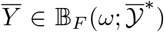 *and* 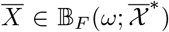 *with spectral norm* 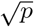 *and* 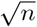 *respectively*

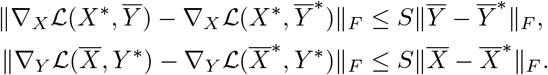

### Assumption D.3

(Uniform statistical error). *Conditioned on X and Y, assume that with probability at least* 1 − *c*/ min{*n*, *p*} *for some universal constant c that*

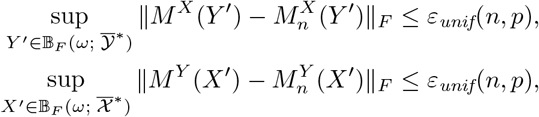

*where ε*_*unif*_(*n*, *p*) *is some function of n and p (and possibly other quantities)*.

In Assumptions D.1 and D.2, the strongly convexity and smoothness enable fast convergence. These assumptions are common in work that study matrix factorization such as Wang et al. (2016) and Yu et al. (2020). Likewise, Assumption D.3 ensures the sample optimizer does not deviate too far from the population optimizer, commonly used in work such as Balakrishnan et al. (2017).

### Assumption D.4

(Initialization condition). *Conditioned on X and Y, assume that for some pair of matrices* 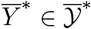*, the initial estimate* 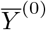 *satisfies*

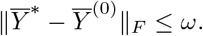

Assumption D.4 is an initialization condition similar to Wang et al. (2016) and Yu et al. (2020) that ensures the alternating minimization steps can allow 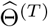 to converge towards Θ. This is described in the following proposition.

### Proposition D.1.

*Conditioned on X and Y, for an appropriate choice of ω under Assumptions 3.1 and D.1–D.4 such that*

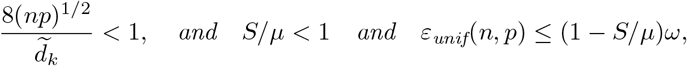

*then for the number of iterations T large enough, with probability at least* 1 - 2*c*/ min{*n*, *p*}, *eSVD described in* (4.1)–(4.4) *achieves the rate*

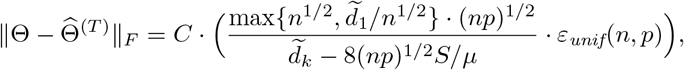

*for some universal constants c and C*.

In short, the above proposition shows that *ε*_unif_(*n*, *p*) is the term that is mainly responsible for driving the rate. To apply Proposition D.1 to a particular model, one needs to compute the terms *μ*, *L*, *S*, and *ε*_unif_(*n*, *p*) for an *ω* that satisfies the conditions in Proposition D.1. This often requires positing assumptions in the context of the particular model that imply Assumptions D.1–D.3. We demonstrate this in the next section with minor modifications to both the estimator and the assumptions.

## E Application of propositions to the curved Gaussian model

In this section, we detail the assumptions needed to apply Proposition D.1 to the curved Gaussian model (4.7). This entails introducing assumptions to help determine the values of *μ*, *L*, and *S* used in Assumptions D.1 and D.2. We introduce some notation beforehand. First, observe that we can rescale the normalization constants in (4.1) and (4.3). Hence, we redefine the sample loss functions to be

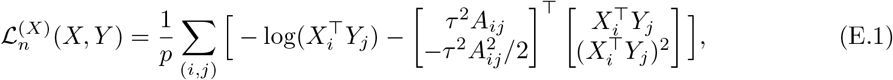

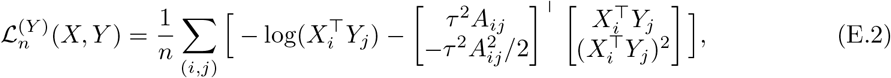

Their corresponding population loss functions that we analyze in this section are

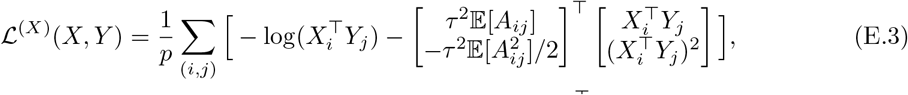

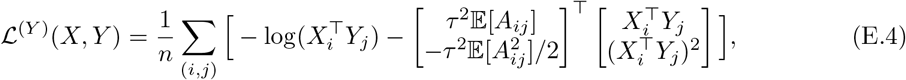

where *A*_*ij*_ follows the distribution (4.7), and the above expectations are understood to be conditioned on Θ. The change from 1/(*np*) in (4.1) and (4.3) to 1/*p* and 1/*n* is for simplicity and facilitates to control the spectrum of the Hessian appropriately. We define the minimization operators we will use in this section as

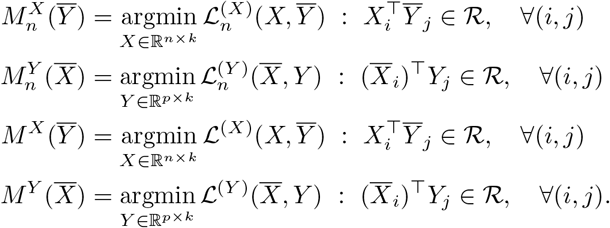

While we state results for both 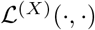 and 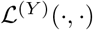 in this section, we prove only statements for 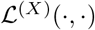 since the proofs for both loss functions are identical.

## Assumptions

The following three assumptions are needed to analyze the curved Gaussian setting.

### Assumption E.1

(Refinement of domain). *Assume for the curved Gaussian distribution* (4.7), *let* 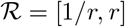 *defined in Assumption 3.1, where there exists a fixed constant r* > 1.

### Assumption E.2

(Inner product error). *For a universal constant c, conditioned on X and Y, assume there exists* 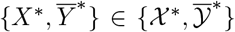 *such that the initialization* 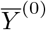 *satisfies for all* (*i, j*) ∈ {1,…, *n*} × {1,…, *p*}

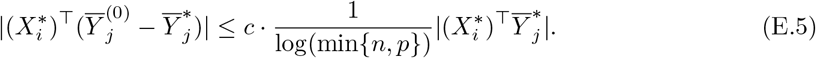

*In addition, conditioned on X and Y, assume for each iterations t* ∈ {1,…, *T*} *throughout the algorithm, there exists* 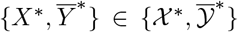 *and* 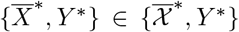 *where* 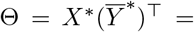 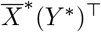 *such that for all* (*i, j*) ∈ {1,…, *n*} × {1,…, *p*}

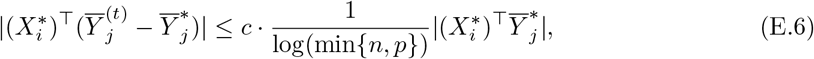

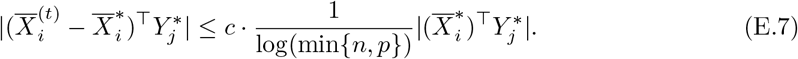

Assumption E.1 effectively ensures 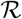 is bounded away from 0, a condition to ensure the Hessian is well-controlled. Assumption E.2 imposes an additional requirement on the original initialization condition in Assumption D.4, and is similar to the incoherence assumption in Ma et al. (2018) and Chi et al. (2019). There, the authors prove that spectral initialization ensures the requirement analogous to (E.5) is met with high probability, and each iteration retains properties analogous to (E.6) and (E.7).

### Assumption E.3

(Fixed statistical error). *Conditioned on X and Y, for any matrices X*′ ∈ ℝ^*n*×*k*^ *or Y*′ ∈ ℝ^*p*×*k*^*, with probability at least* 1 − *c*/ min{*n*, *p*} *for some universal constant c that*

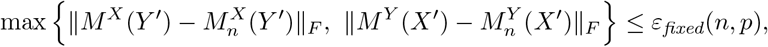

*where ε*_*fixed*_(*n*, *p*) *is some function of n and p (and possibly other quantities)*.

As mentioned in Appendix D, Assumption E.3 is different from Assumption D.3 but enables a simpler analysis. The reason we impose this new assumption is that minimizing objective functions such as (E.1) does not have a closed-form solution, so uniformly bounding the error is difficult for the curved Gaussian model. This simplifying assumption has been used in other work such as Wang et al. (2015). By imposing Assumption E.3 instead of Assumption D.3, eSVD now requires resampling, i.e., a fresh batch of samples every iteration, for all *T* iterations. While this yields a different algorithm that is not practical to use, we believe the theoretical properties we prove also roughly hold for the curved Gaussian model without resampling. We derive the appropriate magnitude of *ε*_fixed_(*n*, *p*) below in Lemma E.3.

## Controlling *μ*, *L*, *S* and *ε*_fixed_(*n*, *p*)

The following lemma implies that 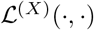 is *μ* = (2 + τ^2^)/*r*^2^-strongly convex and its gradient is *L* = (2 + τ^2^)*r*^2^-Lipschitz.

### Lemma E.1

(Spectrum of Hessian). *For the loss function* (E.3) *and* (E.4) *where A*_*ij*_ *follows the distribution* (4.7), *under Assumption E.1, the eigenvalues of the Hessian* 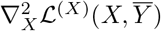 *and* 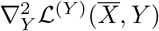 *are bounded between* (2 + τ^2^)/*r*^2^ *and* (2 + τ^2^)*r*^2^.

The following lemma analyzes the gradient smoothness of 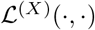 with respect to the alternating variable, assessing the magnitude of *S*.

### Lemma E.2

(Gradient smoothness with respect to alternating variable). *Conditioned on X and Y, for the loss function* (E.3) *and* (E.4) *where A*_*ij*_ *follows the distribution* (4.7), *under Assumptions E.1 and E.2 consider any pairs* 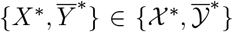 *and* 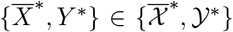. *Then, for* min{*n*, *p*} *large enough, any rescaled orthonormal matrices* 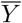 *and* 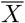 *with spectral norm* 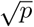 *and* 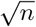 *respectively satisfy*

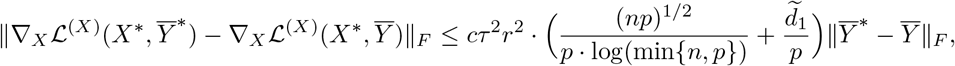

*and*

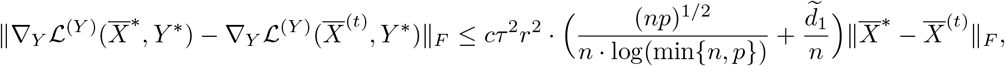

*for some universal constant c*.

We now analyze the difference between the minimizers in (E.3) and (E.1) (or between (E.4) and (E.2)), assessing the magnitude of ε_fixed_(*n*, *p*).

### Lemma E.3.

*Conditioned on X and Y, let A*_*ij*_ *follow the curved Gaussian distribution* (4.7). *For a fixed* 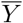, *under the assumptions in Lemma E.1, for p large enough, with probability at least* 1 − 6/*p*,

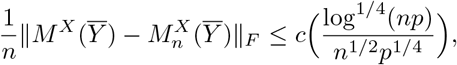

*where c is a constant that depends only on k, τ and r. Similarly, for a fixed* 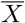, *under Assumption E.1, for n lrage enough, with probability at least* 1 − 6/*n*,

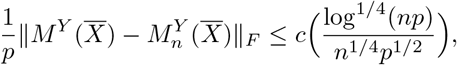

The above corollary means that

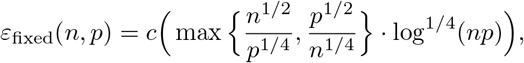

where *c* is a constant that depends only on *k, τ* and *r*. Given the ingredients Lemma E.1 through Lemma E.3, we can derive the following result.

### Corollary E.4

(Application of Proposition D.1 to curved Gaussian model). *Consider the curved Gaussian model* (4.7) *where k, τ, and r are constant, and Assumptions E.1–E.3 hold conditioned on *X* and *Y*. Assuming n* = *O*(*p*) *and p* = *O*(*n*) *are large enough such* 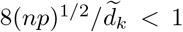, *S/μ* < 1 *and the initialization satisfies* 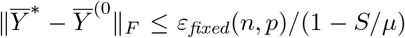, *then for the number of iterations T large enough, the eSVD described in* (4.1)–(4.4) *where each iteration resamples the observed matrix A achieves the rate*

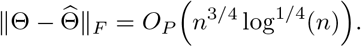

For future work, we believe techniques in Eldridge et al. (2018) and Chi et al. (2019) can help improve our theoretical analysis and simplify the assumptions in the above proposition, as it is not immediately clear yet from a theoretical perspective how often an initialization can satisfy these assumptions. Nonetheless, the following corollary is proved immediately after combining Corollary E.4 with Proposition 5.2.

### Corollary E.5

(Application of Proposition 5.2 to curved Gaussian model). *Assume all the setting and assumptions in Corollary E.4. Then, the eSVD (after reparameterizations* (4.5) *and* (4.6)*) achieves the rate*

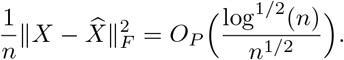

## F Additional simulation details/results

Throughout this section, we use the following notation to parameterize different distributions. The negative binomial and Bernoulli distributions are parameterized by Negative Binomial(*r*, *p*) and Bernoulli(*p*) respectively where *r* is the number of failures (i.e, the dispersion parameter) and *p* is the probability of success. The Gamma distribution is parameterized by Gamma(*a*, *b*) where *a* and *b* are the shape and rate parameters respectively. The Poisson distribution is parameterized by Poisson(λ) where the mean is λ. (Observe that here, the distributions are parameterized by their canonical parameters, not their natural parameters.)

## F.1 Simulation setup

## Generation of natural parameters

To sample from *G* (prior to identifiability conditions), we uniformly sample an equal number of points along 4 connected line segments, where the line segments collectively have endpoints at {(4, 10), (25, 100), (60, 80), (40, 10), (100, 25)} in the Cartesian coordinate system. We then add Gaussian noise with σ = 5 to each of the points. This generates 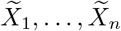 (prior to identifiability conditions).

To sample from *H* (prior to identifiability conditions), we similarly uniformly sample an equal number of points along 2 disconnected line segments. One line segment goes from (1, 4.5) to (1.25, 5) while the other goes from (4.5, 1) to (5, 1.25) in the Cartesian coordinate system. We then also add Gaussian noise to each of the points. This generates 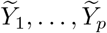 (prior to identifiability conditions). In generating either 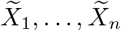 or 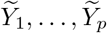, we threshold the values to ensure all entries are positive.

We then compute 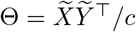, where *c* is a fixed constant chosen beforehand to ensure that the resulting values we generate reasonably lie between 0 and 1000 (depending on which distribution *F* is chosen). Letting the SVD of this matrix be 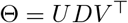, we then output the target embedding we wish to estimate, 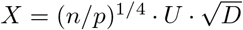 and 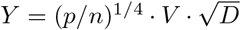, where the square root function is interpreted to be entry-wise.

## Other methods

While SVD, NMF, ICA, UMAP, t-SNE, Isomap and diffusion maps are more common methods in statistical and genomic analyses, ZINB-WaVE and pCMF are methods more specific to single-cell analyses that we briefly overview here.

## • ZINB-WaVE

ZINB-WaVE relies on the negative binomial distribution. Similar to our model, *X*_*i*_ represents the fixed lower-dimensional latent vector for each cell, and *Y*_*j*_ and *W*_*j*_ represent two sets of fixed lower-dimensional latent vectors for each gene. Let *r*_*j*_ denote the dispersion parameter for gene *j*. The generative model is, for any (*i*, *j*) ∈ {1,…, *n*}×{1,…, *p*},

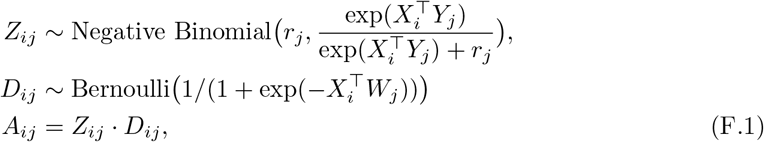

where all the latent variables are independent of one another and *A*_*ij*_’s are all conditionally independent^4^. ZINB-WaVE estimates the parameters *X*, *Y* and *W* via an alternating minimization scheme based on the Broyden-Fletcher-Goldfarb-Shanno (BFGS) quasi-Newton method. We note that the ZINB-WaVE model is able to handle additional covariate information about each cell or gene.

Note, compared to the eSVD model, the ZINB-WaVE model is similar since 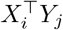 represents the natural parameter of a negative binomial distribution in both models. The key differences are that ZINB-WaVE allows for each gene to have a different dispersion parameter and also models dropout. However, due to the nature of the fitting procedure, ZINB-WaVE does not have a matrix-completion diagnostic to assess the quality of fit (which we developed for the eSVD in Subsection 4.2), and is limited to only modeling the negative binomial distribution.

## • pCMF

pCMF relies on the Poisson distribution. Similar to our model, *X*_*i*_ represents the lower-dimensional latent vector for each cell, and *Y*_*j*_ represents the lower-dimensional latent vector for each gene, but explicit distributions for *G* and *H* are used to facilitate a Bayesian approach. Let π_*j*_ denote the unknown gene-specific dropout rate. The generative model is,

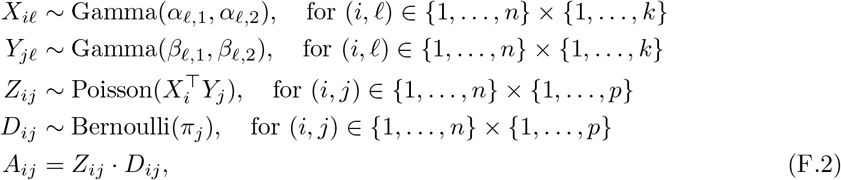

where all the latent variables are independent of one another and *A*_*ij*_’s are all conditionally independent. pCMF estimates the parameters via a variational EM algorithm.

Note, compared to the eSVD model, the pCMF model sets 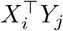 to be the canonical parameter for the Poisson distribution (i.e., the mean), unlike the eSVD which sets 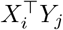 to be the natural parameter. Additionally, pCMF also models dropout. However, similar to the disadvantages of ZINB-WaVE mentioned above, pCMF does not have a matrix-completion diagnostic to assess the quality of fit and is limited to only modeling the Poisson distribution.

We list the `R` packages used for comparisons with other methods. We use the `zinbwave` package for ZINB-WaVE, and set the `K` parameter to 2, the `maxiter.optimize` parameter to 100, the `normalizedValues` parameter to `False`, and `commondispersion` parameter to `False`. We use the `pCMF` package for pCMF and set the `K` parameter to 2 and the `sparsity` parameter to `False`.

For the remaining methods, we use the `dimRed` package for Isomap, ICA (which invokes the `fastICA` package), and NMF (which invokes the `NMF` package), and use the `density` package for diffusion maps, all of which use the default parameters to estimate a two-dimensional embedding. Lastly, for UMAP and t-SNE, we tune the methods’ respective parameters in an oracle-fashion in order to maximize the relative embedding correlation described in Section 6. That is, this tuning requires knowing the true embedding, which is unrealistic in practice but demonstrates the performance of these methods under the most favorable conditions. Specifically, we use the `Rtsne` package for t-SNE, and set the *k* parameter to 2 and tune the `perplexity` parameter in an oracle fashion among 10 values between 2 and 50. We use the umap package for UMAP, and set the `n_components` parameter to 2 as well as `init` to `random`. Additionally, we tune the `n_neighbors` and `min_dist` parameters in an oracle fashion among the values {2, 3, 5, 15, 30, 50} and {10^−5^, 10^−3^, 0.1, 0.3, 0.5, 0.9} respectively (for a total of 36 different parameter settings).

## F.2 First simulation suite: Verification of convergence to *G*

Based on the first simulation suite described in Section 6, we plot the empirical performance of the eSVD when loss function is set to the negative log-likelihood of the negative binomial distribution with *r* = 50, which is correctly-specified model. We plot 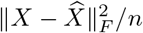 verses *n* in Figure 10.

**Figure 10:**
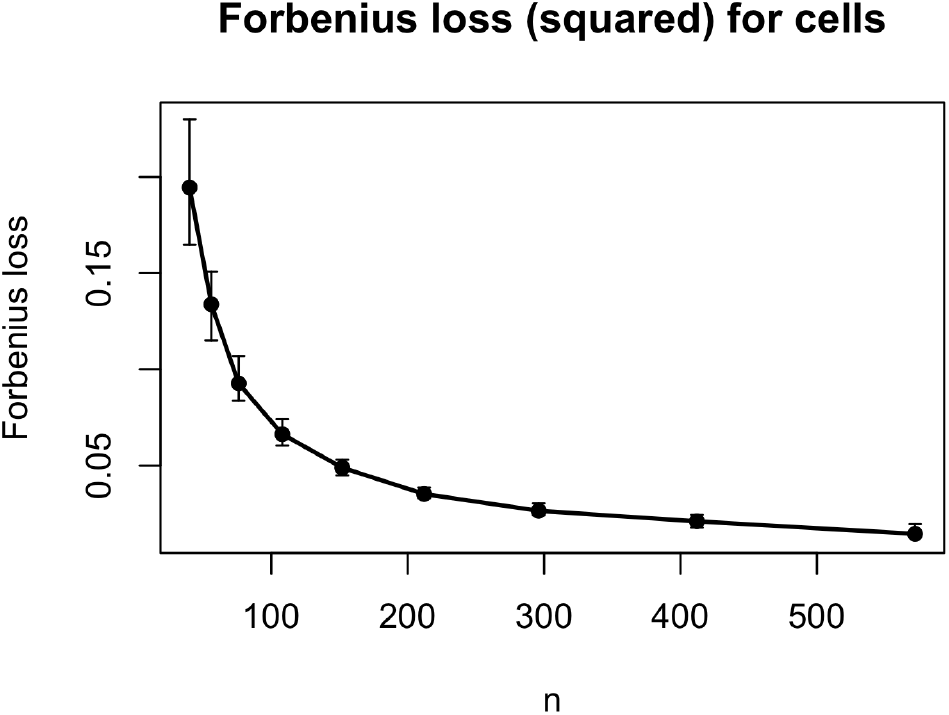
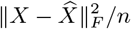 verses *n*, where the solid points represent the median performance over 200 trials and the error bars represent the 25th to 75th quantile.

## F.3 Second simulation suite: Additional comparison of performance under exponential-family models

In this second simulation suite described in Section 6, we compare the eSVD’s performance against the nine other embedding methods where we change both the data-generating distribution and the distribution used by the eSVD simultaneously to be either the Poisson or curved Gaussian distribution. These simulations give us a more holistic understanding of how the data-generating distribution impacts the other nine embedding methods, and also allows us to demonstrate the versatility of the eSVD.

In Figure 11, we see that the eSVD and ZINB-WaVE both perform well in the Poisson simulation, similar to Figure 5. Specifically, despite ZINB-WaVE assuming a negative binomial distribution, it still performs well in this Poisson setting. While it might be surprising that pCMF does not perform well in this setting, as we noted in Appendix F.1, pCMF assumes that 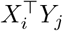 is the canonical parameter of the Poisson distribution (i.e., the mean) while our data-generating model here assumes that 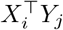 is the natural parameter of the Poisson distribution (i.e., the mean is 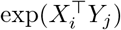, as described in Appendix B.3). This extra exponentiation skews the embedding of the cells outside of the eSVD and ZINB-WaVE, which both operate on the natural-parameter scale.

**Figure 11:**
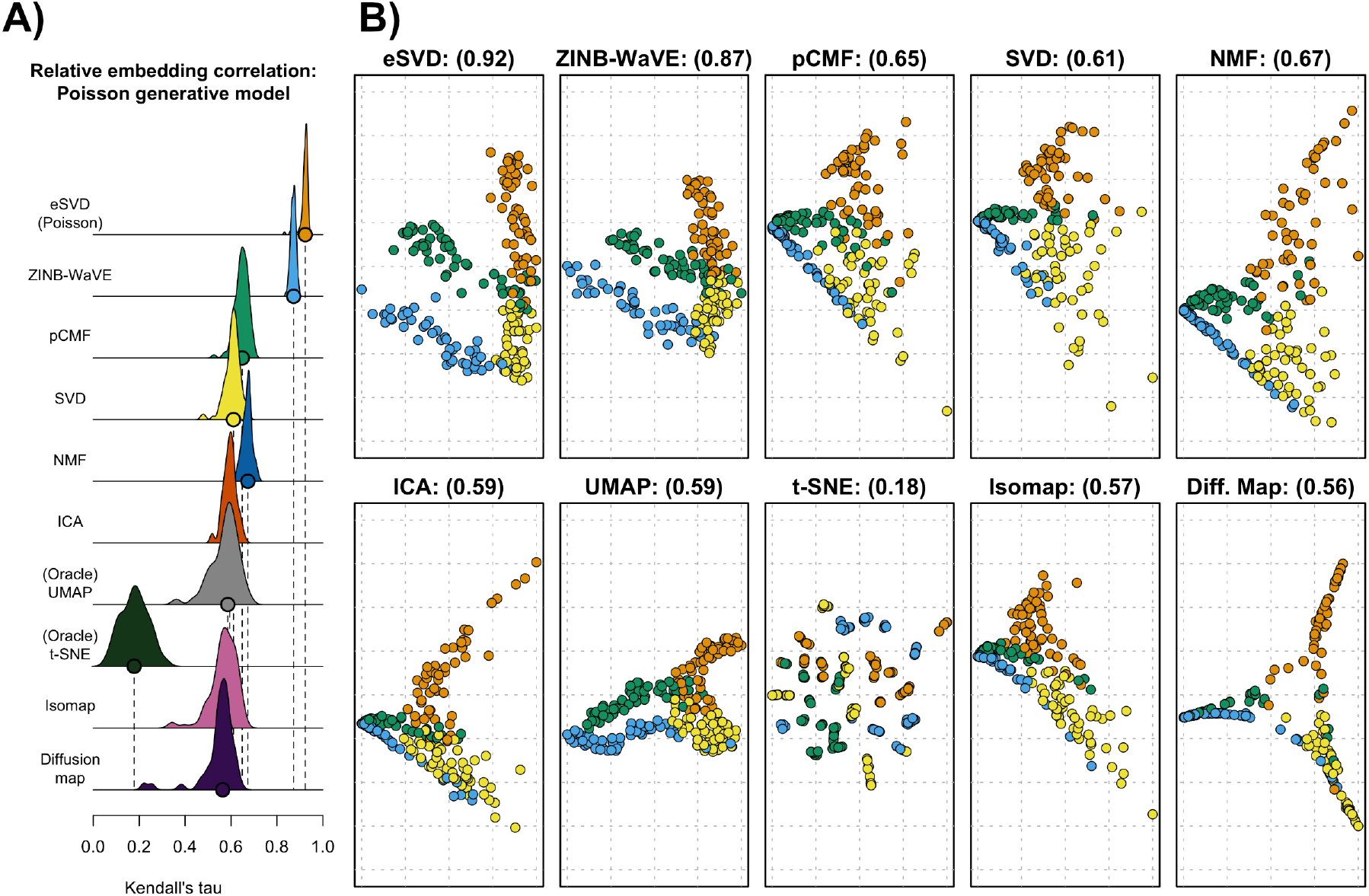
Results for the simulation setting where both the data-generating distribution *F* and the eSVD use the Poisson distribution. The layout of these plots are comparable to those in Figure 5.

In Figure 12, we see that only the eSVD performs well in the curved Gaussian simulation. Here, the true data-generating distribution uses *α* = 2, while the eSVD uses the tuning procedure mentioned in Section 4 to select the most appropriate value of the tuning parameter *α* from the set {1, 2, 4}. We believe the poor performance of the other nine embedding methods is due to the extra variability in the data – the curved Gaussian with *α* = 2 has more variability than the negative binomial distribution or Poisson distribution demonstrated in Figure 5 and Figure 11 respectively.

**Figure 12:**
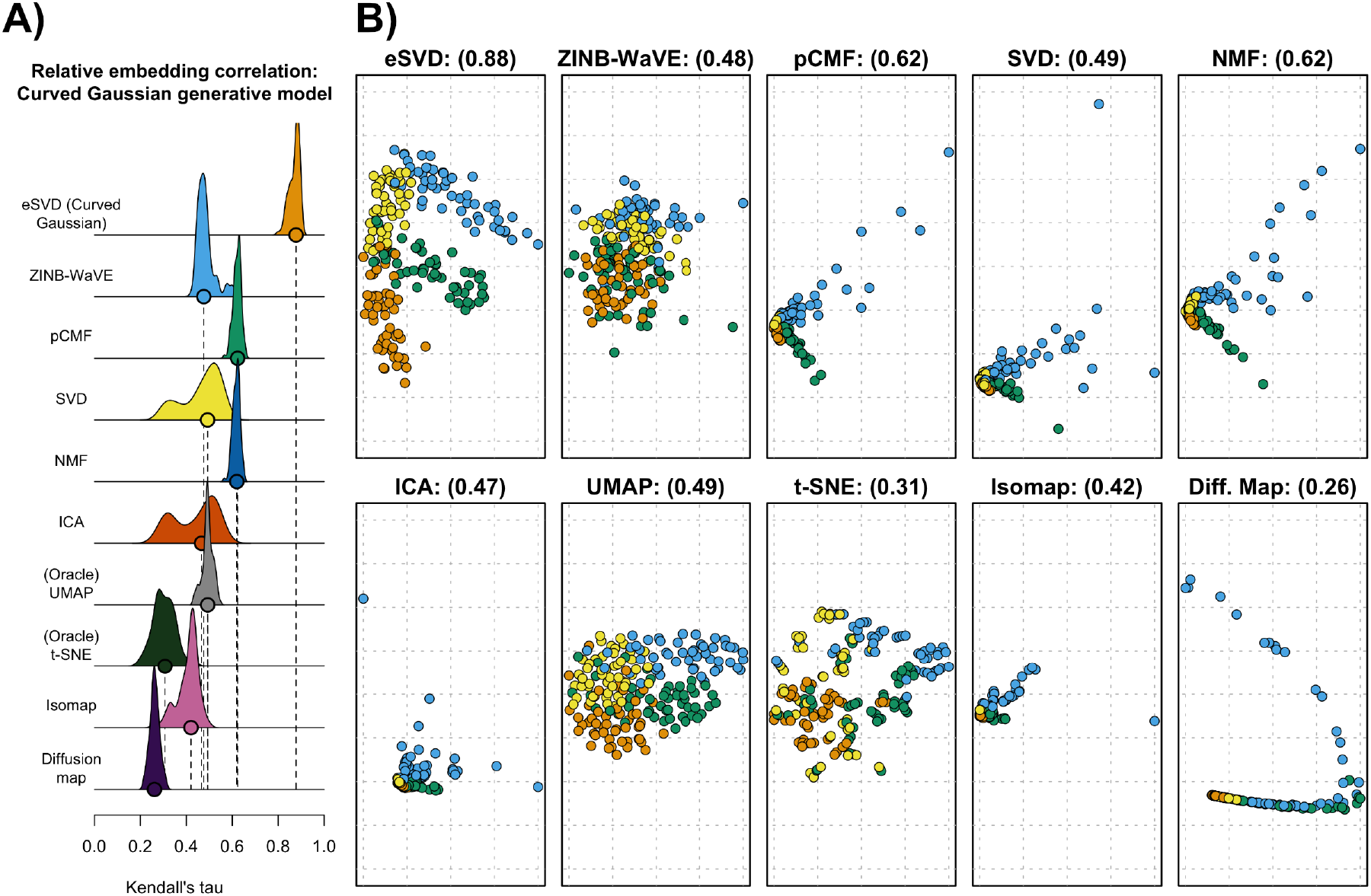
Results for the simulation setting where both the data-generating distribution *F* and the eSVD use the curved Gaussian distribution. The layout of these plots are comparable to those in Figure 5.

## F.4 Third simulation suite: Comparison of performance under misspecified models

We present the additional simulations alluded to as the third simulation suite in Section 6. In this section, we generate models implicitly assumed in ZINB-WaVE or pCMF (see (F.1) or (F.2) respectively). While it is natural to expect that ZINB-WaVE and pCMF will perform the best in their respective simulations, we are additionally interested to see the eSVD can nonetheless be competitive. If so, this gives us empirical evidence that the eSVD’s flexibility and tuning diagnostic are sufficient to capture most of the data’s nuances assumed in these models, despite being misspecified.

## ZINB-WaVE model

In this simulation setting, we build on top of the negative binomial model descried in Section 6, but include additional complexities that make the simulation more realistic according to the generative model for ZINB-WaVE shown in (F.1). Specifically, using the ZINB-WaVE model (F.1), we vary the dispersion parameter so 25% of the genes have a dispersion parameter of *r* = 80, another 25% have a parameter of *r* = 120, and the remaining 50% have a parameter of *r* = 600. Additionally, a dropout term is now included based on a logistic model, unlike the model described in Section 6. Both of these changes ensure that the negative binomial model that the eSVD fits is sufficiently misspecified. As our simulations reassuringly shows though, the eSVD nonetheless estimates the embedding relatively well when compared to other methods aside from ZINB-WaVE itself.

When we fit the eSVD via a negative binomial model, we use the tuning procedure mentioned in Subsection 4.2 to search for a global dispersion parameter of *r* ∈ {50, 100, 500, 1000} and set *k* = 3. Notice that we are using *one* dispersion parameter to model the simulated dataset, although our true generative model uses three different dispersion parameters. Also, we increase the latent space to *k* = 3 since we found empirically, the addition of the dropout factor can be reasonably captured by an extra latent dimension.

We demonstrate our results in Figure 13. We see that ZINB-WaVE performs the best according to the relative embedding correlation metric, but this is unsurprising as our generative model is correctly specified for ZINB-WaVE. However, even though it is misspecified for the eSVD using the negative binomial distribution, the eSVD’s performance is still quite competitive according to our relative embedding criterion, compared among the nine remaining methods. Hence, we believe that the eSVD using the negative binomial distribution, with our tuning procedure, is comparable in performance to ZINB-WaVE in practice. As we mentioned in the main text however, the benefits of using the eSVD is that the eSVD has a solid theoretical foundation, can be easily extended to other one-parameter exponential-family distributions, and can handle missing values to assess model fit.

**Figure 13:**
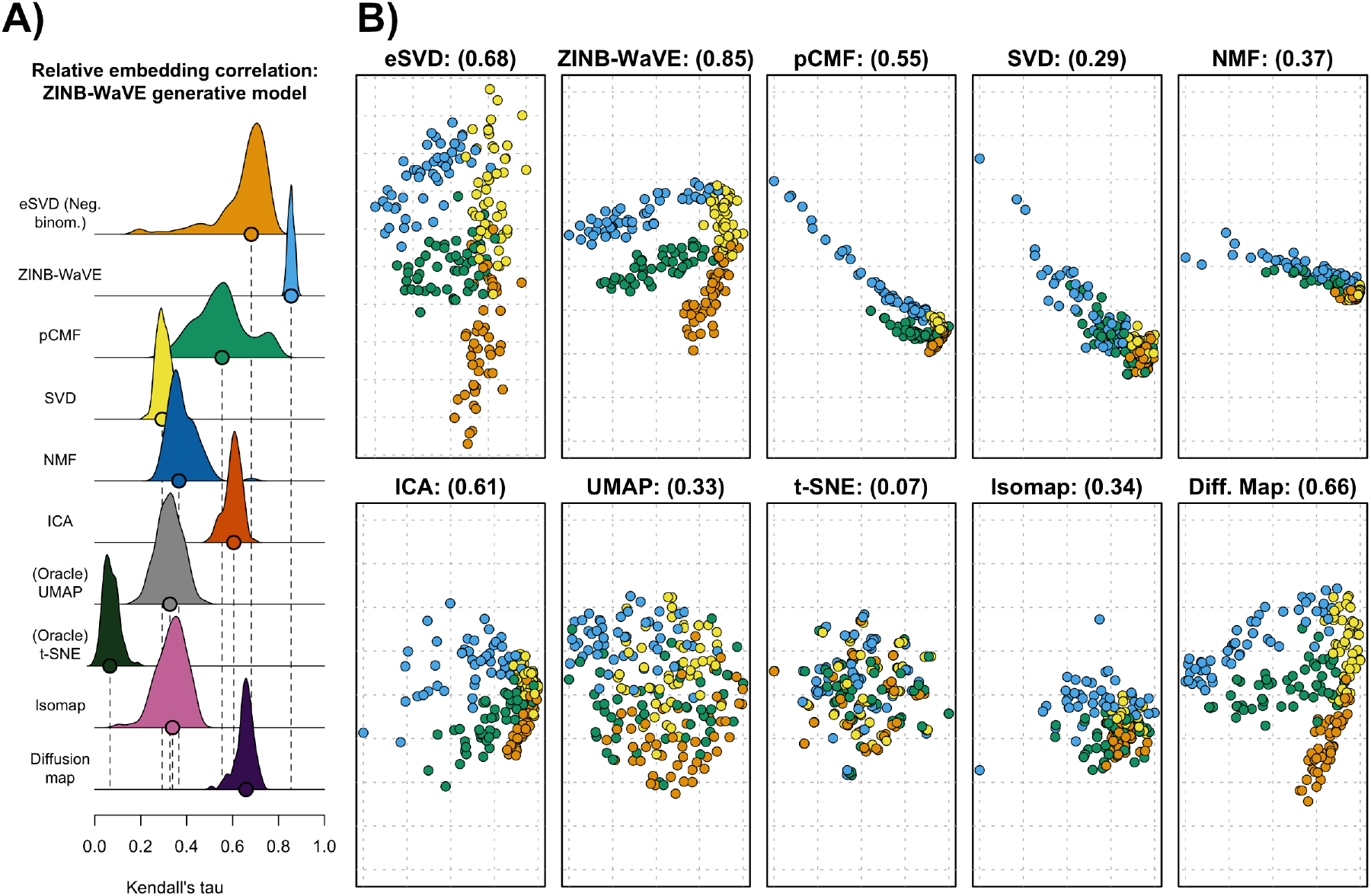
Results for the misspecified simulation setting where the data-generating distribution *F* is generated according to the assumed model by ZINB-WaVE (F.1) and the eSVD use the negative binomial distribution. The layout of these plots are comparable to those in Figure 5.

## Effect of *k* and appropriateness of tuning procedure

To demonstrate the appropriateness of choosing *k* = 3 in the above simulation under the ZINB-WaVE model, we empirically compare how different *k* can effect the quality of the model fit. (We have found this investigation more meaningful under the misspecified model since the effect of varying *k* was not apparent under any of the settings in the second simulation suite.) Specifically, while there is no “correct” value of *k* due to the model misspecification, we are interested if our tuning procedure can select the *k* that yields the most accurate embedding and assess how choosing *k* poorly can impact the embedding.

For this simulation, we retain the ZINB-WaVE model above and use the eSVD with the negative binomial distribution. However, we now use the tuning procedure mentioned in Subsection 4.2 to search for a global dispersion parameter of *r* ∈ {50, 100, 500} as well as *k* ∈ {2, 3, 10}, where the quality of the model fit is computed using the negative log-likelihood as described in Appendix B.2. We use this metric as opposed to the principal angle in this particular simulation since we have already fixed the eSVD to use the negative binomial distribution.

The results in Figure 14 demonstrate that our tuning procedure selects *k* = 3 92% of the time (among all 100 trials), which also yields the embedding with the highest relative embedding correlation on average, and selects *k* = 2 among the remaining trials. Specifically, in Figure 14, the selected pair of parameters (*r*, *k*) based on the negative log-likelihood also often has a high relative embedding correlation. All-in-all, Figure 14 demonstrates why we chose *k* = 3 for the eSVD when producing Figure 13, and demonstrates how our tuning procedure can select *k* that can yield the best-performing embedding despite the model misspecification.

**Figure 14:**
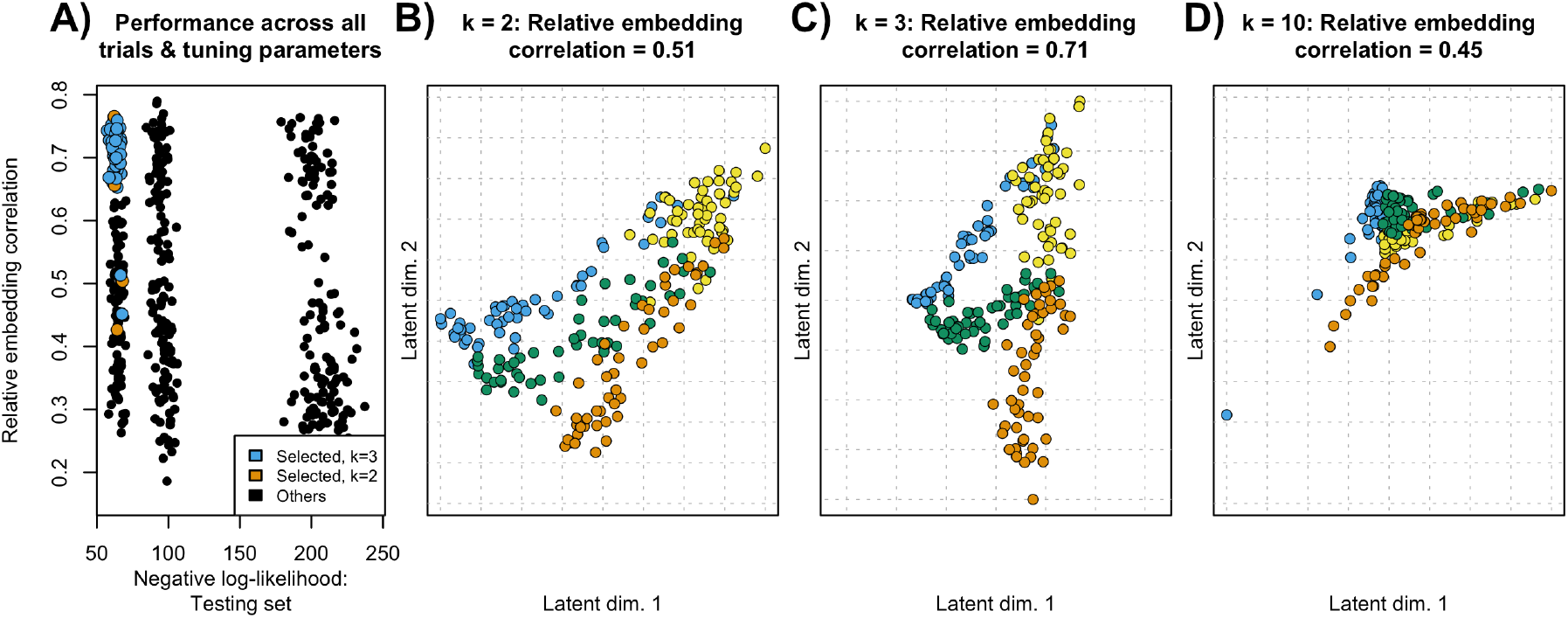
(A) The relative embedding correlation plotted against the negative log-likelihood of the testing set, where each point represents one of 100 trials fitting the eSVD using a different pair of parameters (*r*, *k*) among {50, 100, 500} × {2, 3, 10}, for a total of 900 points shown. Among each of the 100 trials, the selected (*r*, *k*) parameter pair’s corresponding point is highlighted in color, blue for if *k* is chosen to be 3, and orange if *k* is chosen to be 2. (B to D) The embedding for *k* = 2, *k* = 3 and *k* = 10 respectively, trial with the median accuracy, and *r* = 50 (the selected value of *r* according to our tuning procedure). The layout of these plots are comparable to those in Figure 5.

## pCMF model

In this simulation setting, we make the simulation more realistic according to the generative model for pCMF shown in (F.2). Specifically, we set π_*j*_ = 0.3 for all *j* ∈ {1,…, *p*} and use the same way to generate *X*_1_,…, *X*_*n*_ and *Y*_1_,…, *Y*_*p*_ as previously described in Appendix F.1, instead of generating the latent vectors from the prescribed prior distribution. While we set the eSVD to estimate the embedding according the Poisson setting in this simulation, there are two important aspects that makes this a misspecified setting. First, pCMF assumes that the mean of the Poisson distribution is 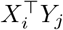, while the eSVD under the Poisson model assumes that the mean of the Poisson distribution is 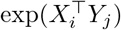. Second, a dropout term is now included. Both of these changes ensures that the Poisson model that the eSVD fits is sufficiently misspecified. As our simulations reassuringly shows though, the eSVD nonetheless estimates the embedding relatively well when compared to other methods aside from ZINB-WaVE itself. Similar to in the ZINB-WaVE simulation above, when we fit the eSVD via a Poisson model, we set *k* = 3, where we found that the extra latent dimension helps to capture the dropout effect.

We demonstrate our results in Figure 15. Similar to before, we see that pCMF performs the best, but this is unsurprising. However, more surprisingly, we see that many embedding methods actually work well in this simulation setting, since the Poisson distribution has a small variance when compared to the negative binomial or curved Gaussian distribution. Hence, the additional variability in the data is insufficient from dramatically distorting many embedding methods, including SVD, NMF, ICA and UMAP. While the eSVD does not rank among the best-performing methods in terms of the relative embedding correlation metric here (Figure 15A), we see that the eSVD’s estimated embedding visually resembles the true layout of latent vectors (Figure 15B, compared to Figure 4). Hence, given this particular simulation as well as all the simulations presented beforehand, we feel the flexibility of the eSVD makes it a competitive embedding method overall.

**Figure 15:**
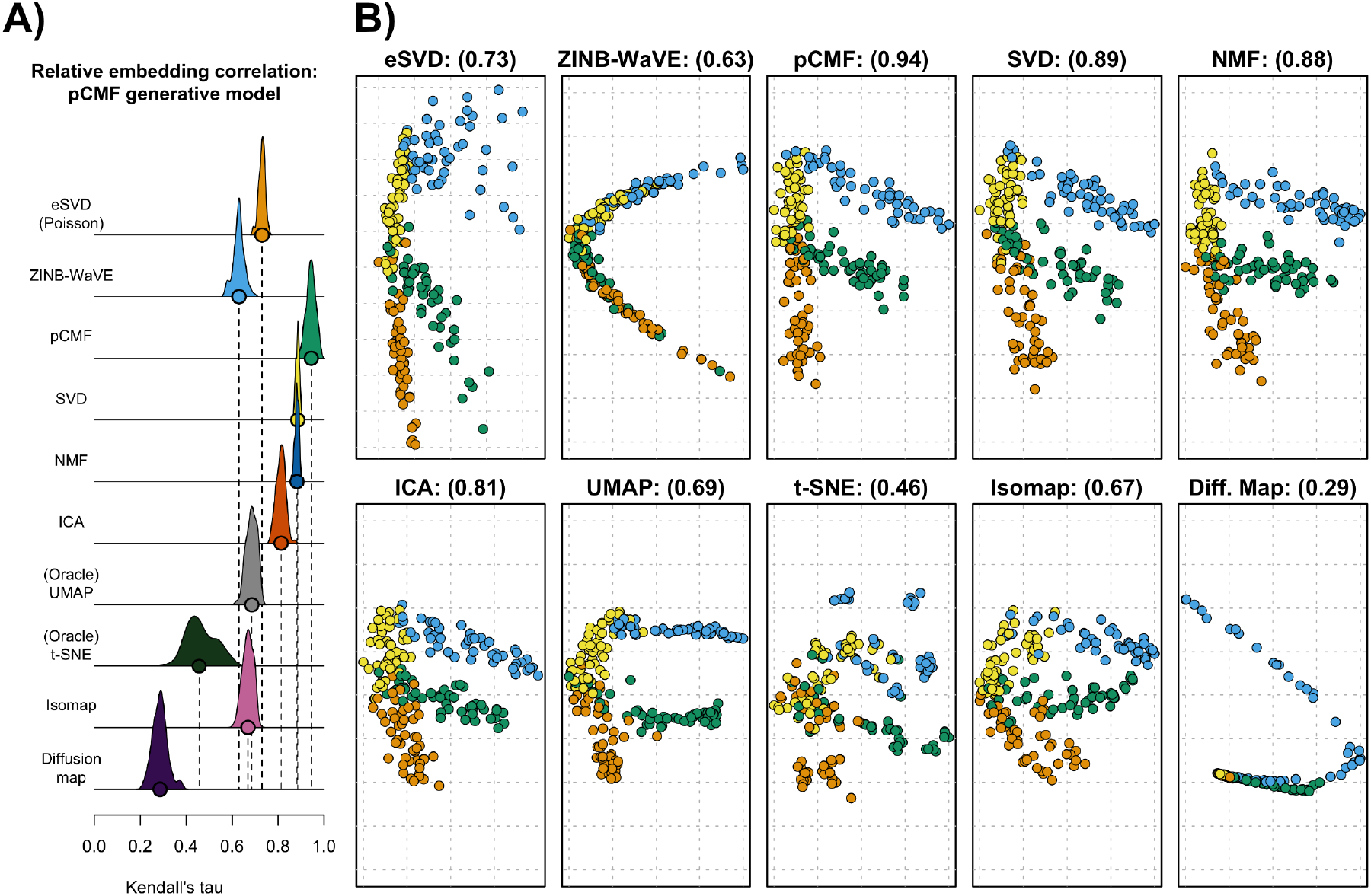
Results for the misspecified simulation setting where the data-generating distribution *F* is generated according to the assumed model by pCMF (F.2) and the eSVD use the Poisson distribution. The layout of these plots are comparable to those in Figure 5.

## G Details on Slingshot and uncertainty tube

## G.1 Modifications to Slingshot

Our implementation of Slingshot differs from its original implementation in Street et al. (2018) in a few aspects. The first two modifications is related to how the lineages (i.e., what the different branches are and the ordering of the cell sub-types) are estimated, and the last modification is related to how the trajectories are estimated given the lineages (i.e., what is the numeric curve that interpolates the points in the latent space).

## • Respecting natural order

Our implementation respects the order among the major cell types (i.e., Pdgfra+ precursor, oligodendrocyte precursor cells, differentiation-committed oligodendrocyte precursors, newly formed oligodendrocytes, myelin-forming oligodendrocytes, and mature oligodendrocytes), but allows the lineage to arbitrarily order or split the cell sub-types within the same cell type.

## • Construction of lineage

Our implementation determines the lineage via a shortest path tree from the starting cluster, where the distance is determined by the multivariate Hotelling’s *t*-test for unequal covariances. This is in contrast with the original Slingshot that uses a minimum spanning tree, where the distance is determined by the multivariate Hotelling’s *t*-test for equal covariances.

## • Enable upsampling

The original Slingshot does not weight clusters, so the lineage curves naturally gravitated towards the larger clusters. To compensate for this phenomenon, we upsampled the cells in each cluster via sampling with replacement so each cluster has the same number of cells. This effectively adds larger weights to these smaller clusters so each cluster is treated equally regardless of its original size. A similar modification is done in Herman and Grün (2018).

## G.2 Construction of uncertainty tube

We construct the uncertainty tubes via a bootstrap-based approach to determine if the different cell developmental trajectories are substantially different. This is done via the following procedure.

## 1. Bootstrapping

For a specific trial, sample with replacement the low-dimensional embedding 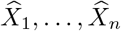 among each cell sub-type. This generates a new dataset with the same proportion of cell sub-types. Run Slingshot on this new dataset, having fixed the order of the cell sub-types in each lineage, to obtain a new set of trajectories.

## 2. Computing the quantile of ℓ_2_ distance between lineage curves

Let 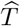 be one of the estimated trajectories based on the original estimated embedding 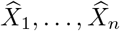. For each estimated trajectory *T*_*b*_ from the newly generated dataset that matches 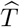 based on the order of cell sub-types, compute the pointwise ℓ_2_ distance between the two respective curves. Specifically, letting {*T*_*b*,1_,…, *T*_*b*,*N*_} and 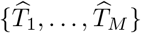 denote the *N* and *M* discrete *k*-dimensional points in order that represent the trajectory *T*_*b*_ and 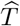 respectively, this pointwise ℓ_2_ distance is computed as the 95% quantile of the following set

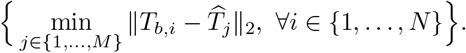

After computing this distance for each of the bootstrapped trajectories, let 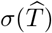 be the the 95th quantile ℓ_2_ distance among all the bootstrapped trajectories, denoting the “margin of error” for a particular original estimated trajectory 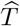. Let σ be the 95th quantile of such values among all the original estimated trajectories, representing the width of the uncertainty tubes.

## 3. Computing the lineage tube

For each lineage in the original dataset, construct a “tube” of radius σ around the lineage curve.

## H Additional analysis results

## H.1 Additional plots of results

In Figure 16 and Figure 17, we show the two-dimensional and three-dimensional plots of the estimated embeddings to provide more clarity to Figure 1 and Figure 6 in the main text.

**Figure 16:**
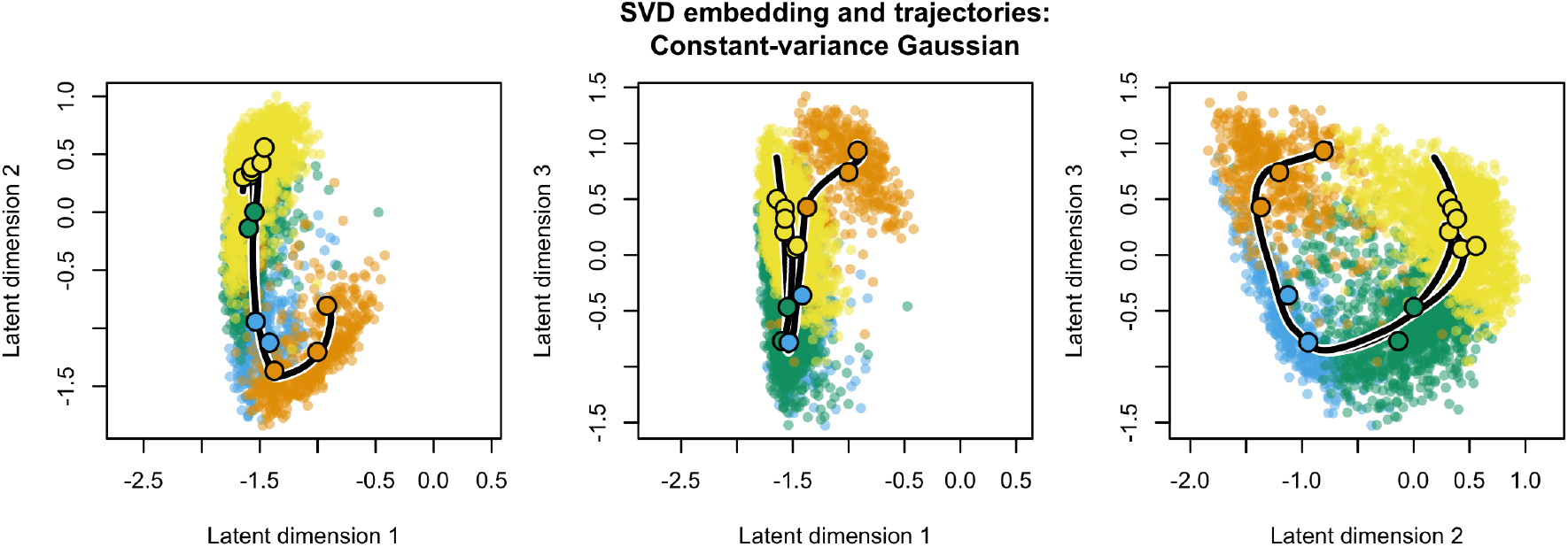
Two-dimensional plots of the SVD embedding, which corresponds to the three-dimensional plots shown in Figure 6. Each cell is color coded using the same color scheme from the aforementioned figure, the large dots represent the cluster centers for each of the thirteen cell sub-types, and the estimated trajectories are overlaid on top, which correspond to the same trajectories in the aforementioned figure. The three plots correspond to the three pairs of latent dimensions, one of the plots being the same as Figure 1.

**Figure 17:**
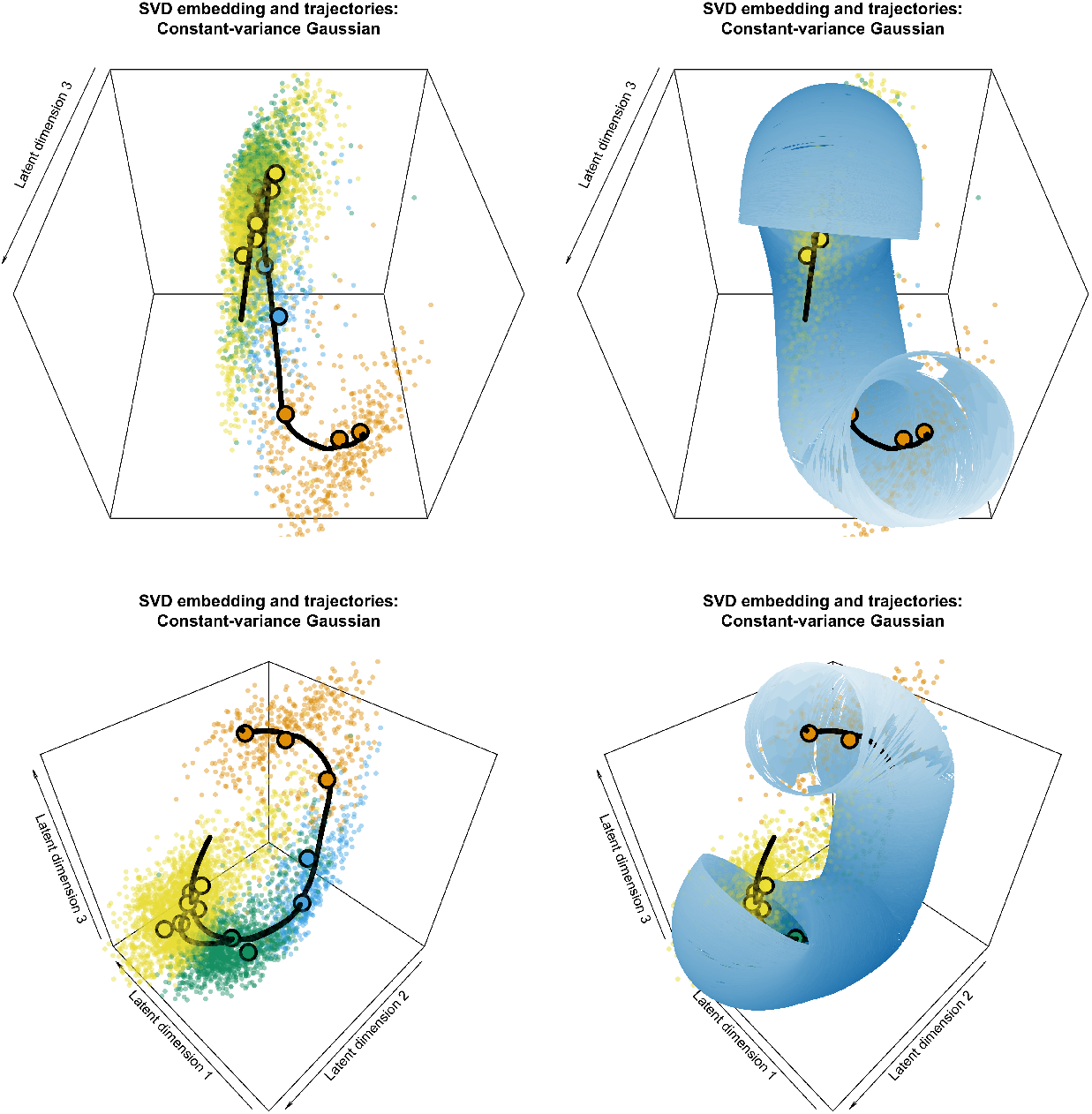
Three-dimensional plots of the estimated latent positions via the SVD embedding, without and with the uncertainy tube overlaid ontop. This plot is of the same estimated embedding as shown in Figure 6 but shown from a different perspective (each perspective represented by a different row).

In Figure 18 and Figure 20, we show the two-dimensional and three-dimensional plots of the estimated embeddings to provide more clarity to plots shown in Figure 8 in the main text.

**Figure 18:**
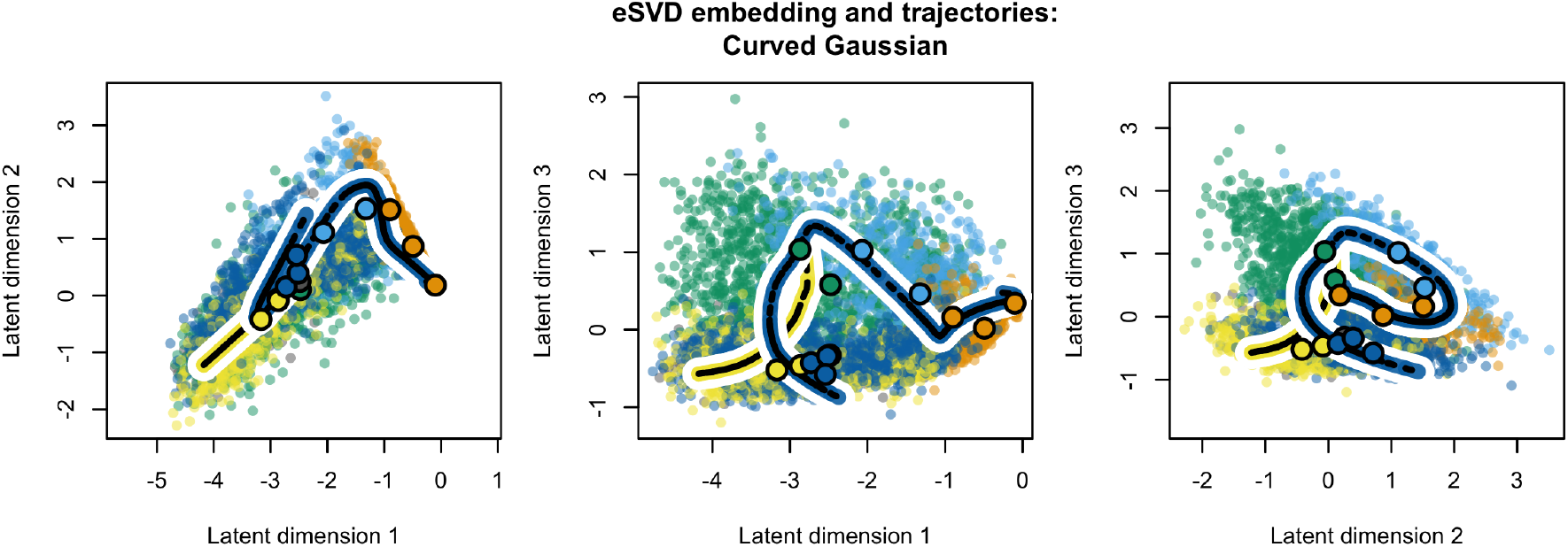
Two-dimensional plots of the eSVD embedding using the curved Gaussian distribution with *k* = 5 and *τ* = 2, which corresponds to the three-dimensional plots shown in Figure 8. Each cell is color coded using the same color scheme from the aforementioned figure, the large dots represent the cluster centers for each of the thirteen cell sub-types, and the estimated trajectories are overlaid on top, which correspond to the same trajectories in the aforementioned figure.

**Figure 19:**
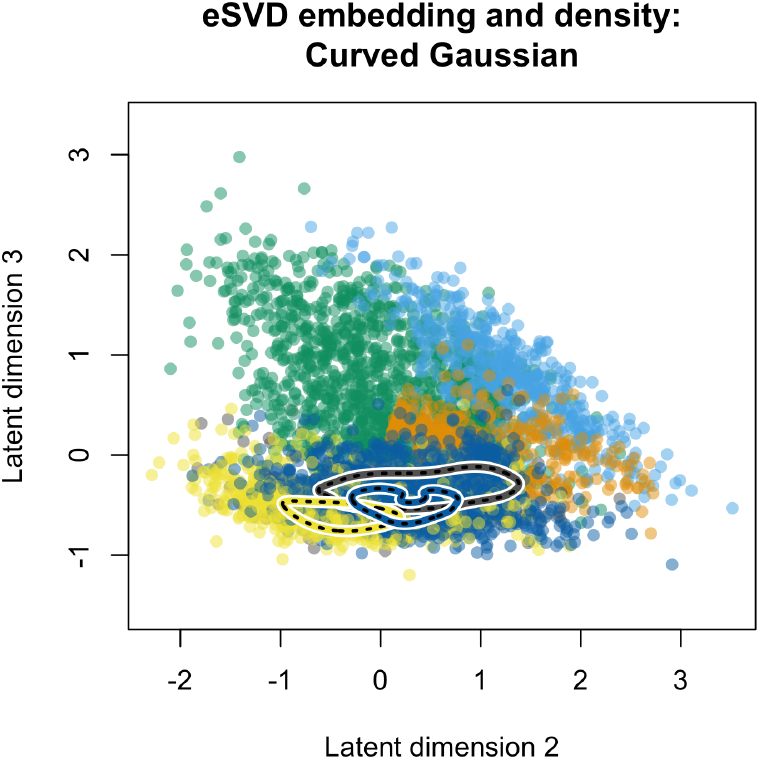
Two-dimensional plot of the eSVD embedding, similar to that shown in Figure 18, but with high-density regions denoted for the mature oligodendrocytes shared between the two trajectories (gray points), and the mature oligodendrocytes unique to each of the estimated trajectories (either blue or yellow points). This plot is directly comparable to Figure 1.

**Figure 20:**
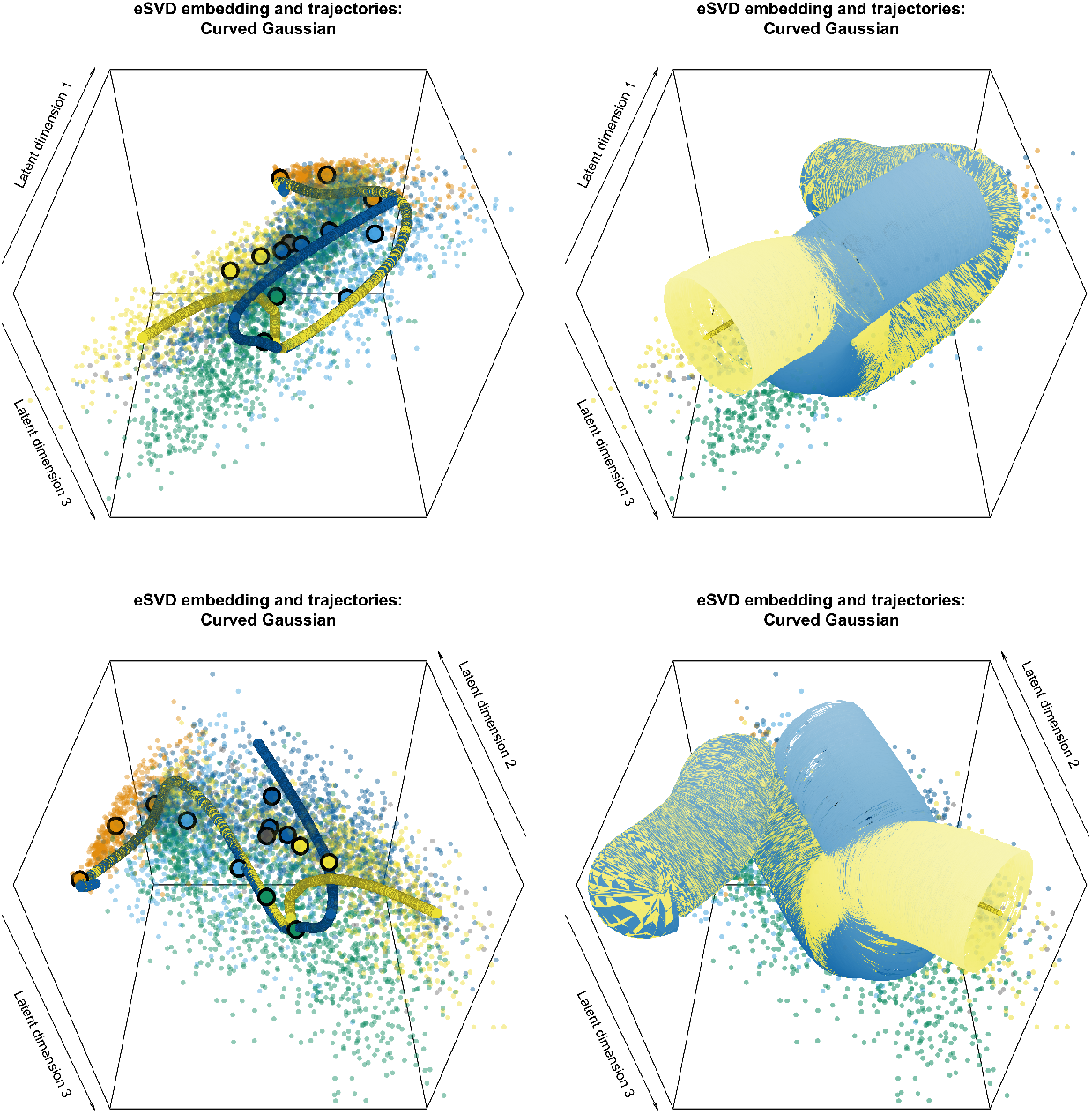
Three-dimensional plots of the estimated latent positions via the eSVD embedding, without and with the uncertainy tubes overlaid ontop. This plot is of the same estimated embedding as shown in Figure 8 but shown from a different perspective (each perspective represented by a different row).

Additionally, Figure 21 shows the same plot as Figure 7, except zoomed in to display the behavior of points near zero. We see that the same trends persists from Figure 7 (mainly, the variance increasing with the mean).

**Figure 21:**
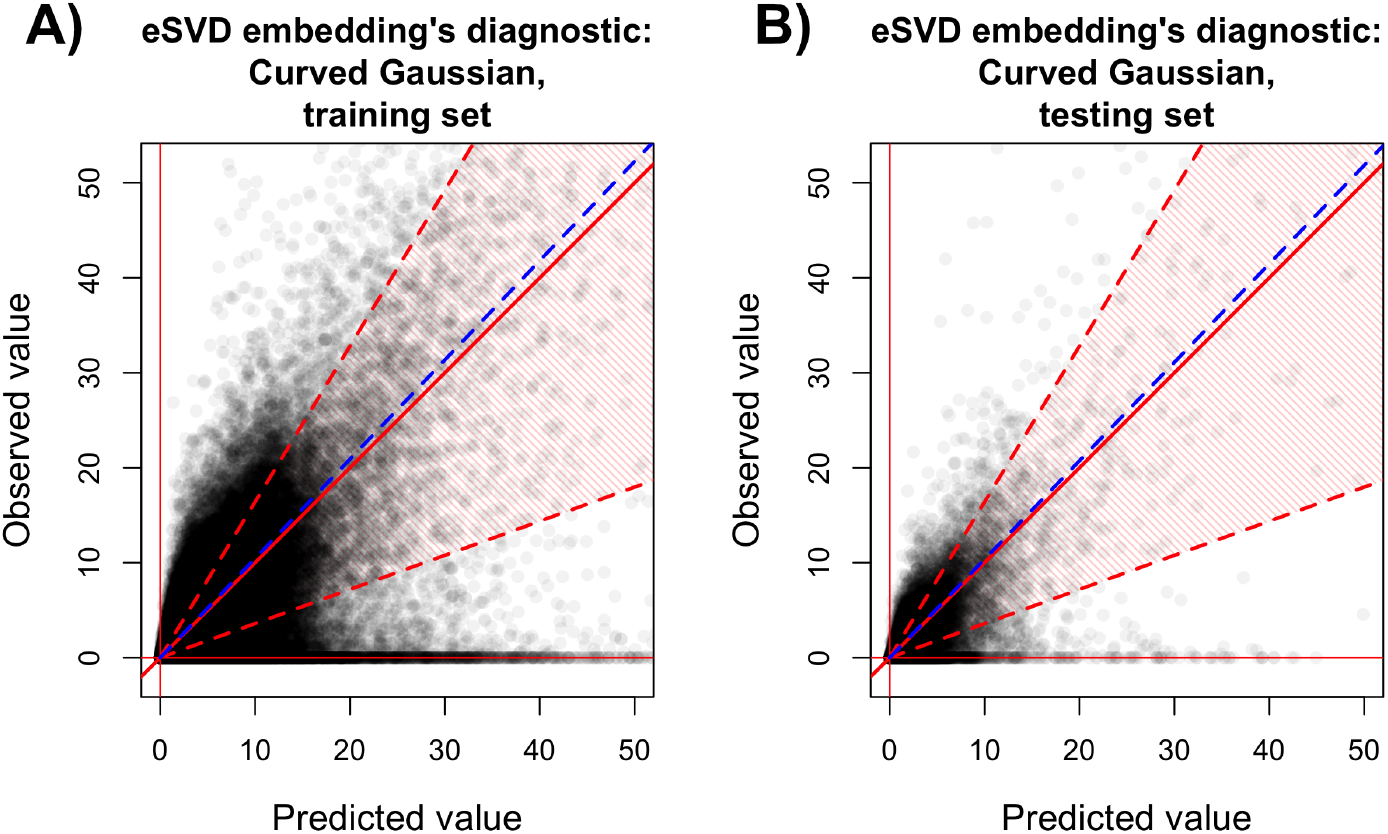
Diagnostic plot showing the same results as in Figure 7, except zoomed in to display values between 0 and 50. (A) The diagnostic plot on the “training set.” (B) The diagnostic plot on the “testing set.”.

## H.2 Additional result for the highly informative genes

To supplement the results in Section 7, we find high informative genes across the different six major cell types which also demonstrate the smooth continuum of gene expression along the two estimated trajectories. This is visualized in Figure 22, where heatmaps of this form are common in work that study the gene expression continuum along trajectories (Marques et al. (2016), Haghverdi et al. (2016) and Bergen et al. (2020)). These heatmaps are sometimes referred to as *cascade plots*. Here, we see that when we order the cells from youngest to oldest based on the estimated pseudotime, there is an ordering of genes such that different genes are highly expressed at different pseudotimes. Additionally, we see that when the two trajectories diverge around the rescaled pseudotime of 0.49, there are genes that are highly expressed for mature oligodendrocytes in one trajectory but not the other.

**Figure 22:**
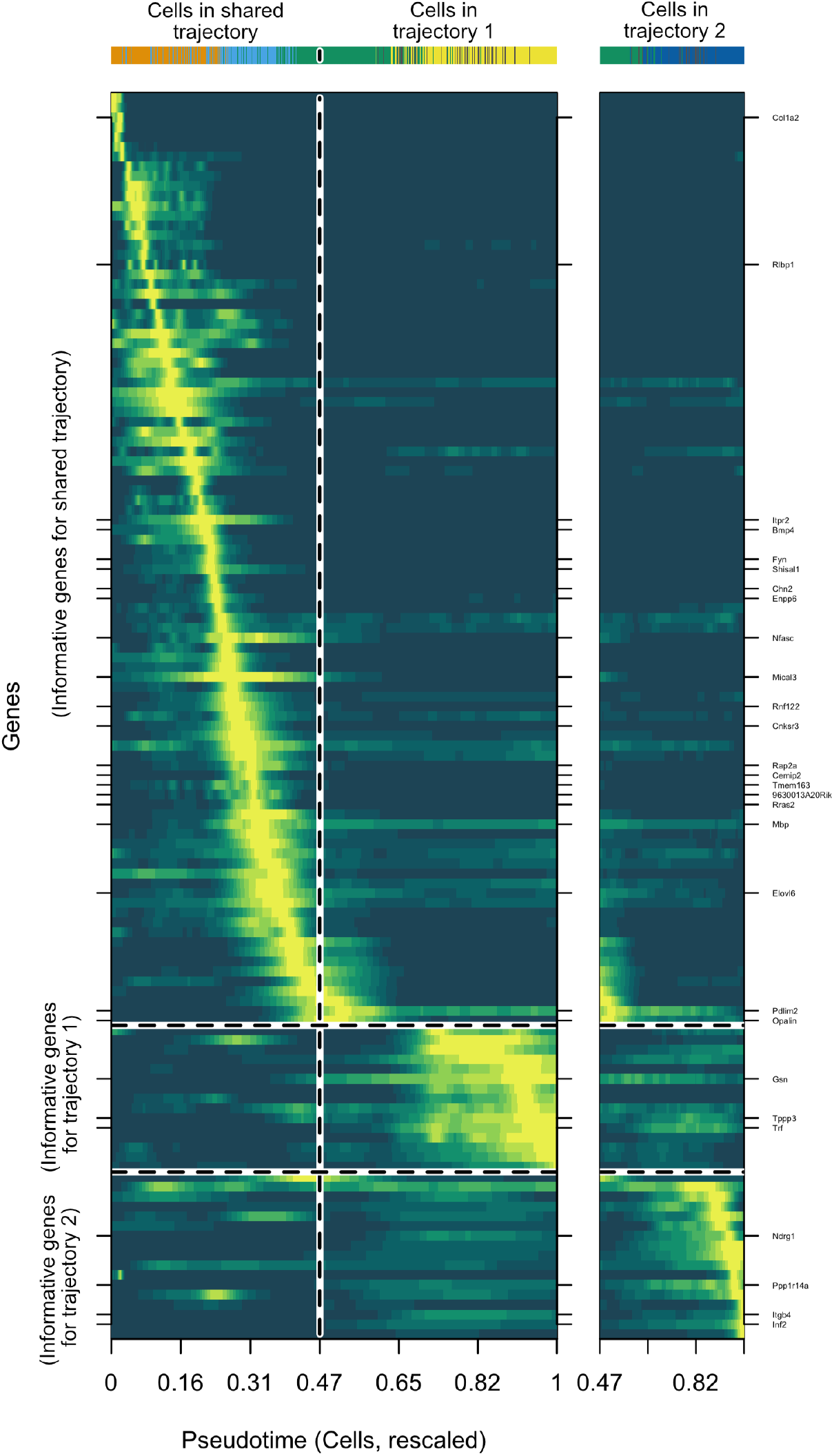
Cascade plot showing the gene expression (rescaled, smoothed) across the cells ordered by the estimated pseudotime. Here, each row denotes a different gene and each column denotes a different cell, where the cell type of each of the corresponding 4449 columns is shown in the top horizontal bar, where the colors correspond to the coloring used in Figure 8. Within this heatmap, yellow denotes high intensity of expression, while dark green denotes low intensity, both relative to all the other cells for a particular gene. The cells prior to a (rescaled) pseudotime of 0.49 are shared between both trajectories, while the cells start to diverge after that pseudotime. The marker genes published in Zhang et al. (2014) for OPCs, NFOLs, and MOLs are explicitly marked in their respective rows for reference.

We briefly describe the procedure to select the genes shown in Figure 22. First, we remove cells that do not easily conform to the estimated trajectories, as by the nature of Slingshot, certain cells belonging to both trajectories might have two distinctly different estimated pseudotimes. We have 4449 cells remaining afterwards, which are used to find highly informative genes from the 983 genes in our analysis. We then order the cells based on their pseudotimes and which trajectories they belong to. This results in two different data matrices, one matrix for the cells shared between both trajectories and the cells unique to the first trajectory, and another matrix for the cells shared between both trajectories and the cells unique to the second trajectory. For each data matrix, we smooth the gene expression across each gene using a univariate kernel nonparametric regression via the function `np::npreg`. Then, for each gene *j*, we use a variant of circular binary segmentation (Olshen et al., 2004) to find a region of cells along either data matrix that represents cells which have high expression for gene *j*. After all this is done, we find the genes that show the highest change in expression (based on circular binary segmentation on the smoothed gene expression) along each value of the pseudotime for both trajectories. This informs us about which genes are informative for the shared trajectory, only the first trajectory, or only the second trajectory. We additionally include marker genes published in Zhang et al. (2014) for reference, which are originally used in the dataset we are investigating Marques et al. (2016).

## H.3 Analysis via eSVD using the negative binomial distribution

For completeness, we include the diagnostic plots on the quality of fit of using the negative binomial distribution to fit the eSVD embedding on the oligodendrocyte data. We tune the negative binomial from a grid of parameters as well, where we choose among the *k* ∈ {3, 5, 10} and dispersion parameter *r* ∈ {500, 2500, 10^4^}. After using our tuning procedure described in Subsection 4.2, we select *k* = 5 and *r* = 10^4^. Analogous to Figure 7, we show the resulting plots on the training and testing set in Figure 23. As we can see, compared to analogous plots for the curved Gaussian distribution (Figure 7), the 10th to 90th quantile of the negative binomial model, denoted by the shaded red polygon, are not wide enough to capture as much of the variability in the data in the testing set.

**Figure 23:**
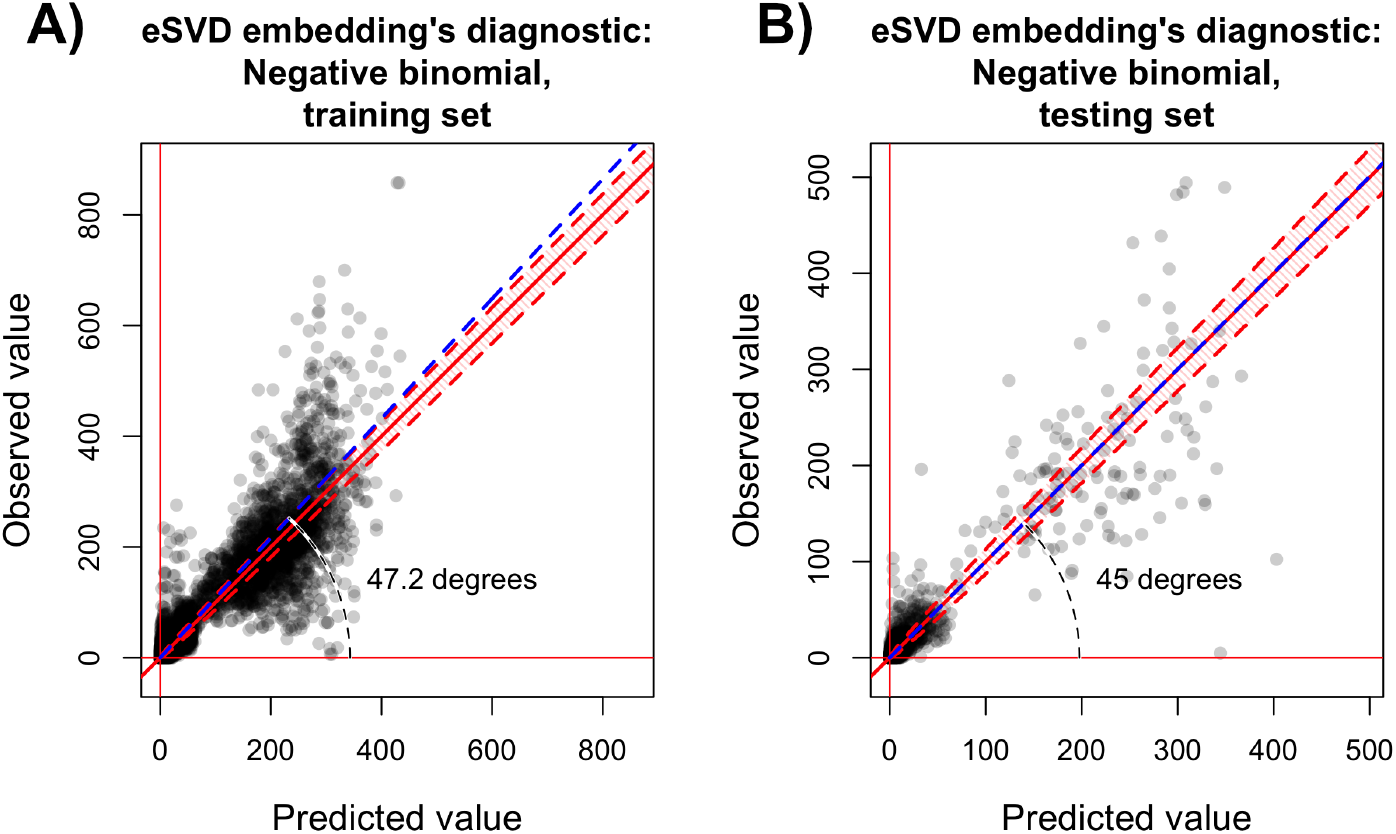
Diagnostic plots analogous to those in Figure 7, except displaying the results of fitting the eSVD embedding with the negative binomial distribution distribution where, after applying the tuning procedure, the number of failures is set to be *r* = 10000 and the latent dimension space is *k* = 5. (A) The diagnostic plot on the “training set.” (B) The diagnostic plot on the “testing set.”.

## H.4 Analysis via UMAP and ZINB-WaVE

As mentioned in the main text, two common embedding methods used in the literature are UMAP (McInnes et al., 2018) and ZINB-WaVE (Risso et al., 2018). For completeness, we present these resulting embeddings derived from these methods for comparison against the eSVD embedding with the curved Gaussian distribution. Our goal here is to convey that using any of these embedding results all qualitatively conclude that the mature oligodendrocytes unique to the first estimated trajectory lie opposite to the mature oligodendrocytes unique to the second estimated trajectory.

In Figure 24, we present the results using the UMAP embedding. Here, we use the `umap R` package, where we set `n_neighbors` to be 30. Analogous to Figure 8, we color the mature oligodendrocytes as gray, yellow or dark blue, depending on whether the cell belongs to the mature oligodendrocyte cell sub-type that is shared between the two estimated trajectories, is exclusively in one of the estimated trajectories, or exclusively in the other estimated trajectory respectively. This UMAP embedding is calculated based on the same dataset used to estimate the eSVD embedding shown in Section 7 (i.e., not log_2_-transformed). Since the UMAP embedding does not rely on a statistical model and does have known statistical consistency, we avoid applying Slingshot on top of this embedding. Nonetheless, we qualitatively see in Figure 24 that the gray points lie between the dark blue and yellow points, suggesting a similar phenomenon to our results in Section 7 where we observe two distinct trajectories that split at the mature oligodendrocytes.

**Figure 24:**
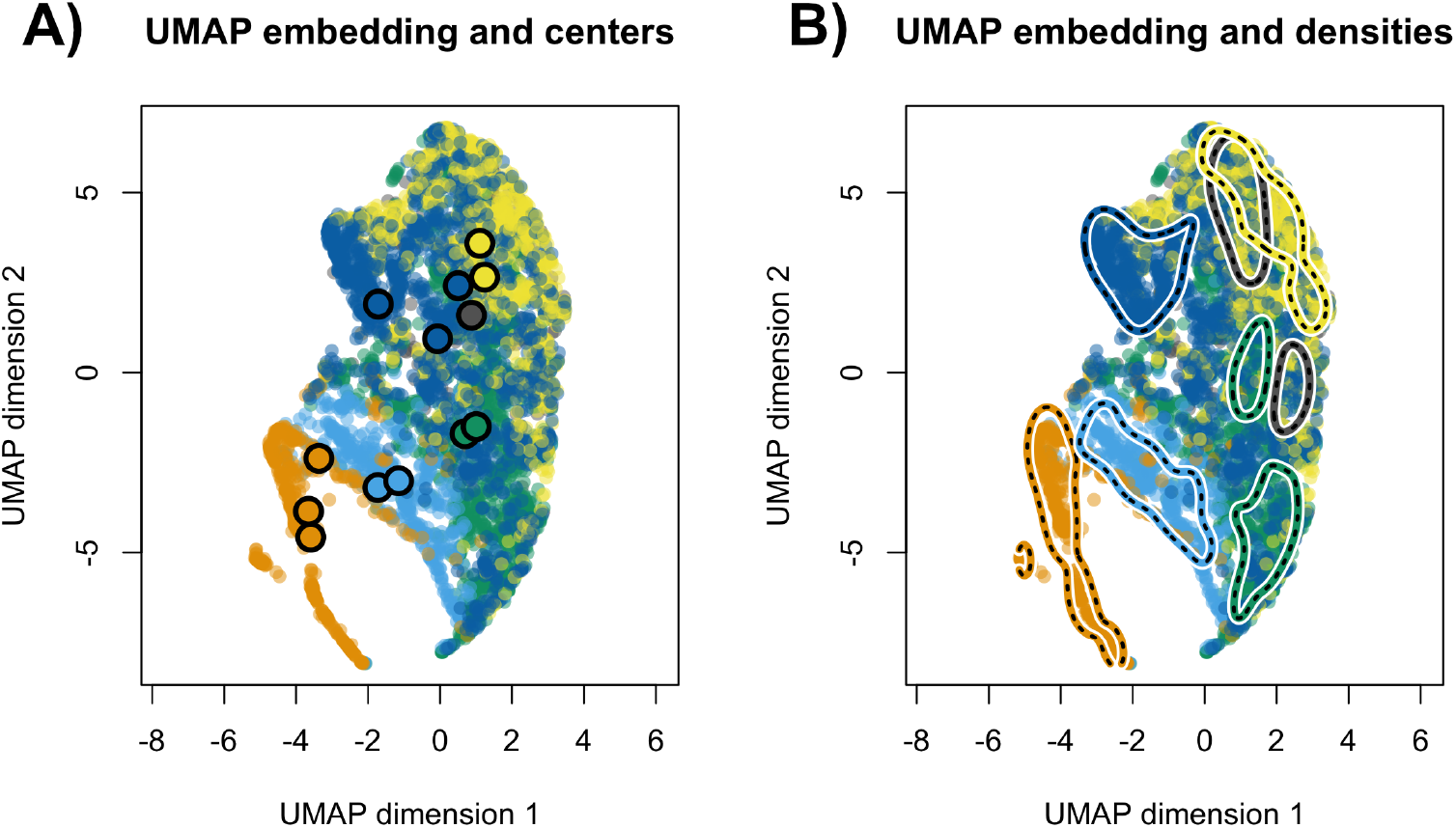
Two-dimensional plot of the UMAP embedding applied to the preprocessed dataset with 5069 cells and 983 genes, (i.e., Steps 1 and 2 as described in Appendix C.1), where the cell types are colored according to Figure 18. (A) UMAP embedding shown alongside the 13 cell cluster centers. (B) UMAP embedding shown alongside with estimated densities to visualize high-density regions (analogous to Figure 19).

Similarly, we present the results using the ZINB-WaVE embedding. This model and its usage is summarized in Appendix F.1. We fit the model using the count data (i.e., *n* × *p* matrix with non-negative integer values) that has undergone only the gene screening step (i.e., preprocessing the data using only Step 1 described in Appendix C.1) and the arguments `K=5, maxiter.optimize=100, normalizedValues=T` and `commondispersion=F`. Similar results in Section 7, the ZINB-WaVE embedding shown in Figure 25 show the gray points lie between the dark blue points and yellow points. This is most visibly seen in the scatter plot between the first two latent dimensions. Hence, these results also suggest a similar phenomenon to our results in Section 7 where we observe two distinct trajectories that split at the mature oligodendrocytes.

**Figure 25:**
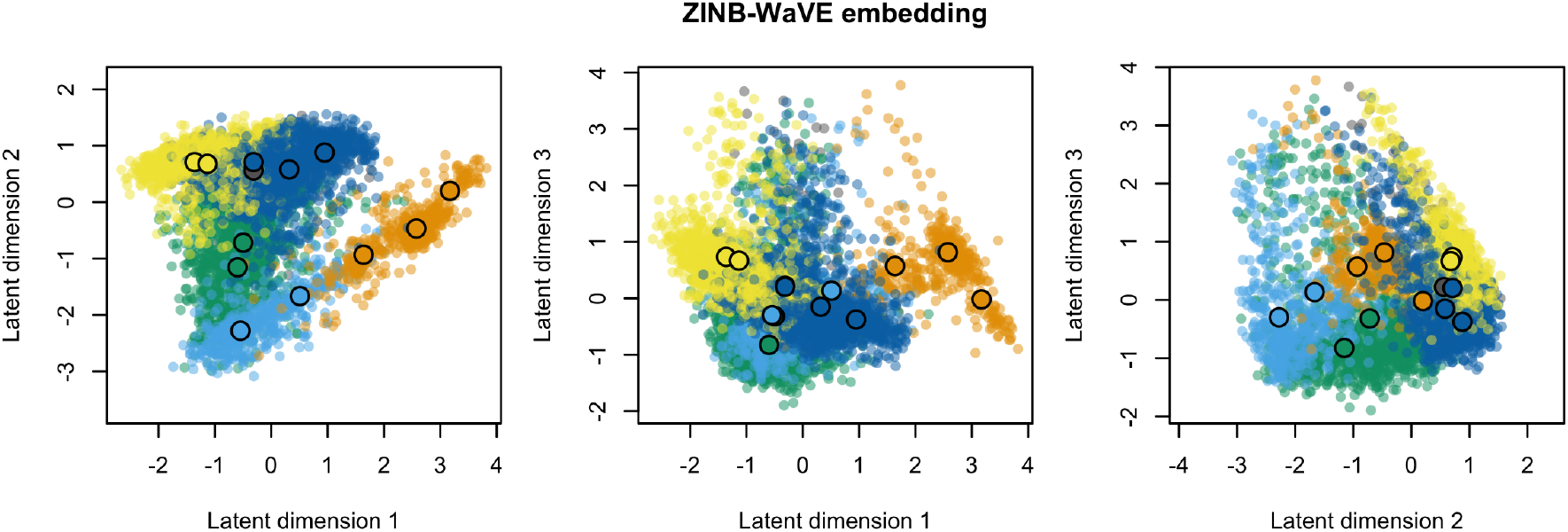
Two-dimensional plots of the ZINB-WaVE embedding applied to the preprocessed count dataset with 5069 cells and 983 genes, (i.e., Step 1 as described in Appendix C.1). The cell types and 13 cell cluster centers are shown to be analogous to those in Figure 18.

## H.5 Additional analysis on a separate dataset

Here, we apply the eSVD to a second dataset to demonstrate its performance on other single-cell datasets outside of the oligodendrocyte dataset studied in the main text. For this, we investigate the eSVD applied to the Zeisel dataset, which is commonly used as a benchmark dataset in other studies (Lun et al., 2016; Zappia et al., 2017; McCarthy et al., 2017; Kiselev et al., 2017; Li and Li, 2018). This dataset studies cells in mice’s nervous systems, and contains 3005 cells across seven different primary cell types (astrocytes ependymal, endothelial mural, interneurons, microglia, oligodendrocytes, pyramidal CA1, and pyramidal SS), of which the least represented cell type has 98 cells. The goal in this particular analysis is to determine if cells of the same cell type are “well-clustered” after embedding each cell into the latent space.

Our analysis strategy proceeds similarly to the one described in Appendix B. Specifically, we first screen the cells in a similar fashion, with exception that now we normalize each cell by its library size prior to screening for genes. After screening, there are 1628 genes remaining. Then, we compare two analyses, comparable to those described in the main text. In the first analysis, we log_2_-transform all the entries of the preprocessed single-cell data and then embed each cell via the SVD. In the second analysis, we do not log_2_-transform the entries, but instead rescale the data uniformly so the maximum value is 1000 and embed each cell via the eSVD with a negative binomial distribution. Our choice of analyzing the Zeisel dataset via the negative binomial distribution here is to demonstrate the eSVD’s versatility in how it can be usefully applied to one-parameter exponential-family distributions other than the curved Gaussian distribution used in Section 7, and we also empirically observe that this data has less noise compared to the oligodendrocyte dataset studied in the main text. We tune the latent dimensionality of both embeddings (among *k* ∈ {10, 20, 50}) and the nuisance parameter for the negative binomial distribution (among *r* ∈ {250, 500, 1000, 10000, 50000}) via the tuning procedure described in Subsection 4.2. For the SVD embedding, this resulted in selecting *k* = 50, and for the eSVD embedding, this resulted in selecting *k* = 20 and *r* = 10000.

To assess how well the cells are clustered within each embedding, we derive a “purity” statistic based on the the shortest path between cells of the same cell type in the nearest-neighbor graph. We do not use statistics based on K-means clustering or other clustering methods since we wish to not confound the quality of the embedding with the appropriateness of the clustering method as much as possible. To describe this statistic, first construct the nearest-neighbor graph by representing each of the 3005 cells as a vertex, and connecting each vertex to its *m* nearest neighbors (for example, *m* = 3). Then, let cell *i* and *j* be of the same cell type *t* (for example, interneurons), and consider the shortest path between vertices *i* and *j*. We measure the purity statistic between vertices *i* and *j* by the percentage of vertices in this shortest path that correspond to cells of the same cell type *t*, excluding cells *i* and *j* themselves. This statistic is in [0, 1] and a larger value is more desirable, as it suggests that more cells of the same cell type are “between” cells *i* and *j*. If there are multiple shortest paths, we pick the one that maximizes this statistic. For each of the seven different cell types, wee sample 5000 random pairs of cells *i* and *j* to measure this purity statistic. This results in 35000 values that describe the quality of the embedding.

Using the purity statistic, we see that the eSVD embedding (without the log_2_-transform) preserves the clustering structure among the seven cell types better than the SVD embedding (with the log_2_-transform). Specifically, if we were to construct a nearest-neighbor graph based on *m* = 3, then the average (mean) purity statistics across all 35000 values for the eSVD embedding and the SVD embedding are 0.93 and 0.91 respectively. However, this statistic is already equal to 1 at the 25th quantile among the 35000 values for either embedding. When we focus on values between the 0th and 25th quantile only, the average (mean) purity statistic for the eSVD embedding and the SVD embedding are 0.73 and 0.65 respectively. When we try different values of *m* from *m* = 3 to *m* = 5, we observe a similar result regardless of how we construct the nearest-neighbor graph (Figure 26). This suggests that our analysis results are robust to the value of *m*. Choosing *m* to be smaller than 3 results in a nearest-neighbor graph that is not connected.

**Figure 26:**
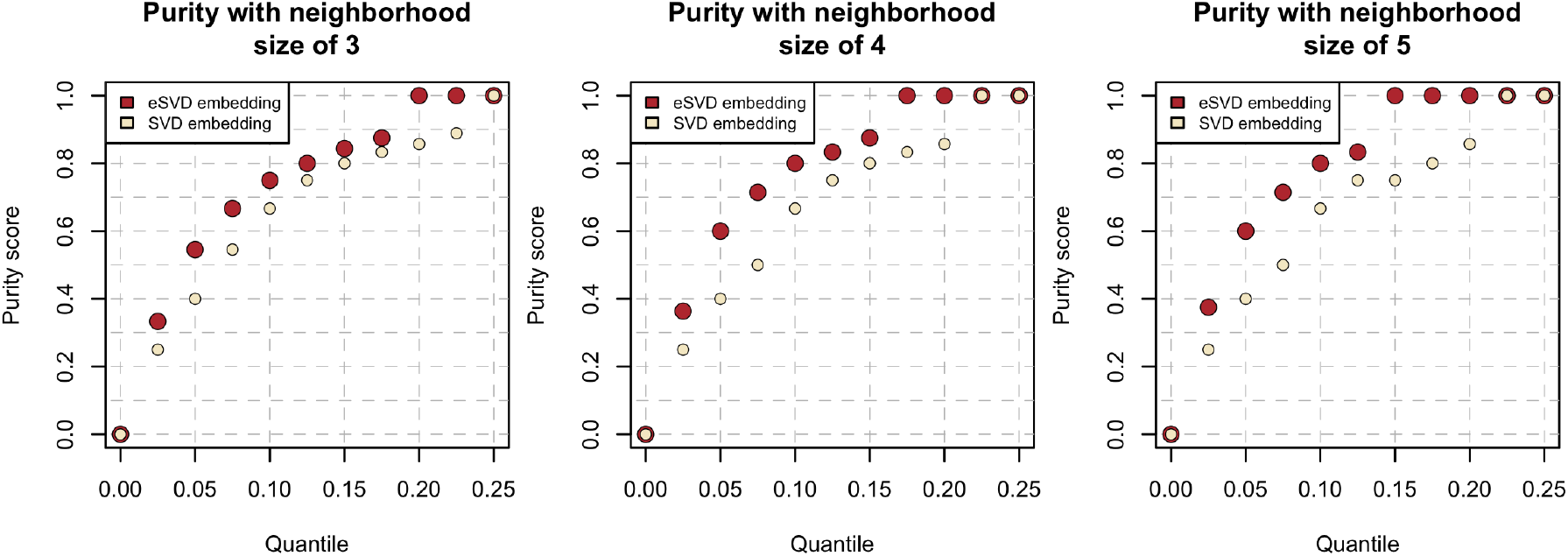
Quantiles of the purity score among the 35000 values for both embeddings when applied to the Zeisel dataset, from the 0th to 25th quantile, across *m* = 3 to *m* = 5.

## I Proofs

In Appendix I.1, we prove Proposition 5.1. In Appendix I.2, we prove Proposition 5.2 and its related lemmas. In Appendix I.3, we prove Proposition D.1 and its related lemmas. In Appendix I.4, we prove the results shown in Appendix E.

Throughout these proofs, for generic matrices *A* and *B*, let ||*A*||_op_ denote the spectral norm, i.e., the largest singular value. If *A* is a square matrix, let tr(*A*) denote the trace of *A*, i.e., the sum of its diagonal elements. We write *A* ⊗ *B* to denote the Kronecker product of *A* and *B*, and if *A* and *B* are of the same dimensions, we write *A* ⪯ *B* if *B* − *A* is positive semidefinite. For two deterministic sequences *a*_*n*_ and *b*_*n*_, let *a*_*n*_ = *o*(*b*_*n*_) denote that *a*_*n*_/*b*_*n*_ → 0. For two random sequences *A*_*n*_ and *B*_*n*_, let *A*_*n*_ = Ω_*p*_(*B*_*n*_) denote *B*_*n*_/*A*_*n*_ is bounded in probability for large enough *n*. Also, let *I*_*k*_ denote the identity matrix of size *k* × *k* and **1**_*n*×*p*_ represent the *n* × *p* matrix of all ones. Throughout these proofs, we implicitly use different equivalent definitions of strong convexity and functions with Lipschitz gradients as stated in Zhou (2018).

## I.1 Proof for Proposition 5.1

## Proof of Proposition 5.1

Define the eigendecompositions of the second moment matrices for all *i* ∈ {1,…, *n*} and *j* ∈ {1,…, *p*},

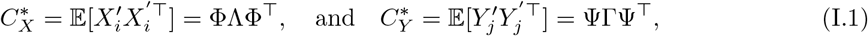

where Φ and Ψ are both *k* × *k* unitary matrices. Our construction of the invertible matrix *R* ∈ ℝ^*k*×*k*^ will be done in two steps. In the first step, we first construct an invertible matrix 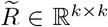 such that

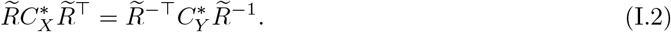

This would yield a transformation matrix to ensure 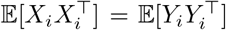. In the second step, we adjust 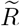 into the desired matrix *R* in order to ensure that both 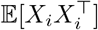 and 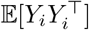 are diagonal.

**Step 1:** Based on the definition (I.1) and the desired goal shown in (I.2), an equivalent goal is to show that

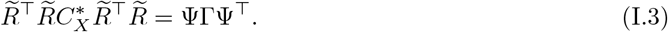

Let *Q* ∈ ℝ^*k*×*k*^ denote any unitary matrix, to be specified later. Since the matrices on both sides of the above display are symmetric, in order to show (I.3), we can instead we can find an *Q* and 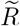such that,

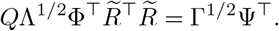

Rearranging the above display, we have

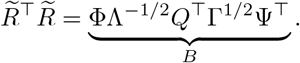

Hence, we are done once we construct a unitary matrix *Q* that makes *B* symmetric. Observe that if the matrix

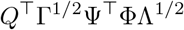

were symmetric, then *B* would be symmetric. (This can be seen by multiplying the above matrix on the left by *E* = ΦΛ^−1/2^ and on the right by *E*^⊤^) Observe that Γ^1/2^Ψ^⊤^ Λ^−1/2^Φ is guaranteed to be full rank (by assumption of 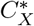 and 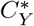 being full rank), so Γ^1/2^Ψ^⊤^ Λ^−1/2^Φ admits a rank-*k* SVD of *UDV*^⊤^. Since the product of two unitary matrices is still unitary, we set *Q* = *UV*^⊤^. Hence, we finished our construction of 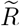.

**Step 2:** Suppose 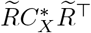 (or equivalently, 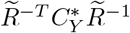 based on (I.2)) has eigenvectors *W*_1_,…, *W*_*k*_. Let *W* be the unitary matrix formed by concatenating these *k* eigenvectors column-wise. By diagonalization, we know 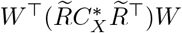 is diagonal. This implies that our final construction is 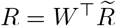.

## I.2 Proof for Proposition 5.2

We first introduce some notation. Let the SVD of *X* and *Y* be denoted as

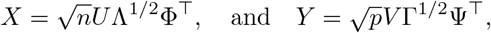

Using this definition, the empirical second moment matrices are

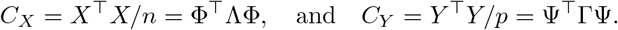

Prior to the proof, we offer a high-level description of the proof strategy. Similar to the proof in Lei (2018), we introduce two levels of approximation. Let us focus on estimating 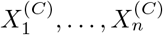, the latent variables that are oriented based on the population covariance matrix 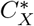. Formally,

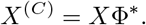

(Recall by Assumption 5.2, since *C*^∗^*X* is diagonal, the columns of Φ^∗^ are the standard basis vectors.) To approximate this, we consider 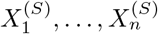, the latent variables rotated by their own right singular vectors,

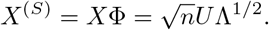

This approximation is driven by *C*_*X*_ being close to 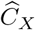. This is in turn estimated by 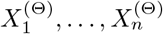 based on the SVD of Θ,

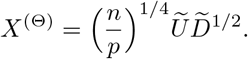

This approximation is driven by 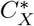 being equal to 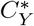. Finally, this is approximated by a quantity we can actually compute from data, our estimates 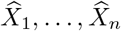 as in (4.5),

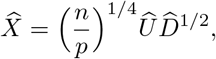

where 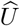 and 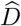 are obtained by an SVD of our estimate 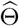 (recalling that we denote the SVD of 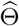 as 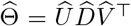). This approximation is driven by 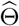 being close to Θ (i.e., our assumption that 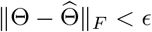.

## Proof of Proposition 5.2

**Step 1:** (Decomposition of error). By the triangle inequality, the rate is dictated by the term,

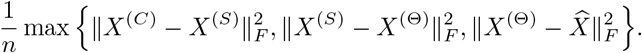

In the following three steps, we bound each term individually, of which the maximum of all three terms concludes the proof.

**Step 2:** (Approximation from *X*^(*C*)^ to *X*^(*S*)^). First, we deduce the relations of the eigenvalue λ_*j*_ and eigenvectors Φ_*j*_. By applying Lemma J.1, we obtain

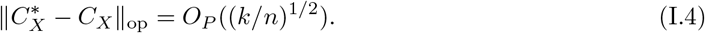

By using Weyl’s inequality and the Davis-Kahan theorem along with Assumption 5.2, this immediately implies

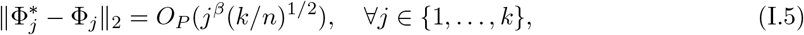

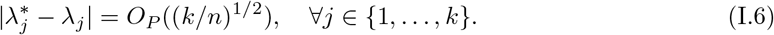

Since *k* = *o*(min{*n*, *p*}) by assumption, this implies

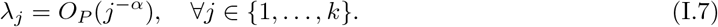

Thus, using the trace function,

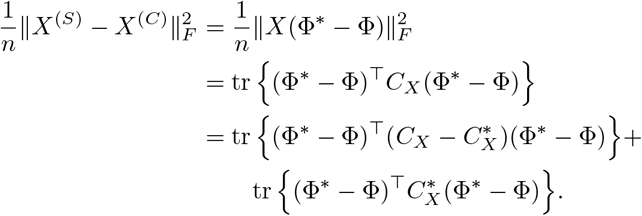

For the first term,

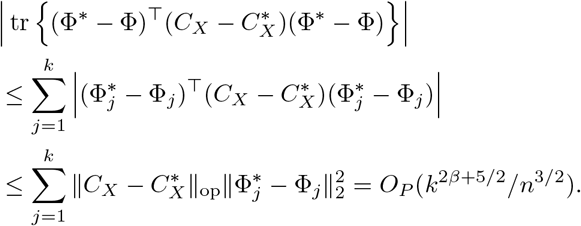

For the second term, recalling that *α* ≤ *β* and Parseval’s identity,

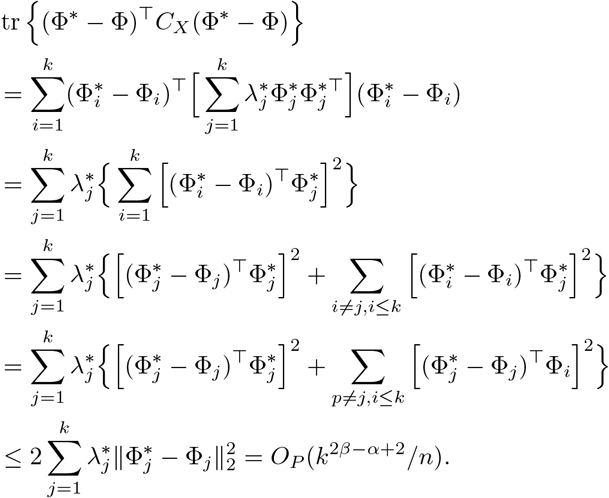

**Step 3:** (Approximation from *X*^(*S*)^ to *X*^(Θ)^). First, observe that 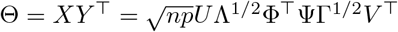, and we would like show it is close to the matrix 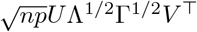, which has an an SVD of 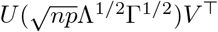. Specifically,

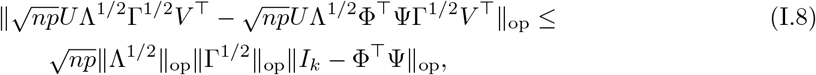

where we can bound the last term by using the submultiplicative property of the spectral norm and 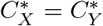,

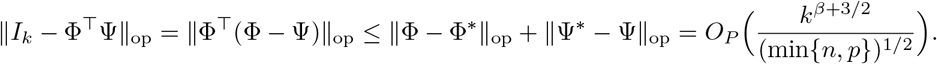

where the last inequality is due to Davis-Kahan. Hence, plugging the above display into (I.8),

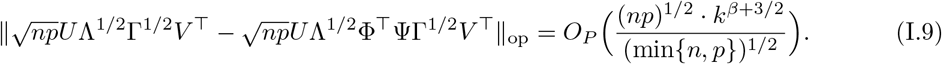

First, we derive the difference in eigenvalues based on (I.9). We apply Weyl’s inequality to the above display to conclude

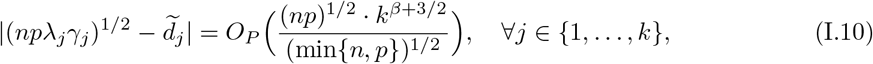

recalling that 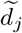 is the *j*th singular value of Θ. Note from (I.6) that we also have for all *j* ∈ {1,…, *k*},

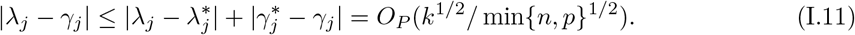

Hence, combining (I.10) and (I.11), noting that max 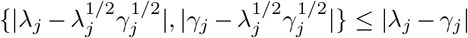, we derive

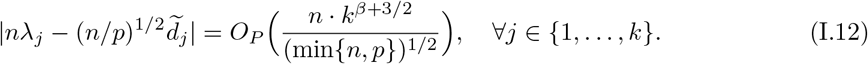

Combining the above display with Assumption 5.2 and (I.7) and given *k* = *o*(min{*n*, *p*}), this implies

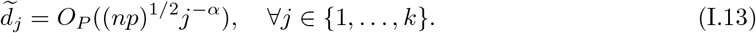

Second, we derive the difference in eigenvectors based on (I.9). Observe that based on (I.6) and *k* = *o*(min{*n*, *p*}), we can derive that 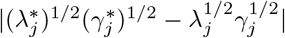 is dominated by Ω_*P*_(*j*^−*β*^). Hence, we can derive the spacing of the singular values of 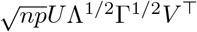,

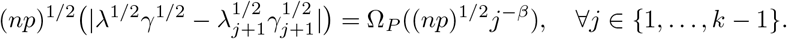

Hence, using the Davis-Kahan theorem by combining (I.9) with the above display, we conclude

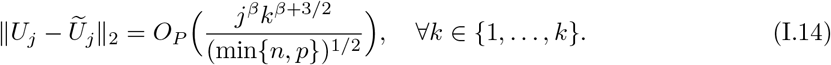

Using (I.7), (I.12), and (I.14), along with Lemma J.3,

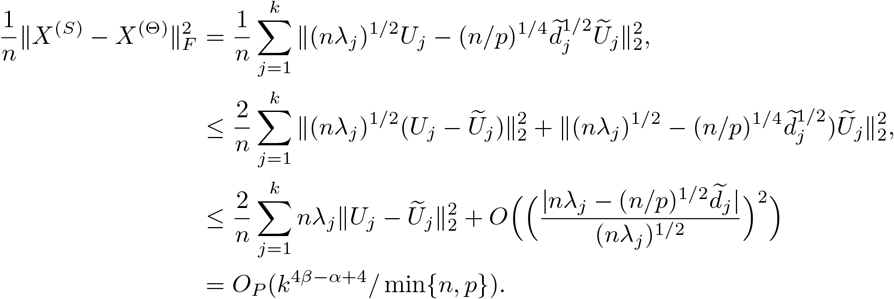

**Step 4:** (Approximation from *X*^(Θ)^ to 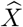). By assumption, we have

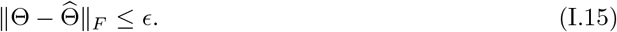

This implies 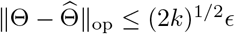. By Weyl’s inequality, we conclude

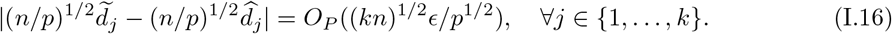

In addition, following a similar logic as above, using the Davis-Kahan theorem along with (I.12) to control the spacing of the eigenvalues, we can derive

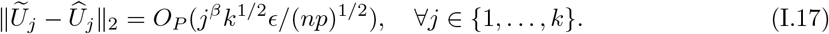

Hence, analogous to the derivation above, using (I.13), (I.16), and (I.17), along with Lemma J.3,

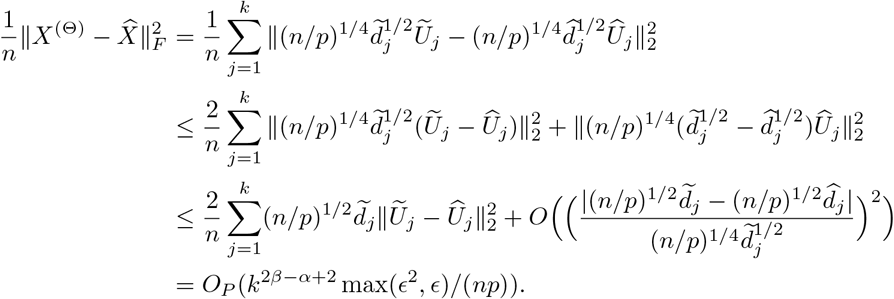

## I.3 Proofs for Proposition D.1

Similar to Balakrishnan et al. (2017), we first analyze the behavior of an iteration of eSVD when using the population loss function 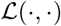. Then, by leveraging Assumption D.3, we can analyze the behavior of eSVD when applied on the sample loss function 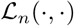.

### Lemma I.1.

*Assume the population loss function* 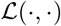 *satisfies Assumptions D.1 and D.2. Conditioned on X and Y, then for any iteration t* ∈ {1,…, *T*}, *there exists matrices* 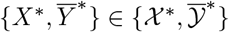 *and* 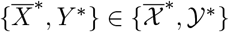 *such that*

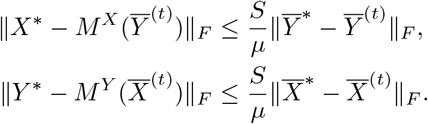

*Proof*. Observe that by the first-order optimality conditions, for any 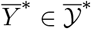, we have

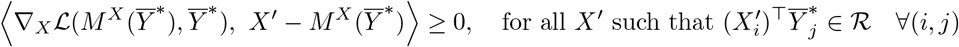

and similarly,

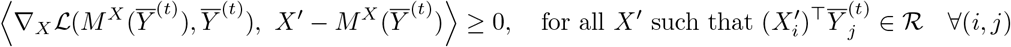

By combining the two inequalities by setting 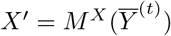 in the first display and 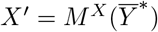 in the second display with some algebra, we get

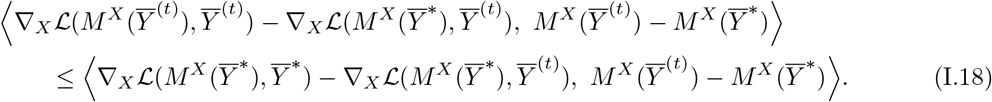

By manipulating the properties of strong convexity assumed in Assumption D.1, we can lower bound the left-hand term in (I.18) by

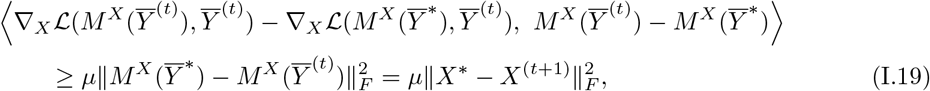

where we plugged in the definition of 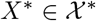 and *X*^(*t*+1)^ in the last display (see (D.5)).

Similarly, by the Lipschitz smoothness assumed in Assumption D.1 with Cauchy-Schwarz inequality, we can upper bound the right-hand term in (I.18) by

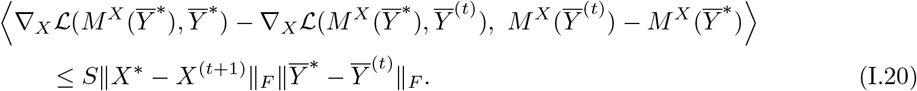

Combining (I.19) and (I.20) into the original equation (I.18) yields

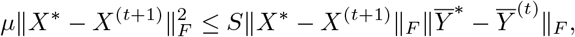

which completes the proof after simple rearrangements.

Similar to Zhao et al. (2015), we now prove Proposition D.1 using the above lemma in conjunction with Lemma J.2 which describes the effect of the LeftSVD operator.

## Proof of Proposition D.1

For simplicity, let *κ* = *S*/*μ*. By triangle inequality and Lemma I.1, we have that there exists some 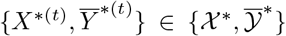 and 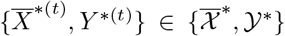 for all *t* ∈ {0,…, *T*},

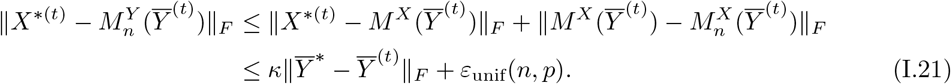

Applying Lemma J.2 and recalling that the smallest non-zero singular value of *X*^^∗^(*t*)^ is 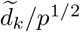, we obtain

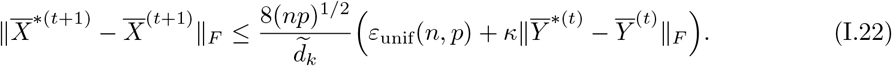

Similarly,

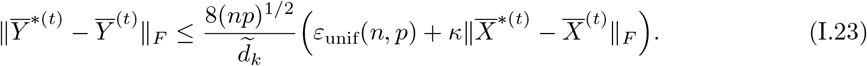

Hence, observe that by the conditions of Proposition D.1, we are ensured that if 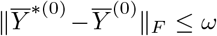 (Assumption D.4), then by iteratively applying (I.21) through (I.23), then max 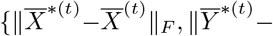 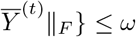 for all *t* ∈ {1,…, *T*}. Therefore, by infinite geometric summation,

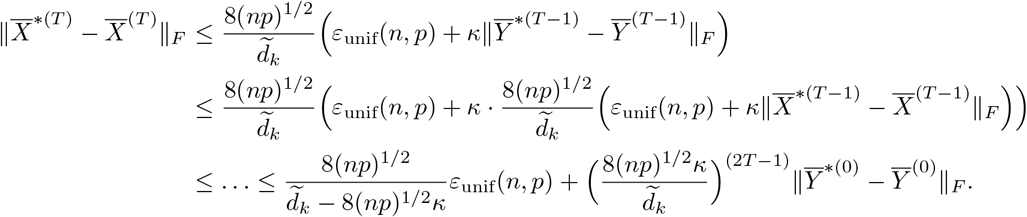

Once again, by the conditions of Proposition D.1, we are ensured that

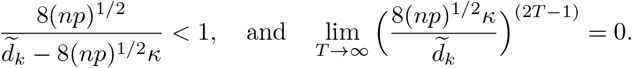

Therefore, for large enough *T*, we conclude for some universal constant *C*,

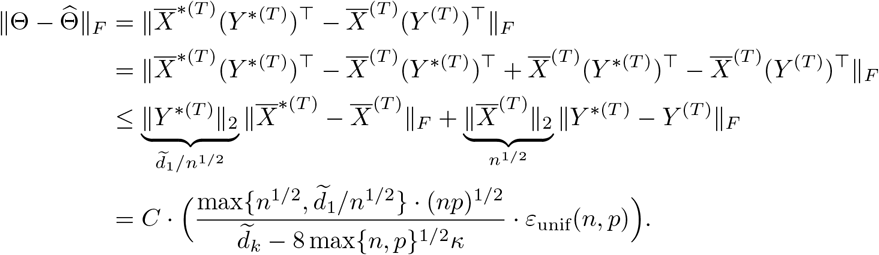

## I.4 Proofs for Lemma E.1 through Corollary E.5

## Useful facts

It is useful to have the following forms of the gradients and Heissan written down.

- The gradient 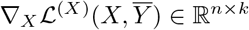 has the *i*th row equal to, for *i* ∈ {1,…, *n*},

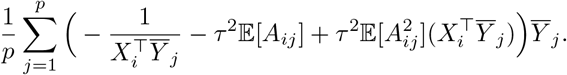
- Let 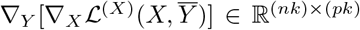 denote the gradient of above function with respect to 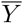. If we focus on a particular block of *k* rows corresponding to a specific *X*_*i*_ and a particular block of *k* columns corresponding to a specific 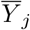, we have

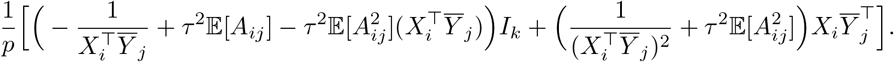
- The Hessian matrix 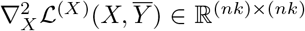 is all 0, except for *n k* × *k* blocks along the diagonal. The *i*th block is equal to, for *i* ∈ {1,…, *n*},

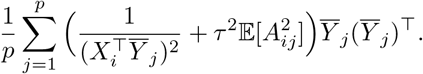 We list properties about the curved Gaussian distribution assumed in (3.1) that will be needed. Recall that 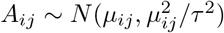, where *μ*_*ij*_ = −1/*θ*_*ij*_.
- (First and second moment):

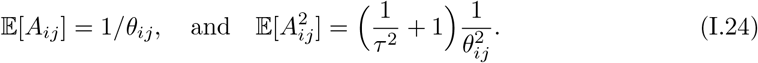
- (Squared random variable):

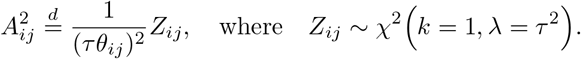

Hence, the variance of 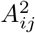 is upper-bounded by

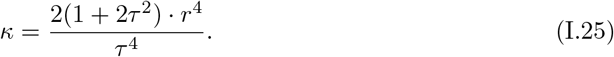

## Proof of Lemma E.1

We start with the lower bound. Observe that minimum eigenvalue of one of the *n k* × *k* blocks will be the minimum eigenvalue of the overall Hessian matrix 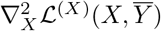. Hence, inspecting one particular block for a specific *i* ∈ {1,…, *n*}, observe that

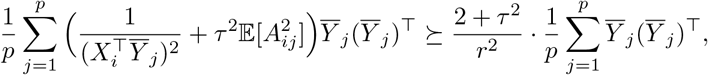

where 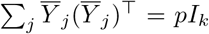 since the columns of 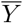 are orthogonal. For the upper bound, we apply the same logic.

## Proof of Lemma E.2

Recall by the multivariate Taylor expansion (Feng et al., 2013), we have

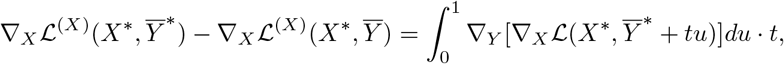

where 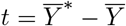. This leads to the following inequality,

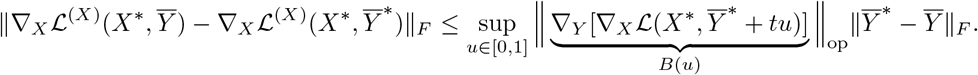

Observe we can split the matrix *B*(*u*) into two matrices, one sparse matrix and one dense matrix. That is, *B*(*u*) = *B*_1_(*u*) + *B*_2_(*u*), all matrices of dimension (*nk*) × (*pk*). The (*i*, *j*)th *k* × *k* block of *B*_1_(*u*) has entries equal to

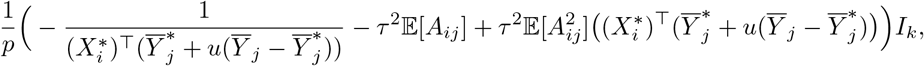

while the (*i*, *j*)th *k* × *k* block of *B*_2_(*u*) has entries equal to

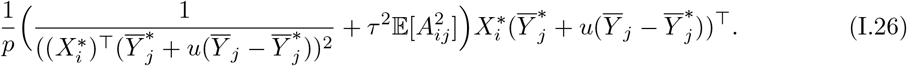

We first analyze *B*_1_(*u*). Plugging in (I.24), we obtain

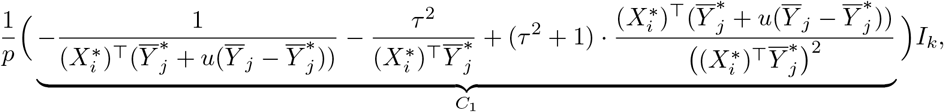

To analyze *C*_1_ in the perspective of Assumption E.2, we analyze

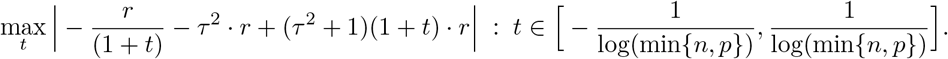

By analyzing the Lagrangian of the above display, we see that for min{*n*, *p*} large enough,

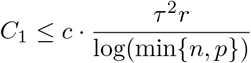

for some universal constant *c*. Hence, we derive that

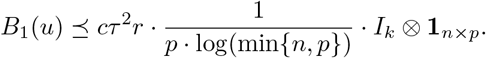

Therefore,

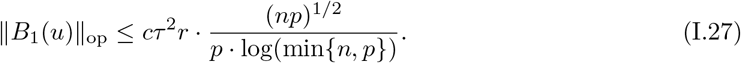

We next analyze *B*_2_(*u*). Plugging in (I.24), we obtain

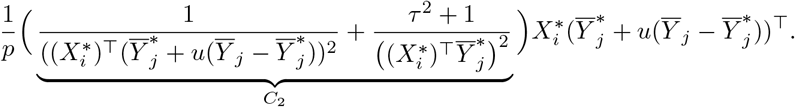

By analyzing *C*_2_ similar to how we analyzed *C*_1_, we derive that for min{*n*, *p*} large enough,

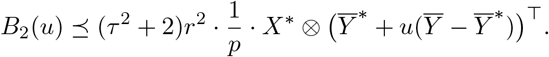

Therefore,

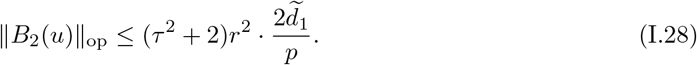

We conclude by combining (I.27) and (I.28) by a triangle inequality.

## Proof of Lemma E.3

Prior to proving Lemma E.3, we need the following set of concentration bounds. Recall that κ upper-bounds the variance of 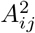 under the curved Gaussian model (see (I.25)).

### Lemma I.2.

*Conditioned on* Θ, *let A*_*ij*_ *follow the distribution* (4.7). *Let κ* = (2(1 + 2τ^2^))*r*^4^/τ^4^. *Under Assumption E.1, for a fixed i* ∈ {1,…, *n*}, *for some universal constant c, for fixed matrices X and* 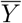, *each of the following four inequalities hold separately*.

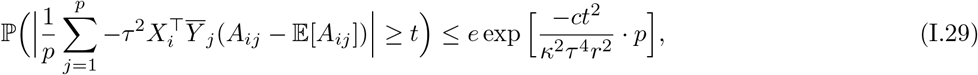

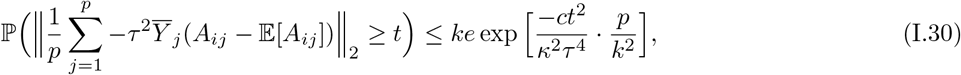

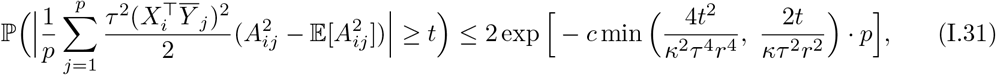

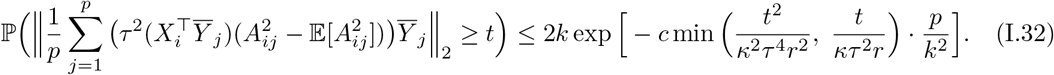

*Proof*. (I.29) and (I.31) are direct applications of Hoeffding’s and Bernstein’s inequality (Vershynin (2010), Proposition 5.10 and 5.16) since 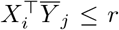. To show (I.30), observe that for a particular coordinate ℓ = 1,…, *k*, we can apply Hoeffding’s inequality to show

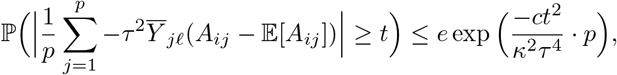

since the ℓ_2_-norm of any column of 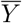 is 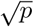. Applying a maximal inequality and then summing over all *k* dimensions, we have

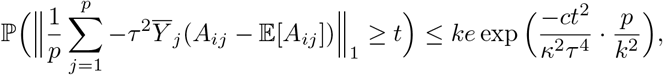

which upper-bounds the RHS of (I.30). The same technique using Bernstein’s inequality can be used to show (I.32).

With these concentration statements, we are ready to proceed with the analysis of ε_fixed_(*n*, *p*) row-wise.

### Lemma I.3.

*Assume the conditions in Lemma E.1. Let*

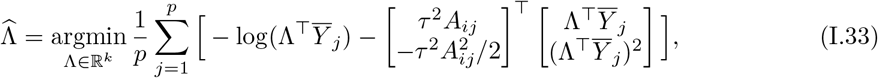

*and*

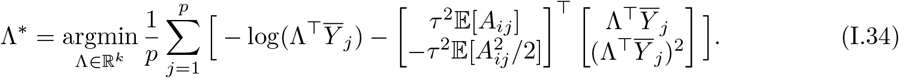

*Then, conditioned on X and Y, with probability at least* 1 − 6/(*np*),

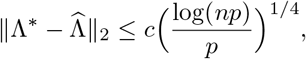

*where c is a constant that depends only on k, μ, L, τ and r*.

*Proof*. **Step 1:** (Setup). Our proof is inspired by Lemma 4 of Candès et al. (2020). The proof is fairly straightforward and relies mainly on the convexity properties 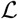 of and the concentration statements established in Lemma I.2.

Throughout this proof, we let *c*_1_, *c*_2_,… denote constants depend on quantities that we will explicitly state, but their explicit form can change from line to line. Let 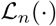 and 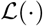 denote functions being minimized in (I.33) and (I.34) respectively throughout this proof only. Since 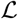 is *μ*-strongly convex, we have

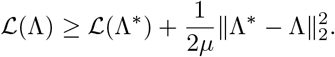

Define 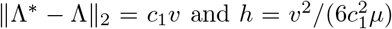 for some constant *c*_1_ that depends on quantity defined later in the proof. Using these definitions, the above display equals

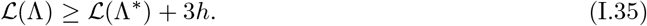

Next, consider the sphere centered at Λ^∗^ with radius *v*, 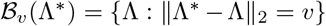. Consider the event *E* defined as

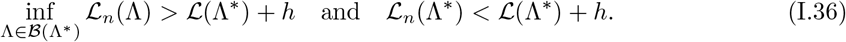

Note that when event *E* occurs, by convexity of 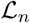, 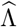 (the minimizer of 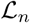) must lie within the sphere 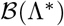, so hence, 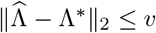.

**Step 2:** (Decomposition of *E*). Let Λ_1_,…, Λ_*M*_ be *M* points on 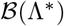, to be specified in the next step. We decompose *E* in the above display into three separate events that we will control individually. Let the event *E*_1_ be defined as

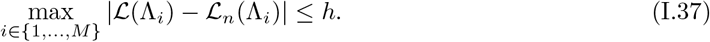

and the event *E*_2_ be defined as

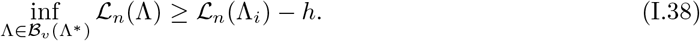

Starting from the event *E*_1_, we have the following line of implications

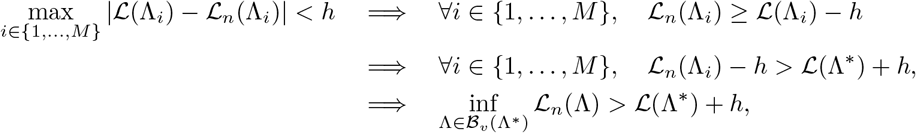

where the second implication follows from the definition of *h* in (I.35), and the last implication holds under the event *E*_2_. Hence, together *E*_1_ and *E*_2_ would imply the first part of *E* in (I.36).

Observe that the following event *E*_3_, defined as,

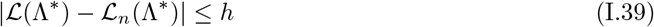

would imply the second part of *E* in (I.36). Hence, since the event *E*_1_⋂*E*_2_⋂*E*_3_ implies event *E*, we have by union bound

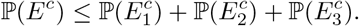

Therefore, the remainder of the proof shows that for a rate of *v*, the summation of the probability of the complementary events is small.

**Step 3:** (Analysis of *E*_1_). We can construct an γ-net of the sphere 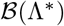. Specifically, by Lemma 9.5 of Ledoux and Talagrand (2013), we require at most most *M* = (3/γ)^*k*^ matrices, denoted as Λ_1_,…, Λ_*M*_, that satisfies,

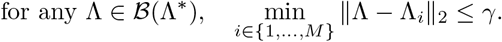

For the rest of the proof, we will set

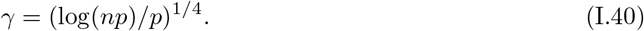

We analyze *E*_1_ in (I.37), the event that 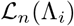 and 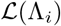 are close along the γ-net. Using a union bound along with concentration bounds shown in Lemma I.2, for *p* large enough, we have with probability at least 1 − 2/(*np*) that event *E*_1_ occurs,

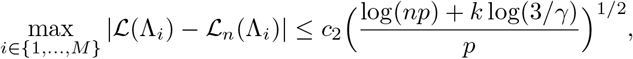

where *c*_2_ depends on *τ* and *r*. Observe that with our choice of γ in (I.40), log(*np*) dominates *k* log(3/γ) for *n* large enough. Hence, treating *k* as a constant, to set the left-hand side of the above display to be equal to *h*, we need

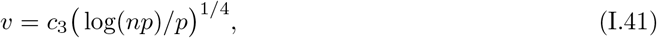

where *c*_3_ depends on *k*, *μ*, τ, and *r*.

**Step 4:** (Analysis of *E*_2_). Next, we analyze *E*_2_ in (I.38). By convexity, letting Λ_*i*_ be the closest vector to Λ in ℓ_2_ distance,

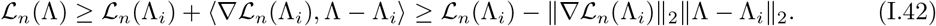

We know ||Λ − Λ_*i*_||_2_ ≤ γ by construction. To bound 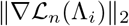, using a union bound and Lemma I.2 once again, we have with probability at least 1 − 2/(*np*) for *p* large enough, event 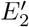 occurs,

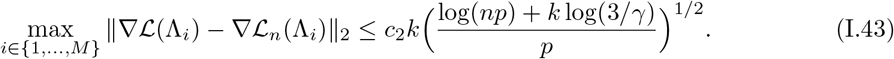

Furthermore, we know that since 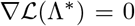 and the gradient of 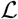 is *L*-Lipschitz smooth, using the definition of *v* in (I.41), we have

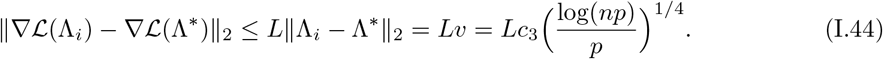

Therefore, combining (I.43) and (I.44), that on event 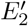,

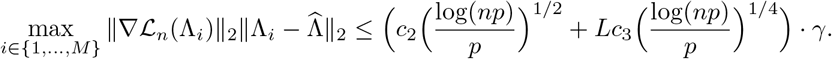

Recalling the rate of γ set in (I.40) and plugging the above inequality back into (I.42), we have on event *E*_2_,

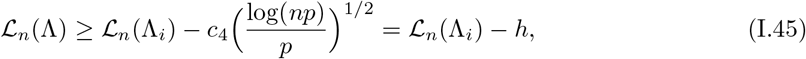

where *c*_4_ depends on *L*, *k*, *μ*, τ, and *r*. This shows *E*_2_.

**Step 5:** (Analysis of *E*_3_ and conclusion). From similar calculations above based on Lemma I.2, we can show that for *p* large enough, we have with probability at least 1 − 2/(*np*) that event *E*_3_ occurs,

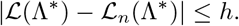

Hence, we conclude the proof, where *c*_1_ initially stated is a constant that depends on *L*, *k*, *μ*, τ, and *r*. By Lemma E.1, we know *L* and *μ* are constants that depend on only τ and *r*.

We are now ready to prove Lemma E.3.

## Proof of Lemma E.3

Let 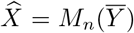 and 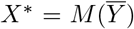 By Lemma I.3, we have with probability 6/(*np*), for any *i* ∈ {1,…, *n*},

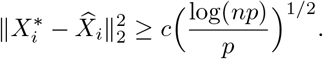

Hence, applying a union bound over all *i* ∈ {1,…, *n*}, we have with probability 6/*p*,

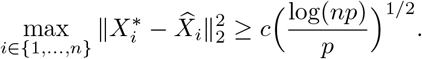

This event implies that,

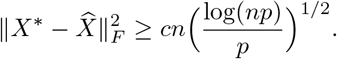

## Proof of Corollary E.4

The proof mainly follows from the proof of Proposition D.1 above, where we replace the term ε_unif_(*n*, *p*) with ε_fixed_(*n*, *p*), which requires us to take a union bound over all *T* iterations.

Due to our invocation of Assumptions 5.1 and 5.2, we have shown already in (I.10) that 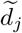 scales with (*np*λ_*j*_γ_*j*_)^1/2^ asymptotically, completing the proof.

## J Auxiliary results and proofs

The first result is a restatement of Proposition 2.1 of Vershynin (2012), which helps us establish the concentration of the estimated second moment matrix.

### Lemma J.1

(Probability bound of operator norm of a covariance matrix). *Let X*_1_,…, *X*_*n*_ *be i.i.d. k-dimensional random variables that satisfies Assumption 5.1 with a fixed constant D (also known as the sub-Gaussian norm). Then, for any δ,*

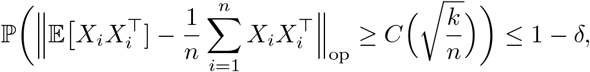

*for n sufficiently large enough, and where C is a constant that depends only D and δ.*

The following perturbation result can be derived using the perturbation results stated in Theorem 2 of Yu et al. (2014) with Hermatian dilation (see, e.g. Tropp (2012)) along with related notions of distance among orthonormal matrices stated in Lemma 1 of Cai et al. (2018), which helps us establish the concentration of the left singular vectors given concentration of the second moment matrices.

### Lemma J.2

(Davis-Kahan theorem for singular vectors). *For any two n* × *p symmetric matrices* Θ *and* 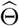 *where m* = min{*n*, *p*}, *let* σ_1_ > … > σ_*m*_ *and* 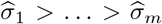 *denote the singular values of each matrix respectively, and U*_1_, …, *U*_*k*_ *and* 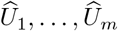 *denote their corresponding left singular vectors. Fix d* ∈ {1,…, *m*} *and let U* = [*U*_1_, …, *U*_*d*_] *and* 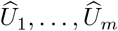 *denote the concatenated matrix of left singular vectors. Then, letting* 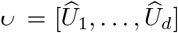 *denote the set of d* × *d unitary matrices,*

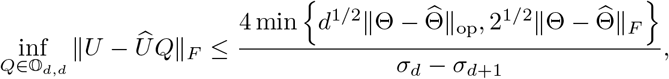

*where we use the convention that* σ_*m*+1_ = 0.

We note that when we apply the above lemma in our proofs throughout Appendix I, the uniqueness of the eigenvalues and the spacing of the eigenvalues assumed in Assumption 5.2 absolves us from needing to worry about the above unitary matrix *Q*. Instead, we simply need to worry “up to sign” as alluded to Proposition 5.2, where we need to possibly multiply th columns of 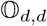 by ±1.

The next result on random sequences can be easily verified by direct calculation.

### Lemma J.3.

*For two positive deterministic sequences an and b*_*n*_, *consider a random sequence X*_*n*_ = *O*_*P*_(*a*_*n*_) *and a random positive sequence Y*_*n*_ = Ω_*P*_(*b*_*n*_). *If a*_*n*_ = *o*(*b*_*n*_), *then*

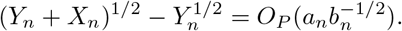

## Footnotes

1 We use the term “single-cell data” to refer to single-cell RNA-sequencing data specifically in the remainder of this article.

2 We call it a curved Gaussian distribution since this distribution is a curved exponential-family distribution.

3 We use “up to sign” similar to Fan et al. (2018), where each column of 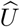 can be multiplied by ±1 since the SVD is not unique.

4 Their paper actually parameterizes the negative binomial distribution by its mean and inverse dispersion parameter, but is equivalent to the one we describe.

